# Temperature-dependent performance scales with maximum heat tolerance across ectotherms

**DOI:** 10.64898/2026.03.21.713427

**Authors:** Amanda S. Cicchino, Jessica Collier, Carling Bieg, Kaleigh Davis, Cameron. K. Ghalambor, Alison J. Robey, Jennifer Sunday, David A. Vasseur, Joey R. Bernhardt

**Affiliations:** Centre for Ecosystem Management, University of Guelph; Department of Integrative Biology, University of Guelph; Department of Biology, Case Western Reserve University; Department of Biology, Norwegian University of Science and Technology; Department of Ecology and Evolutionary Biology, Yale University; Department of Biology, McGill University

**Keywords:** thermal tolerance, CT_max_, thermal performance curve (TPC), ectotherm, climate warming

## Abstract

Temperature fundamentally shapes ectotherm biology, from continuous changes in sublethal performance to acute functional failure at upper thermal limits. Sublethal performance is commonly characterized using thermal performance curves (TPCs), which describe continuous changes in performance across temperatures, whereas acute heat tolerance is often quantified by the critical thermal maximum (CT_max_), the temperature marking organism-level functional failure. Whether these dimensions of thermal physiology vary independently or covary as components of a coordinated warm-end thermal phenotype remains unclear. Synthesizing data from more than 100 ectotherm species, we tested how CT_max_ scales with two key warm-end TPC traits: the thermal optimum (T_opt_) and the upper limit of performance (TPC_max_). Across diverse taxa, both T_opt_ and TPC_max_ scaled approximately 1:1 with CT_max_, and these relationships were broadly consistent across the biological, experimental, and environmental contexts represented in our dataset. Although both slopes were individually consistent with 1:1 scaling, when T_opt_ and TPC_max_ were compared directly, TPC_max_ increased more steeply with CT_max_ than did T_opt_, such that the declining portion of the TPC widened as CT_max_ increased. Species with greater acute heat tolerance therefore tended to retain positive performance across a broader range of supra-optimal temperatures before reaching the upper performance limit. Together, these relationships show that sublethal warm-end performance and acute heat tolerance covary systematically across species, supporting their interpretation as broadly coordinated components of the thermal phenotype rather than independent axes of thermal biology.

**Significance:** As climate warming pushes temperatures beyond the conditions species have previously experienced, predicting ectotherm vulnerability requires understanding how measures of thermal biology relate. Heat tolerance measures the maximum temperature organisms can survive, whereas temperature-dependent performance measures how they function across temperatures. Whether these measures capture related aspects of thermal biology has remained unclear. We synthesized thermal trait data from more than 100 ectotherm species to test how heat tolerance relates to temperature-dependent performance. We show that species with higher heat tolerance also reach peak performance and maintain function at higher temperatures, with these traits scaling approximately 1:1 across diverse contexts. This relationship links two major approaches to thermal biology and provides a quantitative basis for understanding ectotherm responses to warming.

## Introduction

For the vast majority of Earth’s species—the ectotherms—temperature fundamentally underlies biochemical, cellular, and physiological processes that shape fitness and population growth (1–4). As such, thermally sensitive traits have long been central to understanding the ecology and evolution of ectotherms (5, 6) and are increasingly used to predict species’ vulnerability to warming by relating trait variation to current and projected habitat temperature (e.g.,(7–11)). Two commonly used experimental approaches to quantify biological responses to temperature include estimating thermal performance curves (TPCs) and acute thermal tolerance limits (Figure 1). While TPCs describe performance across a range of sublethal temperatures (12–14), acute thermal tolerance limits represent the critical thresholds of biological failure (e.g., critical thermal maximum, CT_max_) (15). Despite their widespread use, it remains unclear how TPCs and thermal tolerance limits are related and whether the processes governing sublethal performance are fundamentally shared with or distinct from those underlying failure at thermal extremes (16). Developing an integrative understanding of biological responses to temperature is critical to understanding the ecology of ectotherms and for informing predictions of species performance in the context of global change.

**Figure 1.**
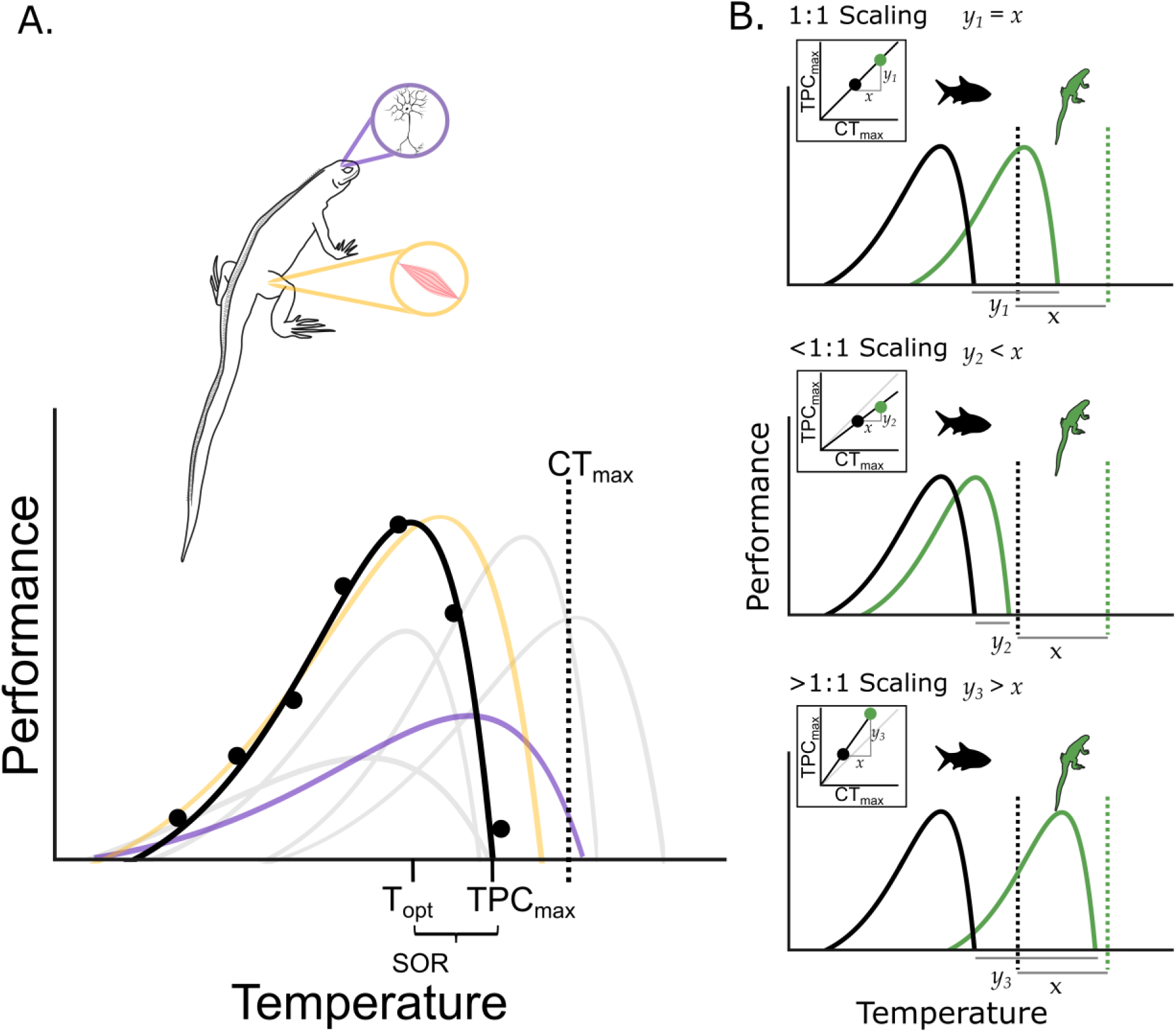
Conceptual integration and hypothesized scaling of thermal performance with thermal tolerance. (A) Conceptual illustration of how a thermal performance curve (TPC) for a measure of performance can emerge from the integration of multiple underlying temperature-dependent processes. For example, a TPC for lizard running speed (black) may reflect the effects of multiple contributing physiological processes (grey), including muscular (yellow) and neuronal (purple) performance. The TPC defines key traits, including the thermal optimum (T_opt_), the upper limit of performance (TPC_max_), and the supra-optimal range (SOR: the temperature interval between TPC_max_ and T_opt_). The critical thermal maximum (CT_max_; vertical dashed line) is an endpoint typically measured using acute heat-exposure assays and generally, though not always, occurs at temperatures greater than T_opt_ and TPC_max_. (B) Species (illustrated here as a black fish and green lizard) may vary in their TPCs and CT_max_ estimates (corresponding colors; CT_max_ shown as vertical dashed lines). Across species, CT_max_ and TPC traits—illustrated here using TPC_max_—may exhibit different scaling relationships. Because we predicted positive relationships between CT_max_ and TPC traits, the panels illustrate three possible forms of positive scaling; null and negative relationships are not illustrated. A 1:1 scaling relationship (top panel) indicates that increases in CT_max_ (*x*) are associated with equivalent increases in TPC_max_ (*y_1_*). A less than 1:1 scaling relationship (middle panel) indicates that increases in CT_max_ are associated with smaller increases in TPC_max_ (*y_2_*), such that TPC_max_ increases less rapidly than CT_max_ across species. Conversely, a greater than 1:1 scaling relationship (bottom panel) indicates that increases in CT_max_ are associated with larger increases in TPC_max_ (*y_3_*), suggesting that TPC_max_ increases more rapidly than CT_max_ across species. Grey lines in the middle and bottom panels indicate the 1:1 relationship for reference.

Temperature-dependent performance likely arises from the integration of several distinct processes, including kinetic effects on biochemical reaction rates (17, 18), the balance between cellular and molecular damage and repair (12, 19–21), and other constraints imposed by external factors such as oxygen supply and demand (22, 23) or dehydration (24, 25). Although a unifying framework has yet to be established, the near universality of TPC shape across ectotherms suggests common underlying constraints (26, 27) that may extend to heat tolerance limits (17, 18, 28). In this context, thermal tolerance limits may reflect the functional endpoint where constraints on sublethal performance ultimately propagate to whole-organismal failure. Within the damage-repair framework, for instance, performance declines as damage accumulation outpaces repair capacity; thermal tolerance limits would thus represent the point at which cumulative damage effects propagate to organismal failure (18, 28). Similarly, under the oxygen supply/demand framework, performance decreases when metabolic demands outpace the capacity to deliver oxygen (22, 23, 29); thermal tolerance limits may capture the endpoint when transport capacity is fully exhausted, triggering a transition to anaerobic respiration and eventual systemic collapse (30). Covariation among thermal traits may also arise through evolutionary processes that link trait responses to selection. Genetic correlations, pleiotropic effects, and correlated selection can generate coordinated evolutionary changes among traits, such that selection acting on one component of thermal performance may indirectly influence other components of the thermal niche (31–33). Thus, shared constraints and/or evolutionary correlations may produce predictable relationships among these traits across species, even if TPC traits and thermal tolerance limits represent distinct physiological processes.

Alternatively, thermal tolerance limits may reflect functional endpoints arising from processes that decouple them from sublethal performance. TPCs primarily span temperatures where organismal function is maintained, reflecting the balance of metabolism, resource allocation, and damage repair. In contrast, thermal tolerance limits may emerge from the abrupt breakdown of critical physiological systems. For example, at high temperatures, organisms may experience loss of neural function (34), disruption of cardiac electrical excitability (35), destabilization of membrane and ion channel regulation (36), or oxygen or hydric limitation (17, 34, 37), with CT_max_ representing the first system to fail. As a result, thermal tolerance limits may be better characterized by functional endpoints rather than as outcomes of continuous, rate-dependent processes like those captured in TPCs, and may operate on a different timescale than the steady-state dynamics shaping performance (i.e., “stressful range” versus “permissive range”) (18, 28). These differences are potentially reinforced by experimental design: thermal tolerance limits are often measured using rapid, acute exposure to thermal stress, whereas TPCs are quantified across a broader range of exposure durations. Longer exposure times can allow plasticity and bioenergetic constraints to influence performance (38), potentially integrating processes that are not captured by the rapid temperature exposure typical of acute tolerance assays. For example, acute heat stress and prolonged thermal acclimation can elicit distinct molecular responses (39–41), suggesting that the physiological processes engaged during acute thermal tolerance assays may not fully overlap with those shaping TPCs across longer timescales. Together, these differences suggest that temperature-dependent performance and thermal tolerance limits may represent partially or wholly independent traits, governed by distinct cellular, molecular, and genetic mechanisms (42, 43), and capable of evolving independently.

Resolving whether TPC traits and thermal tolerance limits represent related or independent dimensions of the thermal phenotype requires explicit comparative tests across taxa. Here, we synthesize a novel dataset of TPCs and thermal tolerance limits across ectotherms (Figure S1) to quantify the scaling relationships among these thermal traits. Given the increasing use of thermal traits in assessing ectotherm risk from warming (16), we focus our analyses on two upper-temperature TPC traits: the optimal temperature for performance (T_opt_) and the maximum temperature of performance (TPC_max_,(18). We compare these TPC-derived traits with critical thermal maximum (CT_max_), an acute measure of organismal heat tolerance (Figure 1). Our null expectation is that TPCs are unrelated to CT_max_ because they reflect functionally independent aspects of the thermal phenotype. Alternatively, if TPC traits and CT_max_ covary across species, we expect a positive relationship between these traits. Although negative relationships are possible, we predicted a positive relationship because species capable of maintaining performance at higher temperatures should also be expected to tolerate higher temperatures before reaching critical heat limits. The slope of this relationship describes how variation in TPC traits scales with variation in CT_max_. A slope of 1 indicates 1:1 scaling, such that a 1°C increase in CT_max_ is associated with a 1°C increase in the TPC trait, representing equal rates of change between the traits. Slopes below or above 1 indicate sublinear (< 1:1) or super-linear (> 1:1) scaling, respectively, such that TPC traits increase less or more than CT_max_ with each 1°C increase in CT_max_ across species.

A fundamental question is whether scaling relationships between TPC traits and vary across ecological and experimental contexts. Because these contexts are diverse, we focus on a subset of ecological and experimental contexts that we can test with our dataset. Environmental context, including climatic regime and habitat realm, may be associated with differences in the selective landscape shaping thermal trait evolution (5, 8, 11). Where exposure to temperatures near the heat tolerance limit is more frequent, selection may favor maintaining performance at high temperatures, potentially producing closer correspondence between TPC traits and CT_max_. We therefore expect scaling relationships to be closer to 1:1 in tropical than temperate taxa and in aquatic than terrestrial taxa (11). Similarly, experimental context, including the type of performance trait measured (13, 44), the biological scale at which performance is measured (13), and the duration of thermal exposure captured by an assay (38), may also influence relationships between TPC traits and CT_max_. Because CT_max_ is typically assessed as an acute, whole-organism endpoint, often based on neuromuscular responses, we expect locomotor TPCs, organismal TPCs, and acute TPCs to exhibit closer-to-1:1 scaling with CT_max_ than bioenergetic (e.g., metabolic rate), sub-organismal (e.g., liver enzyme reaction rate), and prolonged-duration (> 30 minute duration) TPCs, respectively.

TPC_max_ and T_opt_ together define the supra-optimal range (SOR; the difference between TPC_max_ and T_opt_), which captures the breadth of declining performance at temperatures above the optimum (Figure 1A). We therefore examine whether SOR varies with CT_max_, testing whether greater heat tolerance is associated with a wider temperature interval between T_opt_ and TPC_max_, or whether these traits shift together such that SOR remains unchanged. By examining the scaling of T_opt_ and TPC_max_ with CT_max_, and separately assessing how SOR varies with CT_max_, we test whether their patterns of covariation reflect an integrated warm-end thermal phenotype or instead independent aspects of ectotherm thermal physiology.

## Results

We initially compiled 3,349 thermal performance datasets (Figure S1). Following a filtering process which included resolving taxonomy and filtering for thermal performance datasets with at least five temperature treatments, including at least one treatment below and above the temperature at which the highest performance was observed, we retained 2,069 performance datasets representing 850 species. We matched 410 of these performance datasets, representing 200 species, with CT_max_ data for the same species, standardized to a common acclimation temperature of 15°C using taxon-summarized acclimation response ratios (see Methods; Figures S1, S2); we refer to these standardized values as CT_max_ hereafter. We then fit thermal performance curves (TPCs) to datasets of thermal performance and subjected the resulting curves to a quality control process which we designed to retain TPCs for which we could accurately and confidently estimate our thermal traits of interest (T_opt_ and/or TPC_max_) (Figure S1). This process yielded 204 TPCs (102 species) with high-confidence T_opt_ estimates, and 159 TPCs (79 species) with high-confidence TPC_max_ estimates; 136 TPCs (68 species) with high confidence estimates of both T_opt_ and TPC_max_, which were retained for SOR calculations. The final datasets spanned nine taxonomic classes and included locomotor and bioenergetic performance (SI Appendix, A.2. List 1), organismal and sub-organismal measurements, acute and prolonged exposures (SI Appendix, 2. List 2), aquatic and terrestrial taxa, and temperate and tropical climates (Figures S3, S4). Representation was uneven across these contexts, particularly for sub-organismal and tropical TPCs (Figure S3).

### TPC traits consistently scale with CT_max_ across ectotherms

Across ectotherms, T_opt_ and TPC_max_ increased with CT_max_ (Figure 2) and exhibited slopes consistent with 1:1 scaling, as the 95% confidence intervals overlapped the isometric slope expectation of 1 (T_opt_: 0.82 [0.61, 1.02]; TPC_max_: 0.95 [0.69, 1.20]; Tables S1 & S2, respectively). These scaling relationships were consistent across taxa when potential differences in slopes among taxonomic classes were evaluated for the three well-represented classes (Actinopterygii, Amphibia, and Reptilia) for which sufficient data were available to estimate class-by-CT_max_ interactions (Figure S5; Table S3).

**Figure 2.**
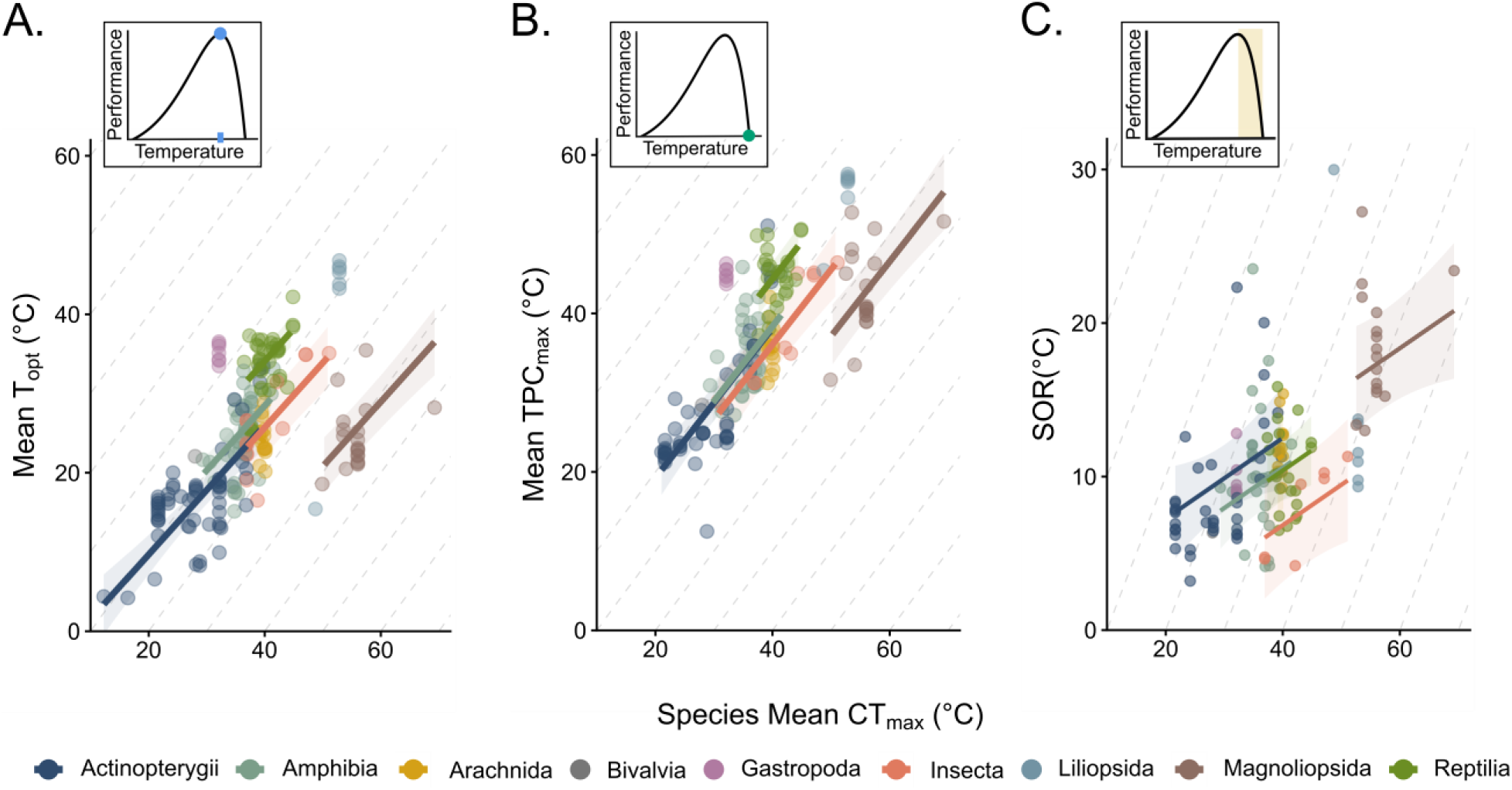
Relationships between thermal performance traits and heat tolerance. (A) T_opt_, (B) TPC_max_, and (C) the supra-optimal range (SOR) increased with CT_max_ across species. T_opt_ and TPC_max_ exhibited approximately 1:1 scaling relationships with CT_max_, while SOR scaled less than 1:1 with CT_max_. Points represent the mean of 1,000 bootstrapped estimates per TPC. CT_max_ values were standardized to a common acclimation temperature (ARR-standardized) of 15℃ (Eq. S2) and mean-centered in our analyses; axes here show back-transformed values on the original temperature scale. Solid lines show model-predicted relationships with 95% CIs (shaded). Grey diagonal dashed lines denote 1:1 scaling and are shown in all panels for visual consistency. For Topt and TPCmax, 1:1 scaling represents the hypothesized expectation under coordinated proportional shifts with CTmax. SOR is not interpreted relative to the 1:1 line; equal shifts in Topt and TPCmax would instead produce a constant SOR and a slope of zero. Regression lines are displayed only for taxonomic classes with sufficient representation to support interpretation of class-specific relationships. All represented classes were retained in the analyses. Sample sizes: T_opt_: N = 204, 102 species; TPC_max_: N = 159, 79 species; SOR: N = 136 TPCs, 68 species.

Trait variation among species was similar across thermal traits when evaluated using a shared set of species and TPCs with estimates available for both T_opt_ and TPC_max_ (136 TPCs representing 68 species; Figure S6). Within this dataset, variation was comparable between TPC_max_ (SD = 9.44 [7.35, 11.10]) and CT_max_ (SD = 9.74 [6.91, 11.6]). T_opt_ was slightly less variable (SD = 8.17 [6.27, 9.73]), although the 95% species-clustered bootstrap confidence intervals of all three traits overlapped (1,000 resamples). Thus, although they quantify distinct aspects of ectotherm thermal physiology, the three traits exhibited broadly similar magnitudes of variation within this shared dataset.

### Scaling relationships are consistent across biological and environmental contexts

Across performance response categories, we found no evidence that scaling relationships between CT_max_ and either thermal performance trait (TPC_max_ and T_opt_) differed between locomotor- and bioenergetic-based TPCs (T_opt_: CT_max_-by-response type interaction β = −0.09 [−0.29, 0.11], *p* = 0.37; TPC_max_: CT_max_-by-response type interaction β = 0.16 [−0.09, 0.41], *p* = 0.20; Figure 3; Table S4). In the subsequent additive models, locomotion-based TPCs exhibited higher TPC_max_ values at a given CT_max_ than bioenergetic TPCs (β = 5.40 [3.64, 7.16], *p* < 0.001), whereas T_opt_ did not differ between performance trait categories (Table S4).

**Figure 3.**
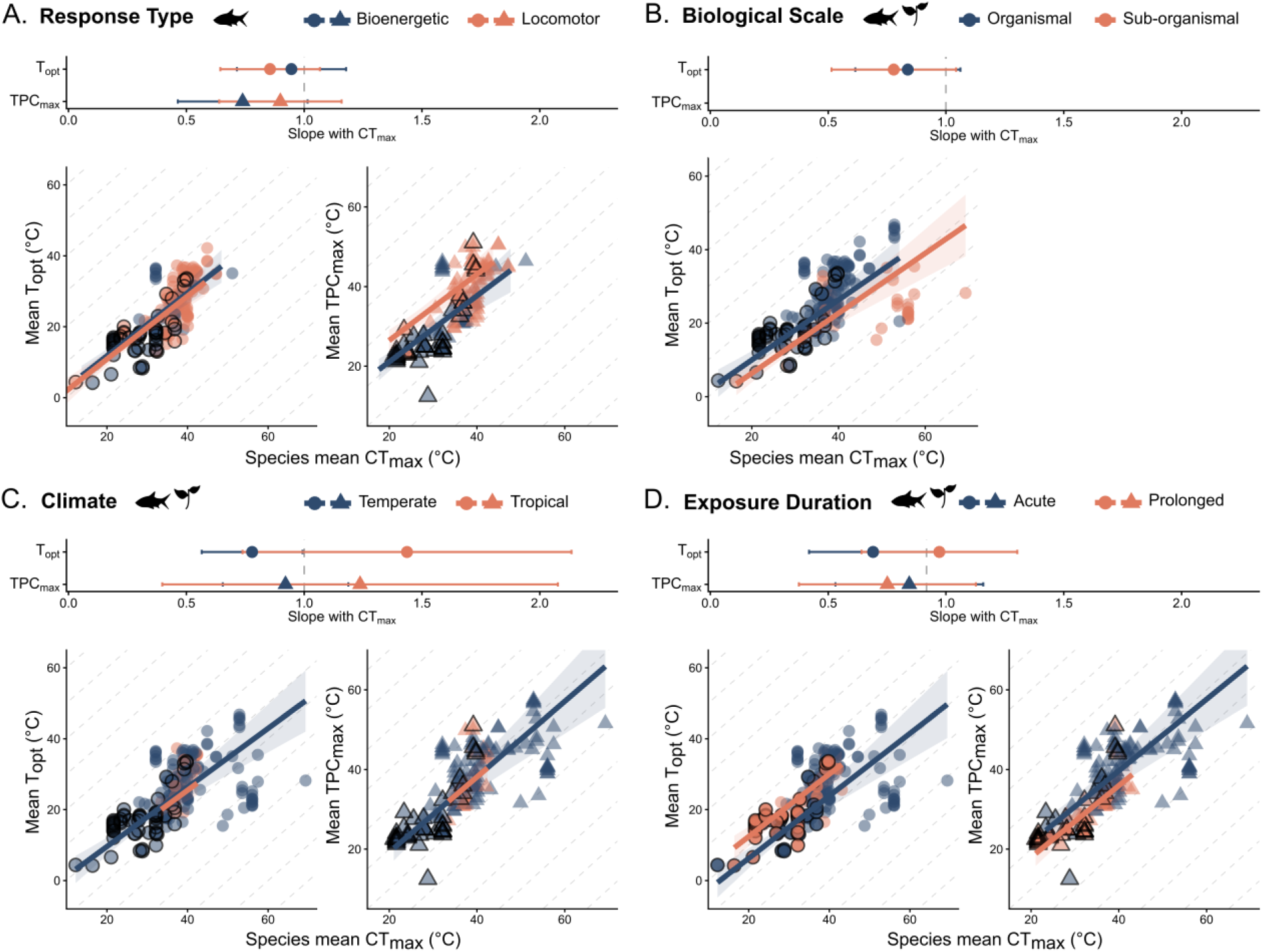
Context dependence of the scaling relationships between warm-end thermal performance curve traits and. CT_max_. Results are shown for (A) performance response type, (B) biological scale, (C) climatic zone, and (D) exposure duration. Top panels show category-specific slope coefficients and 95% confidence intervals describing the scaling of thermal optimum (T_opt_) and the upper limit of performance (TPC_max_) with CT_max_, estimated from models containing the context-by-CT_max_ interaction. None of the context-by-CT_max_ interactions were statistically supported, indicating no significant differences in slopes between context categories. Taxon icons indicate the data included in each analysis: paired fish and plant icons denote analyses including both animal and plant taxa, whereas a fish icon alone denotes an analysis restricted to animals (response type). Here, the fish icon represents animals generally rather than Actinopterygii specifically. The grey dashed background gridlines show the expectation under 1:1 scaling. Diagonal dashed lines denote 1:1 scaling and are shown in all panels for visual consistency. For Topt and TPCmax, 1:1 scaling represents the hypothesized expectation under coordinated proportional shifts with CTmax. SOR is not interpreted relative to the 1:1 line; equal shifts in Topt and TPCmax would instead produce a constant SOR and a slope of zero. Because estimates are derived from different hypothesis-specific models, they should be compared qualitatively rather than interpreted as direct statistical comparisons among hypotheses. Lower panels show relationships between species mean CT_max_ and mean T_opt_ or TPC_max_. Each point represents the T_opt_ or TPC_max_ estimate from an individual thermal performance curve, with black-outlined points indicating the reference taxonomic class for each model (Actinopterygii). Lines and shaded regions show predicted relationships and 95% confidence intervals from the best-supported hypothesis-specific model, holding taxonomic class at its reference level. Predictions are therefore shown from the interaction model when the context-by-CT_max_ interaction was supported and from the additive model otherwise. Diagonal dashed lines indicate 1:1 relationships. The figure presents contexts for which either the context-by-CTmax interaction or the additive context effect was supported. Environmental realm is omitted from the figure because neither effect was supported; complete environmental-realm results are reported in Table S7. TPC_max_ was not evaluated by biological scale because sample sizes were insufficient for that comparison. Climate zone results should be interpreted cautiously because tropical taxa were substantially underrepresented relative to temperate taxa. Sample sizes are given as TPCs, species: T_opt_—locomotor: 104, 62; bioenergetic: 64, 29; organismal: 174, 87; sub-organismal: 28, 14; aquatic: 97, 39; terrestrial: 107, 63; temperate: 175, 81; tropical: 29, 21; acute: 143, 76; prolonged: 61, 26. TPC_max_—locomotor: 75, 26; bioenergetic: 53, 41; aquatic: 78, 29; terrestrial: 81, 50; temperate: 137, 62; tropical: 22, 18; acute: 108, 55; prolonged: 51, 24.

We found no evidence that the T_opt_–CT_max_ relationship differed among biological scales (i.e. between organismal and sub-organismal scales; CT_max_-by-biological scale interaction β= −0.06 [−0.32, 0.20], *p* = 0.66; Figure 3; Table S5). Sub-organismal observations were too sparse to robustly evaluate the corresponding TPC_max_ relationship (N = 21), and 19 of 21 observations were derived from plants (Figure S3). In the subsequent additive model, sub-organismal TPCs exhibited lower predicted T_opt_ values at a given CT_max_ than organismal TPCs (β = –3.61 [–4.15, – 3.07], *p* < 0.001; Table S5).

T_opt_–CT_max_ scaling did not differ between acute- and prolonged-exposure assays (duration-by-CT_max_ interaction β = 0.26 [−0.06, 0.58], *p* = 0.12; Figure 3; Table S6), nor did TPC_max_–CT_max_ scaling (duration-by-CT_max_ interaction β = −0.09 [−0.49, 0.31], *p* = 0.67; Figure 3; Table S6). Exposure duration was, however, associated with intercept differences for both traits. At a given CT_max_, prolonged-exposure assays had a higher predicted T_opt_ (subsequent additive model β = 6.15 [5.18, 7.11], *p* < 0.001) but a lower predicted TPC_max_ (subsequent additive model β = −3.72 [−6.38, −1.06], *p* = 0.006) than acute-exposure assays (Figure 3; Table S6).

Scaling relationships between thermal traits and CT_max_ were consistent in terrestrial and aquatic environments. We found no evidence that environmental realm modified either the T_opt_–CT_max_ relationship (CT_max_-by-realm interaction: β = −0.25 [−0.67, 0.18], *p* = 0.25) or the TPC_max_–CT_max_ relationship (CT_max_-by-realm interaction: β = −0.35 [−0.86, 0.15], *p* = 0.17; Table S7). Subsequent additive models also provided no evidence that environmental realm was associated with differences in either trait at a given CT_max_ (Table S7).

We found no evidence that CT_max_–thermal performance scaling relationships varied across climatic realms (tropical vs. temperate). Climate did not modify the TPC_max_–CT_max_ relationship (CT_max_-by-climate interaction: β = 0.32 [−0.55, 1.19], *p* = 0.48; Figure 3; Table S8) or the T_opt_– CT_max_ relationship (CT_max_-by-climate interaction: β = 0.66 [−0.07, 1.39], *p* = 0.08; Figure 3; Table S8). In the subsequent additive models, tropical taxa exhibited lower T_opt_ values than temperate taxa at a given CT_max_ (β = −0.47 [−0.90, −0.04], *p* = 0.03), whereas TPC_max_ values did not differ between climatic zones (Table S8). However, this difference in T_opt_ should be interpreted cautiously given the uneven representation of climatic realms in our dataset (temperate: 175 TPCs, 81 species; tropical: 29 TPCs, 21 species).

### Supra-optimal range increases sub-linearly with CT_max_, reflecting steeper scaling of TPC_max_ relative to T_opt_

Because the supra-optimal range (SOR) is calculated as the difference between TPC_max_ and T_opt_, its relationship with CT_max_ is expected to reflect any difference in how these two traits scale with CT_max_. We calculated SOR for a subset of TPCs from which we were able to confidently estimate both T_opt_ and TPC_max_ (see Figure S1), such that both T_opt_ and TPC_max_ were estimated from the same performance curve. SOR scaled positively, but sub-linearly with CT_max_ (slope = 0.26 [0.01, 0.52], p = 0.04; Table S9; Figure 2C). To evaluate whether this pattern reflected differences in the scaling of T_opt_ and TPC_max_, we directly compared the two traits within this dataset. TPC_max_ scaled with CT_max_ at a significantly steeper rate than T_opt_ (CT_max_-by-trait type interaction: β = 0.31[0.30, 0.32], *p* < 0.001; TPC_max_ slope = 0.99 [0.74, 1.24] vs. T_opt_ slope = 0.68 [0.43, 0.93]; Table S10; Figure S7), a difference strongly supported by a likelihood ratio test comparing models with and without this interaction (χ² = 8936.86, p < 0.001). Consistent with this difference, the absolute gap between these two traits widened with CT_max_ but increased proportionally more slowly than CT_max_, producing the observed sub-linear SOR–CT_max_ relationship. Together, these results indicate that although both TPC traits increase with CT_max_, high-CT_max_ species exhibit supra-optimal performance declines that begin at increasingly cooler temperatures relative to their CT_max_, resulting in broader supra-optimal ranges at higher CT_max_.

## Discussion

Whether performance across sublethal temperatures and acute functional failure represent independent or coordinated dimensions of ectotherm thermal physiology has remained unclear. Across datasets spanning more than 100 ectotherm species, we found approximately 1:1 scaling between warm-end performance traits and acute heat tolerance. Specifically, species with higher CT_max_ tended to exhibit correspondingly higher thermal optima (T_opt_) and the upper limit of performance (TPC_max_; Figure 2). These relationships were observed across diverse taxa and were broadly consistent across diverse taxa, response types, biological scales, thermal exposure durations, environmental realms, and climatic zones (Figure 3). Within this overall scaling pattern, however, TPC_max_ increased more steeply with CT_max_ than did T_opt_, producing a positive relationship between the supra-optimal range (SOR) and CT_max_ (Figure 2). Species with higher CT_max_ therefore exhibited broader SORs, indicating that the two warm-end traits (T_opt_, TPC_max_) did not shift uniformly and that the declining portion of the TPC widened with increasing heat tolerance. Together, the scaling of T_opt_ and TPC_max_ and the systematic increase in their separation with CT_max_ provide evidence that sublethal thermal performance and acute heat tolerance are broadly coordinated components of the thermal phenotype rather than independent axes of thermal biology.

Although T_opt_ and TPC_max_ describe distinct components of thermal performance that may experience different selective pressures (6, 31, 45, 46), both T_opt_ and TPC_max_ exhibited remarkably similar 1:1 scaling relationships with CT_max_. The similarity in scaling suggests that variation in heat tolerance is accompanied by coordinated shifts across multiple warm-end components of thermal performance. Our result aligns with previous evidence that components of the TPC are positively evolutionarily correlated (31, 47), while extending this understanding in an important way: previous evidence for evolutionary covariation among T_opt_ and TPC_max_ has been based on traits estimated from the same performance curve, whereas our results show similar covariation among independently estimated TPC traits and acute heat tolerance across species. Previous work suggests that heat tolerance limits may be more evolutionarily constrained than other components of the thermal niche (8, 48–51) but see (52). If CT_max_ were uniquely constrained relative to other warm-end thermal traits, we might expect it to exhibit substantially less realized variation. Instead, variation was similar among CT_max_, T_opt_, and TPC_max_ within the shared SOR dataset, with only slightly lower variation in T_opt_. This pattern could indicate that CT_max_ is not uniquely constrained relative to other warm-end thermal traits, or that evolutionary constraint extends more broadly across the warm-end thermal phenotype, although the physiological and evolutionary processes underlying this coordination remain unresolved. Together, the similar scaling relationships and comparable magnitudes of variation suggest that T_opt_, TPC_max_, and CT_max_ represent coordinated components of the thermal phenotype.

The scaling relationships we observed were generally consistent across the experimental and environmental contexts tested in our dataset. We found no evidence that response type, biological scale, exposure duration, environmental realm, or climatic zone altered the slopes of the relationships between CT_max_ and either T_opt_ or TPC_max_. Where context effects occurred, they primarily involved differences in intercepts, altering expected thermal trait values at a common CT_max_ in context- and trait-specific ways. For example, locomotor responses exhibited higher predicted TPC_max_ than bioenergetic responses despite exhibiting similar thermal optima, suggesting that locomotor performance may be maintained at higher temperatures, or decline more slowly, than many bioenergetic processes in species with similar heat tolerance. Similarly, prolonged-exposure assays were associated with higher predicted T_opt_ but lower predicted TPC_max_ than acute-exposure assays at a common CT_max_. Although the mechanisms underlying these contrasting associations remain unresolved, this pattern raises the possibility that sustained thermal exposure affects warm-end performance traits differently, potentially through trait-specific plastic responses to prolonged exposure (38). Thus, experimental and environmental contexts were associated with differences in the expected values of warm-end performance traits but did not alter how those traits scaled with heat tolerance. The approximately 1:1 scaling relationships therefore remained broadly robust across the contexts tested here.

Despite deep evolutionary divergence in ectotherm thermal physiology (8, 48–50, 53), we found no evidence that TPC–CT_max_ scaling differed among the best-represented classes in our dataset (Actinopterygii, Amphibia, and Reptilia). This pattern suggests that coordinated variation between heat tolerance and thermal performance may represent a general feature of ectotherm evolution rather than a lineage-specific phenomenon. Importantly, this similarity in scaling occurred despite differences among classes in the absolute values of thermal traits. Class effects on intercepts indicated differences in the expected thermal traits value at a common CT_max_, whereas class-by-CT_max_ interactions provided no evidence that scaling slopes differed. Thus, taxonomic divergence may influence the magnitude of thermal traits more strongly than their covariation with heat tolerance. However, only three vertebrate classes were sufficiently represented to evaluate lineage-specific differences; our results thus support broad conservation of scaling among the represented lineages but do not establish invariance broadly across ectotherms.

The positive relationship between the supra-optimal range (SOR) and CT_max_ shows that the warm end of TPCs did not simply shift uniformly along the temperature axis as acute heat tolerance increased. Instead, TPC_max_ increased more steeply with CT_max_ than did T_opt_, such that the temperature span of the declining portion of the TPC (SOR) widened with increasing CT_max_. Consequently, species with higher acute thermal limits tended to retain positive performance across a wider range of supra-optimal temperatures before reaching the upper limit of performance. Together, these relationships support a coordinated warm-end thermal phenotype in which species with higher CT_max_ tended to exhibit higher T_opt_ and TPC_max_ and broader SORs. Variation in SOR may be particularly consequential under warming. As warming shifts experienced body temperatures beyond T_opt_, organisms will increasingly operate on the declining portion of the TPC, and the thermal interval over which they retain positive performance may become increasingly important for persistence (7, 10, 54, 55). Our results suggest that species with higher CT_max_ tend to have a larger post-optimum thermal buffer across this supra-optimal region before performance reaches zero.

Considerable variation in T_opt_ and TPC_max_ remained after accounting for CT_max_ and taxonomic class, including both among-species variation and residual heterogeneity among TPCs. Thus, CT_max_ captures an important axis of variation in thermal performance but does not account for the full diversity of thermal phenotypes. Numerous sources of variation that were not examined here may contribute to this remaining variation and warrant further investigation, including variation among populations and individuals, TPC plasticity, and thermal environments (3, 56–58). Nevertheless, the consistency of the scaling relationships across our comparatively coarse-grained measures of biological, experimental, and environmental context is notable. The relationships emerged despite substantial heterogeneity among the underlying studies, highlighting the robustness of the association between heat tolerance and warm-end thermal performance.

Our results also provide additional context for what CT_max_ can reveal about thermal performance and its potential utility for comparative assessments of vulnerability to warming (16, 59–61). Although CT_max_ captures a failure endpoint that may rarely be experienced in nature (49, 62), its approximately 1:1 scaling with TPC traits demonstrates that it also captures meaningful variation in thermal traits relevant to temperatures frequently experienced in nature and directly tied to fitness. Consequently, in the absence of TPCs, CT_max_ may provide a useful *comparative* indicator of differences among species in their potential to maintain performance as temperatures increase. However, CT_max_ alone cannot capture the nonlinear and cumulative effects of temperature on performance that occur before acute performance failure. While the widespread availability and relative ease of measuring CT_max_ make it a practical starting point for large-scale assessments of thermal risk across diverse taxa, predicting responses to chronic and sublethal warming still requires TPCs to quantify how performance changes across the full thermal environment.

Ultimately, we demonstrate a consistent quantitative relationship between acute heat tolerance and multiple components of thermal performance across diverse ectotherms. The approximately 1:1 scaling of T_opt_ and TPC_max_ with CT_max_ across ectotherm species, together with the consistency of these patterns across the various contexts we tested, indicates that heat tolerance and warm-end thermal performance are broadly coordinated components of the thermal phenotype. Thus, acute heat tolerance provides meaningful insight about the broader thermal phenotype rather than representing an isolated endpoint of physiological failure. These scaling relationships provide a basis for integrating widely available CT_max_ data with temperature-dependent performance in models of organismal and population responses to climate warming (20, 21, 63–65). Such integration could allow CT_max_ to inform expectations about the temperatures associated with peak and upper performance where TPCs are unavailable, while TPCs remain necessary for resolving how performance changes across the thermal environment and under chronic or sublethal warming. Moving forward, determining how phylogenetic history, genetic variation, plasticity, and environmental exposure generate variation around these broadly conserved scaling relationships will be important for understanding the limits and generality of this coordination.

## Materials and Methods

To test our hypotheses about the relationships between thermal performance curves (TPC) and maximum temperature tolerance (CT_max_), we assembled a global dataset of TPCs and CT_max_ from terrestrial, marine, and freshwater ectotherms. We compiled thermal performance data used to fit TPCs, from which we derived the temperature of peak performance (thermal optimum, T_opt_), the maximum temperature of positive performance (TPC_max_), and the supra-optimal range (SOR; the range of temperatures between T_opt_ and TPC_max_). We then used hierarchical mixed-effects models to quantify the strength and proportionality of relationships between TPC traits and CT_max_, and to test for variation in these relationships across the biological response category captured in the TPC, the biological scale of the TPC, and TPC assay duration.

### Data compilation and trait classification

We compiled data to fit TPCs from published studies and existing datasets (66, 67), retaining studies that had performance values across at least five temperatures (Figure S1). We standardized performance values to common units where necessary (e.g., durations were converted to rates) to ensure comparability across studies. We compiled CT_max_ values from published thermal-limit databases and literature syntheses (9, 68–70), which compile experimentally measured thermal limits from published studies across diverse ectothermic taxa, as well as targeted searches for species with TPC data (SI Appendix, A.1.i). These sources use a range of endpoints to characterize maximum thermal limits. We included all endpoints of maximum thermal limits (e.g., onset of muscle spasms, loss of righting response, 50% tissue mortality) and refer to them collectively as CT_max_ (see SI Appendix, A.1.iv.d. *Sensitivity analysis of CTmax endpoints*).

To account for the effects of acclimation temperature on CT_max_ (71–73), we compiled CT_max_ acclimation response ratios (ARR), which quantify the change in CT_max_ per degree increase in acclimation temperature (SI Appendix, Equation S1), from published studies (70, 74–77). We standardized CT_max_ values to a common acclimation temperature (SI Appendix, Equation S2) using taxon-summarised ARRs. Because standardized and unstandardized CT_max_ estimates were highly correlated across taxonomic levels (Table S11), we used phylum-level mean ARRs for all estimates. We present our results using CT_max_ values ARR-standardized to 15°C, a pragmatic choice as it is the mean global surface temperature (78), although our conclusions were robust to the choice of standardization temperature (Figure S2). While we evaluated sensitivity to endpoint definition within the animal Topt dataset (SI Appendix, A.1.iv.d), other experimental influences on CTmax, including heating rate (79), could not be standardized because the relevant protocols were not reported consistently or comparably across taxa and studies.

To support our hypothesis testing, we classified performance assays by response type (bioenergetic, locomotion, life history; SI Appendix, A.2, List ii; Figure S3) and level of biological organization (sub-organismal, organismal, population). We classified thermal performance assays by exposure duration to capture the potential for plasticity and bioenergetic constraints to influence performance during the experiment. Assays lasting ≥ 30 minutes—long enough for physiological responses such as the upregulation of heat-shock proteins (80)—were classified as “prolonged” in duration, while assays < 30 minutes were classified as “acute”. We also categorized species based on environmental realm (aquatic or terrestrial) and climatic region. Climatic region was assigned using the geographic location of the sampled population when available; otherwise, we used the species’ known geographic distribution. Species were classified as tropical if the sampling location (or species distribution) occurred between 23.5° N and 23.5° S, and as temperate otherwise.

### TPC fitting and trait estimation

We fit thermal performance curves (TPCs) to performance data using non-linear least squares (nlsLM, *minpack.lm* in R) and the Norberg model (81) as modified by (82) (referred to as the Norberg-Thomas model) (Equation 1). We modeled temperature-dependent performance as:

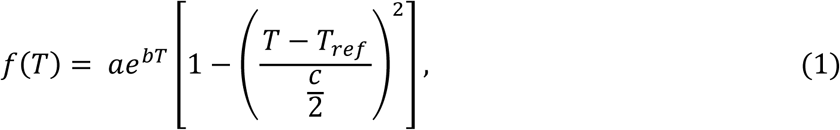

where performance, *f*, varies as a function of temperature, *T* (°C), *a* acts as a scaling parameter, *b* determines the slope of the exponential rise and the direction of the curve skew, *c* captures the breadth of positive performance, and *T_ref_* is the temperature at which the declining quadratic function peaks. We chose the Norberg-Thomas model because it accommodates both left- and right-skewed curves, requires only four parameters, is suitable for datasets with ≥ 5 temperatures, and provides reliable estimates of thermal traits under varying levels of data quality (see SI Appendix, A.1.iii for sensitivity analyses and comparisons of the Norberg-Thomas model to other thermal performance curve models).

We derived TPC traits from each fitted TPC’s parameter sets. We calculated T_opt_ via numerical optimization of the fitted curve. The upper thermal limit of performance (TPC_max_) has historically been quantified in several ways, including direct experimental measurement, assuming it is equivalent to CT_max_, or deriving it from a fitted thermal performance curve. Here, we estimated TPC_max_ from the fitted TPC model, ensuring that all TPC traits were derived using a common analytical framework. Specifically, TPC_max_ was calculated directly from model parameters (TPC_max_ = *T_ref_* + *c*/2; Equation 1), and SOR as the difference between TPC_max_ and T_opt_. To quantify uncertainty in TPC-derived traits, we used a residual bootstrap with the Boot function in the *car* package in R (83) with 10,000 iterations per TPC, and calculated T_opt_, TPC_max_, SOR from each refitted curve. We applied quality control filters to exclude poor fits (see Figure S1 for data pipeline), including those with high uncertainty (95% confidence intervals estimated from bootstraps) in trait estimates or parameter estimates near model bounds (SI Appendix, A.1.ii). We assessed data quality independently for each TPC-derived trait to ensure that low-confidence estimates for one trait did not lead to the unnecessary exclusion of robust data for another trait.

### Statistical Analyses

We conducted all analyses in R v4.5.1 (84). We tested for relationships between TPC traits (T_opt_ and TPC_max_) and CT_max_ using multilevel meta-regressions fitted using restricted maximum likelihood using the *rma.mv* function in the *metafor* package (85). This approach incorporates known sampling variances, accommodates unbalanced data, and allows flexible random-effects structures. For each model, we used the mean of 1,000 randomly-selected bootstrap-derived TPC trait estimates as the response and their variance as the sampling variance. We included species as a random intercept to account for non-independence among TPCs from the same species and taxonomic Class as a fixed effect to account for broad taxonomic differences among taxa. We selected Class as the taxonomic level for the fixed effect following model comparison via Akaike Information Criterion (AIC, (86), Table S12). We assessed model assumptions using residual diagnostics and found no substantial departures from linearity or other model assumptions.

As an initial step, we fitted baseline meta-regression models with centered CT_max_ as the sole fixed effect to quantify the overall direction and strength of trait associations. Because CT_max_ and the TPC traits measured here are expressed on the same temperature (°C) scale, we evaluated their relationships on the raw °C scale, which provides direct biological interpretation by estimating the expected change in TPC traits associated with a 1°C change in CT_max_ (see SI Appendix, A.1.iv for exploration of alternative formulations). These baseline models established the direction, strength, and scaling of TPC–CT_max_ relationships in the absence of additional covariates.

We subsequently tested hypotheses predicting that TPC–CT_max_ relationships vary according to response type (locomotor or bioenergetic processes), biological scale (organismal or sub-organismal), exposure duration (acute or prolonged), environmental realm (aquatic versus terrestrial), or climatic region (temperate versus tropical). Analyses of response type were restricted to TPCs derived from locomotor and bioenergetic traits; TPCs derived from life-history traits (n = 7) traits were excluded from this comparison because they were insufficiently represented and were not directly comparable to these response categories. Similarly, analyses of biological scale were restricted to organismal and sub-organismal measurements because population-level TPCs (n = 2) were too sparse for reliable inference. TPCs excluded from either moderator analysis were retained in all other applicable analyses. For each hypothesis, we fitted a meta-regression model including the focal predictor as a main effect and in interaction with CT_max_ to test whether the context modified the CT_max_–trait slope. Where there was no evidence of an interaction, we subsequently fitted an additive model including the focal predictor to test whether context was associated with differences in trait values at a given CT_max_.

Finally, because SOR is calculated as the difference between TPC_max_ and T_opt_, we separately tested its relationship with CT_max_ using TPCs for which both TPC_max_ and T_opt_ were estimated from the same performance curve. We also directly compared TPC_max_ and T_opt_ scaling with CT_max_ using a model including trait type and its interaction with CT_max_ to determine whether the two traits differed in their scaling slopes.

We assessed model support using AIC by comparing each hypothesis-specific model with a null model including Class as a fixed effect and species as a random intercept. We evaluated the overall significance of each hypothesis-specific model using omnibus tests of moderators (QM) and assessed individual predictor and interaction effects using Wald-type tests, and 95% confidence intervals. We also calculated a *pseudo*-R^2^ as the proportional reduction in the species-level random-effect variance relative to the null model and assessed residual heterogeneity using the QE statistic.

To compare variation among thermal traits, we calculated the observed standard deviation of T_opt_, TPC_max_, and CT_max_ across the shared (SOR) dataset. We quantified uncertainty using 1,000 cluster bootstrap resamples, sampling species with replacement and retaining all TPCs associated with each selected species. We calculated 95% percentile confidence intervals from the 2.5th and 97.5th percentiles of each bootstrap distribution.

## Acknowledgements

We thank Helen Vanos for logistical support.

## Funding

This study was the result of a working group supported by the Canadian Institute for Ecology and Evolution (ASC & JRB) and the University of Guelph Centre for Ecosystem Management (ASC & JRB). This work was also supported by the Natural Sciences and Engineering Research Council of Canada (NSERC-PDF to ASC; NSERC Discovery Grant to JRB).

## Data Availability

All data and code required to reproduce the analyses will be available on Zenodo.

## Author Contributions

Conceptualization, methodology, Writing - Review & Editing: All authors

Investigation: ASC, JC, JRB

Validation: ASC, AR, JRB

Data Curation Visualization, Writing - Original Draft: ASC

Project administration, Funding acquisition, Formal analysis: ASC, JRB

Supervision: JRB

## Notes

### Competing Interest Statement

The authors have declared no competing interest.

### Summary of Updates

The analytical approach has been updated, yielding additional results, including analyses examining variation across climatic realms and between terrestrial and aquatic habitats.

