## Supporting Information for "Temperature-dependent performance scales with maximum heat tolerance across ectotherms"

Affiliations:

**This PDF file includes:**

**Part A: Supplementary Text**

**Part B: Supplementary Tables**

**Part C: Supplementary Figures**

**Part D: TPC Fits and trait estimates**

**References**

### Part A: Supplementary Text

#### 1. Supplementary Methods

**i. Data sources and compilation methods**

Our data compilation includes i) measures of thermal performance, ii) upper thermal tolerance limits, and iii) plasticity in upper thermal tolerance limits for ectotherms across the tree of life.

*Thermal performance curves (TPCs)*

We performed an extensive literature and database search to compile TPC datasets across ectotherms. We first performed a literature search, querying the Web of Science Core Collection using search terms including "temperature OR thermal OR thermos*" and a list of performance traits commonly measured for TPCs of animal ectotherms (SI List 1). This search yielded 16,795 published studies, which we imported to Covidence (1) for title and abstract screening. We excluded studies where temperature was not directly experimentally manipulated and those where traits were measured across fewer than three temperatures. Following full-text screening, we kept studies for data extraction if they had at least four temperature treatments. We extracted metadata and performance values for traits from text, tables, or from figures using WebPlotDigitizer v5.2 (2).

Given that the objective of this study was to evaluate the relationship between TPC traits ($T_{opt}$ and $TPC_{max}$) and $CT_{max}$, after the screening of 386 studies yielded 26 studies for data extraction, we switched to targeted searches for species with known $CT_{max}$ (see below). We performed searches in Google Scholar using the species' name as a search term along with "AND (therm* OR temp*) performance curve". We screened search results using the same exclusion criteria as above and included studies that fit our criteria but were not explicitly searched for. We attempted to maintain taxonomic representation in our dataset by searching for species in each family until a minimum of 25% of species in our $CT_{max}$ list were searched. Following data extraction, all data were converted to consistent measurement units if needed (e.g., durations were converted to rates). We combined the results of our TPC searches with data from published datasets of ectotherm thermal performance ((3) $N_{TPC}$=2739; (4) $N_{TPC}$=194). Finally, we retained TPC data with a minimum of five temperature points, including one temperature point on either side of the maximum observed trait value, to allow for TPC model fitting.

For each study, we also collected or categorized metadata to facilitate comparative analyses. We broadly categorized each trait into a physiological response category representing bioenergetic processes (e.g., consumption rate, metabolic rate), locomotion (e.g., sprinting speed), and life history (e.g., fecundity). We also categorized TPCs by assay duration to capture the potential for active plasticity (*sensu* (5, 6)) and bioenergetic constraints to influence performance. Performance measured over timescales where plasticity or bioenergetics could affect outcomes

(≥30 minutes; e.g., (7)) was classified as "prolonged" duration, while performance measured at shorter timescales (<30 minutes), where these processes are unlikely to occur, was classified as "acute" (*SI Appendix, A.1, List iii*). Lastly, we included the biological scale of the TPC response: population, organismal, and sub-organismal. We categorized responses measured at the level of a single organ or tissue (e.g., liver metabolism) as sub-organismal, whereas responses that integrate sub-organismal processes but are quantified at the whole-organism level (e.g., whole-organism respiration) were categorized as organismal. We also categorized species based on environmental realm (aquatic or terrestrial) and climatic region (tropical or temperate). Climatic region was assigned using the geographic location of the sampled population when available; otherwise, we used the species' known geographic distribution. Species were classified as tropical if the sampling location (or species distribution) occurred between 23.5° N and 23.5° S, and as temperate otherwise.

*Maximum heat tolerance ($CT_{max}$)*

We compiled upper thermal limit data by combining existing thermal limit databases and literature-synthesized datasets from ectotherms (8–11), retaining all measures of heat tolerance. For animals, these measures included (e.g., dynamic $CT_{max}$, lethal dose 50%). For species with available TPC data but no corresponding $CT_{max}$ estimates, we conducted targeted literature searches for $CT_{max}$ using the species name in combination with the terms "$CT_{max}$" or "upper thermal limit." We also extracted $CT_{max}$ values when they were reported in the same studies that provided TPC data. For all $CT_{max}$ data we collected, we noted acclimation temperatures for the $CT_{max}$ experiments when they were available. When using existing $CT_{max}$ compilations, we filled in missing acclimation temperatures by referring to the original studies from which $CT_{max}$ data were compiled. When acclimation treatments were diurnally fluctuating, we used the daytime temperature as the acclimation temperature. If $CT_{max}$ data were collected from study organisms in the field or recently collected from the field, we used the habitat temperature reported in the original study as the acclimation temperature.

*Plasticity in heat tolerance*

To account for the effects of acclimation temperature on $CT_{max}$ (12–15), we compiled data on plasticity in $CT_{max}$. We estimated plasticity in $CT_{max}$ using acclimation response ratios (ARR), which measure the magnitude of change in $CT_{max}$ per degree increase in acclimation/holding temperature (Equation S1). We compiled species-specific $CT_{max}$ ARRs from existing literature-synthesized datasets ((10) N=179) ((16) N=320) ((17) N=95) ((18) N=21) ((19) N=24). For studies reporting $CT_{max}$ estimates from multiple acclimation temperatures, we calculated ARR using estimates from the minimum and maximum acclimation temperatures (Equation S1) as:

$$ARR = \frac{CTmax_{\max acclimation\ temperature} - CTmax_{\min acclimation\ temperature}}{\max acclimation\ temperature - \min acclimation\ temperature}. \qquad (S1)$$

We calculated mean ARR at multiple taxonomic levels (kingdom, phylum, class, order, family) and used these means to standardize $CT_{max}$ values to a common acclimation temperature. We calculated $CT_{max}$ at the standardized temperature, $n$, ($CT_{maxn}$) as a function of $CT_{max}$ at the experimental acclimation temperature ($CT_{maxexp}$) and the phylum-average ARR ($ARR_{phylum}$) (Equation S2) as:

$$CTmax_n = CTmax_{exp} + [ARR_{phylum}(acclimation\ temperature_n - acclimation\ temperature_{exp})]. (S2)$$

Having found that $CT_{max}$ values standardized using ARR means from different taxonomic levels were highly correlated with one another and with unstandardized $CT_{max}$ values (Table S11), we used phylum-level mean ARRs for all subsequent analyses. We present our results using $CT_{max}$ values standardized to 15°C, the mean global surface temperature (NOAA 2024), however, to assess sensitivity to the choice of standardization temperature, we repeated our TPC trait~$CT_{max}$ multilevel meta-regression models using $CT_{max}$ standardized from 0–50°C in 1°C increments, which did not qualitatively alter the results (Figure S1). Importantly, the standardization does not assume that these organisms were acclimated at 15°C; rather, it provides a common reference point for comparing $CT_{max}$ estimates obtained under different acclimation conditions.

*Merging compiled datasets by taxonomy*

For all species with TPC, $CT_{max}$, or $CT_{max}$ plasticity data, we standardized taxonomy to enable dataset merging. Species names were matched to the GBIF backbone (via *rgbif* in R), and unresolved names were sequentially queried against NCBI, Catalogue of Life, and the Open Tree of Life using the Global Names Architecture. Remaining mismatches were rechecked in GBIF with synonym matching. For each species, we recorded the original and accepted names, full taxonomy, and a unique identifier used for merging.

From our merged dataset, we excluded species that were missing either $CT_{max}$ or TPC data. To ensure reliable TPC model fitting, we also excluded species with TPC data that had fewer than five temperature treatments or without temperature treatments on either side of the observed maximum performance (i.e., the temperature of the maximum response trait value).

**ii. TPC fitting and quality control**

We fit thermal performance curves (TPCs) using the Norberg-Thomas model (Equation S3) to performance data using non-linear least squares regression using the nlsLM function from the *minpack.lm* package in R. Because source studies varied in experimental design and data structure—ranging from repeated measures on individuals to studies reporting only summary statistics (e.g., mean performance per temperature treatment)—we fit curves to the highest resolution data available from each source. For studies reporting individual-level observations, curves were fit directly to those observations; for studies reporting only treatment means, curves were fit to the reported means. $CT_{max}$ values were not included in model fits; in cases where original studies included $CT_{max}$ as a data point within a TPC, we intentionally excluded it during data extraction.

The Norberg model (20) as modified by (21) (referred to as the Norberg-Thomas model), models temperature-dependent performance (*f(T)*) as the product of an exponential component and a declining quadratic component (Equation S3) as:

$$f(T) = ae^{bT}\left[1 - \left(\frac{T - T_{ref}}{\frac{c}{2}}\right)^2\right], \quad (S3)$$

where $a$ acts as a scaling parameter, $b$ determines the slope of the exponential rise and the direction of the curve skew (negative values lead to right-skew), $c$ captures the breadth of positive performance, $T_{ref}$ is the temperature at which the declining quadratic function peaks, and $T$ is temperature (°C). While many models to fit TPCs exist, no one model consistently outperforms others (3). We chose to use the Norberg-Thomas model because it estimates only four parameters, making it feasible to fit TPCs to datasets with as few as five data points. The model is flexible enough to capture both left- and right-skewed curves and allows direct derivation or estimation of key thermal traits, including $T_{opt}$ and $TPC_{max}$. $TPC_{max}$ is calculated from the model parameters as $T_{ref} + (c/2)$.

We fitted TPCs in three sequential rounds, with quality control applied after each (Figure S1). Because curves yielding high confidence in $T_{opt}$ were not always the same as those providing high confidence in $TPC_{max}$, quality control was performed independently for each $T_{opt}$ and $TPC_{max}$. However, for supra-optimal range, SOR, we performed quality control simultaneously to retain curves with high confidence estimates of both $T_{opt}$ and $TPC_{max}$. In the first round of fitting, curves were fit with starting parameters informed by the data and parameter bounds set to $a$ = 0–500, $b$ = –10–10, $c$ = 0–700, and $T_{ref}$ = –100–100. In the second round of fitting, we refit curves using the same protocol (nlsLM) but using starting parameters from the results of 500 iterations of a constrained random search using the mulstart_fit function from the *nls.multstart* R package (22, 23). In the third round of fitting, we used the same fitting protocol as the second-round, but constrained $b$ to positive values (0–10) to enforce left-skewed fits.

After each round of fitting, we quantified uncertainty in our TPC fits by generating 10,000 non-parametric bootstrap curves (Boot function, *car* in R, (24, 25)) per TPC. We calculated TPC traits ($TPC_{max}$, $T_{opt}$) from each bootstrapped fit. We calculated $TPC_{max}$ by setting the second term in Equation S3 equal to 0 and solving for $T$, ($TPC_{max} = T_{ref} + c/2$). We estimated $T_{opt}$ from the fitted TPC using numerical optimization (via the optim function in R). We excluded TPCs from our final dataset if parameter estimates fell within 0.1% of their bounds that we set in the model fitting (i.e. "bound-hitting" parameter estimates) (Figure S1iv) or if they had uncertain trait estimates (95% trait CI >15% of the TPC thermal breadth) (Figure S1v). We retained curves with

≥1,000 valid bootstrap samples. Curves failing these criteria were refitted in the second, then the third rounds of our pipeline.

From the curves that passed these quality control steps, we retained 1,000 bootstrapped parameter fits (and corresponding trait estimates) by randomly drawing from those falling within the 95% confidence interval of the focal trait. As a final step, we excluded curves whose resulting 95% CI in the trait of interest exceeded 5°C in width (Figure S1v). Therefore, our final datasets only included successfully fit curves with 1,000 randomly sampled parameter estimates from our bootstrapping and corresponding $TPC_{max}$/$T_{opt}$ estimates. SOR was calculated for each TPC as the difference between $TPC_{max}$ and $T_{opt}$ for each of the 1000 bootstrapped curves that passed the simultaneous quality control.

**iii. Data summaries**

Across all sources, we compiled 3349 thermal performance datasets from 1584 species units (i.e., including species with unresolved taxonomy), 8215 $CT_{max}$ estimates from 3109 species units, and 772 ARR estimates from 639 species units. Following filtering performance datasets with fewer than five temperature treatments or without observations on both sides of an observed performance peak, and resolving taxonomy across datasets (Figure S1ii), we retained, we retained 2,069 TPCs for 850 species, 8077 $CT_{max}$ estimates for 2932 species, and 678 ARR estimates for 475 species. Following model fitting and quality control (Figure S1iv,v), we retained 204 TPCs (102 species) with $T_{opt}$ estimates, and 159 TPCs (79 species) with $TPC_{max}$ estimates, and 136 TPCs (68 species) were retained for supra-optimal range, SOR, calculations (Figure S1vi). The SOR dataset was also used to compare variation among thermal traits, as it is comprised of high confidence estimates of both $T_{opt}$ and $TPC_{max}$ as well as SOR. These final datasets spanned nine taxonomic classes and included locomotor and bioenergetic performance measures, organismal and sub-organismal measurements, aquatic and terrestrial taxa, temperate and tropical climates, and acute and prolonged exposures (Figure S3). For analyses evaluating specific environmental and experimental contexts, we restricted individual models to categories with sufficient representation and relevant biological comparability. Thus, some TPCs were omitted from particular moderator analyses but retained in the primary analyses and in other context-specific analyses for which they met the relevant inclusion criteria.

To evaluate the robustness of TPC fitting and trait estimation to data limitations and model choice, we conducted a series of sensitivity analyses described below.

**iv. Sensitivity Analyses**

**iv.a. Sensitivity to limited sampling above $T_{opt}$.**

To evaluate whether limited sampling above $T_{opt}$ affects model fit and $TPC_{max}$ estimation, we conducted a simulation-based sensitivity analysis and an empirical-based resampling sensitivity analysis. For the simulation-based approach, we generated datasets from the Norberg-Thomas model of thermal performance with set parameters (*a* = 10, *b* = 0.05, *c* = 18, *Tref* = 25°C, resulting in a $TPC_{max}$ = 34°C). We then systematically varied (i) the number of temperature treatments above $T_{opt}$ (ranging from 1 to 4 temperature levels above $T_{opt}$), (ii) the position of those observations relative to $T_{opt}$ (i.e., the retention of the maximum temperature sampled), and (iii) the level of observational noise in the generated dataset (SD = 1, 3, 6 10). These datasets were subsequently fitted using the same nonlinear fitting/bootstrapping pipeline and passed through the same quality control (QC) as the empirical data (described in section ii above and illustrated in Figure S1).

The results from these simulations indicate that the number of observations above $T_{opt}$ influences the precision and accuracy of model fits and $TPC_{max}$ (Figure S8), particularly when sampling in this region is sparse. However, we find that even a small number of well-positioned observations above $T_{opt}$—particularly observations of performance near zero performance—can provide sufficient constraint to recover the high-temperature decline of the curve with reasonable accuracy (Figure S8B, 95% confidence intervals overlapping the 'true' estimate). Importantly, fits that were poorly constrained due to insufficient or poorly positioned high-temperature data consistently demonstrated $TPC_{max}$ confidence intervals greater than the threshold of our inclusion criteria for fit quality (Figure S8C). As a result, low-confidence trait estimates from limited high-temperature sampling would be excluded from downstream analyses.

For the empirical resampling analysis, we used a subset of TPCs from our empirical synthesized dataset that contained extensive observations above $T_{opt}$. These well-sampled curves served as reference datasets from which we could simulate progressively reduced sampling in the temperatures beyond $T_{opt}$. We selected four representative performance datasets (CIEE 357, CIEE 358, CIEE 387, and KON 40686; Figure S9A). For each curve, we first fit the Norberg-Thomas model to all available observations to estimate $T_{opt}$ and $TPC_{max}$ as described above (SI A. ii. *TPC fitting and quality control).* The TPC traits derived from the model fit to this full dataset served as the benchmark against which we compared subsampled fits.

We generated reduced datasets by retaining all observations at or below $T_{opt}$ and systematically reducing the hottest temperature observation until $T_{opt}$. For each reduced dataset, we refit the Norberg-Thomas model, estimated $TPC_{max}$ using the same methods described in the section *ii. TPC fitting and quality control*, and its uncertainty via residual bootstrapping using the Boot function in the *car* package in R (25), and used the same filters described in our quality control pipeline (SI A. ii. *TPC fitting and quality control*, Figure S1). $TPC_{max}$ estimates and their confidence intervals were then compared to the benchmark full-data trait estimate to assess sensitivity to high-temperature sampling density.

Consistent with the results our model-generated simulation results, we found that estimates of $TPC_{max}$ were sensitive to the number of observations beyond $T_{opt}$. As the number of excluded hottest temperature treatments increased, the estimates of $TPC_{max}$ became less accurate (were further from the benchmark $TPC_{max}$) and had wider 95% CIs (Figure S9B). Confidence intervals for some of these resampled curves spanned over 100°C, indicating substantial parameter uncertainty and poor TPC fits to the descending portion of the performance curve. Importantly, across all four resampled datasets, $TPC_{max}$ estimates with high uncertainty that were the result of decreased sampling of high temperatures leading to poor TPC fits were identified and excluded from our downstream analyses by our quality control pipeline (Figure S9B, red points).

**iv.b. Sensitivity to limited data and TPC model formulation.**

To evaluate how thermal performance curve models perform (in terms of accuracy and precision of TPC trait estimates) in cases where data are limited, we conducted simulation-based analyses using generated thermal performance data. Given that temperatures above $T_{opt}$ may be under-sampled in the literature, we were interested in how these models differ in their estimates of $TPC_{max}$. Specifically, we compared the performance of four commonly used TPC models with four estimated parameters: the Norberg-Thomas model (described above), the Brière II model, the high-temperature inactivation Sharpe–Schoolfield model, and the flexTPC model.

The Brière model II (26) describes temperature-dependent performance as an asymmetric function with an explicit upper limit (Equation S4):

$$f(T) = a\,T\,(T - T_{\mathrm{min}})\sqrt{T_{\mathrm{max}} - T}, \qquad (S4)$$

where $a$ is a scaling parameter, $b$ is a shape parameter, $T_{min}$ and $T_{max}$ the lower and upper temperatures at which performance is zero, respectively, and $T$ is temperature (°C). Performance is constrained to zero for $T_{min}$ and $T_{max}$. The Brière model produces left-skewed curves and directly estimates $TPC_{max}$ (referred to as $T_{max}$ in the equation) as a fitted parameter.

The flexTPC model (27) describes temperature-dependent performance using a flexible asymmetric function with explicit lower and upper thermal limits, maximum performance, and parameters controlling curve shape and breadth (Equation S5),

$$f(T) = r_{\mathrm{max}}\left[\left(\frac{T-T_{min}}{\alpha}\right)^{\alpha}\left(\frac{T_{max}-T}{\alpha}\right)^{1-\alpha}\left(\frac{1}{T_{max}-T_{min}}\right)\right]^{\frac{\alpha(1-\alpha)}{\beta^2}}. \qquad (S5)$$

In this model, $T_{min}$ and $T_{max}$ define the lower and upper temperatures at which performance is zero, $r_{max}$ defines maximum performance, α is a shape parameter defining the symmetry and direction of the curve skew, and β adjusts the breadth of the curve. Like the Brière II model, flexTPC directly estimates $TPC_{max}$ as the fitted upper thermal limit, $T_{max}$.

The Sharpe–Schoolfield model (28) describes temperature-dependent performance based on enzymatic reaction rates, incorporating both activation and deactivation processes (Equation S6):

$$f(T) = \frac{r_{T_{\mathrm{ref}}} e^{\left[\frac{E}{k}\left(\frac{1}{T_{\mathrm{ref}}} - \frac{1}{T}\right)\right]}}{1+e^{\left[\frac{E_h}{k}\left(\frac{1}{T_h} - \frac{1}{T}\right)\right]}}, \quad (S6)$$

where $r_{T_{\mathrm{ref}}}$ is the rate at a reference temperature $T_{ref}$, $E$ is the activation energy governing the exponential increase in performance, $E_h$ is the deactivation energy at high temperatures, $T_h$ is the temperature at which half of enzyme activity is lost due to thermal inactivation, and $k$ is the Boltzmann constant. Here, temperature, $T$, is in kelvin. In our implementation, $T_{ref}$ was fixed at 15°C and the remaining four parameters were estimated from the generated data. Unlike the Norberg-Thomas model or the Brière model, the Sharpe-Schoolfield model does not explicitly model a temperature at which performance drops to zero and indeed cannot model negative values. Thus, we defined $TPC_{max}$ as the upper temperature where performance drops below 1% of its maximum

We simulated thermal performance datasets using each model, with parameter values chosen to produce biologically realistic thermal performance curves and comparable upper thermal limits. The generated thermal performance data had a true $TPC_{max}$ of 34; Gaussian noise (SD = 2) was added to simulated performance values to approximate empirical variability. We constructed sampling scenarios that varied in the number of temperature observations above the temperature of peak performance ($T_{opt}$), as this region is often under-sampled relative to the rest of the curve and is important for $TPC_{max}$ estimates (see section iii below). Our simulated performance datasets consisted of eleven temperature levels between 15°C and 35°C (Figure S10A). In one scenario we included all eleven temperature levels, with the maximum temperature sampled at 35°C, (i.e., “Max 35”), in one scenario we included 10 temperature levels, with the maximum temperature sampled at 33°C (i.e. “Max 33”), and one scenario in which we included 9 temperature levels, with the maximum at 31°C. (i.e. “Max 31”). Each simulated dataset was then fitted using all candidate models with residual bootstrapping used to estimate uncertainty in $TPC_{max}$ for each fitted model.

Our goal in comparing candidate TPC models was not simply to identify the model that produced the narrowest confidence intervals. Rather, we sought a model that performed well when high-temperature sampling was adequate and that either failed to fit or produced increasing uncertainty when high-temperature sampling was insufficient to constrain the descending limb of the TPC. This behavior was important because overly precise estimates under sparse sampling could allow poorly constrained $TPC_{max}$ estimates to pass downstream quality control.

The Brière II model converged across sampling scenarios and often produced finite estimates of $TPC_{max}$ (Figure S10B). However, convergence did not always indicate that $TPC_{max}$ was well constrained by the data. Because Brière II estimates $TPC_{max}$ directly as the fitted parameter $T_{max}$, it can return an upper thermal limit even when the descending limb is weakly sampled, potentially reflecting the decreased parameter space used to estimate $TPC_{max}$ relative to the other models. Across generating models, Brière II confidence interval widths ranged from 1.24–4.13 under Max 35 sampling, 1.53–13.42 under Max 33 sampling, and 7.30–48.81 under Max 31 sampling. In several cases, Brière II confidence intervals widened as high-temperature sampling was reduced, such as for Brière-generated data, where CI width increased from 3.48 under Max 35 sampling to 13.42 under Max 33 sampling. However, in other cases Brière II produced finite estimates with comparatively modest uncertainty despite reduced high-temperature coverage, suggesting that its apparent precision may partly reflect model structure rather than strong empirical constraint on the upper thermal limit (Figure S10C, D). For example, under Max 31 sampling, Brière II estimated $TPC_{max}$ between 32.66 and 39.36 for three of the four generating models but produced an implausibly high estimate for Schoolfield-generated data (mean = 68.94, 95% CI = 31.19–80.00). Under the sparsest Schoolfield-generated scenario, Brière II produced an implausibly high $TPC_{max}$ estimate with an upper confidence interval at the imposed search limit, further indicating sensitivity to model mismatch and sparse descending-limb data.

The flexTPC model also includes an explicit upper thermal limit and generally converged across sampling scenarios (Figure S10B). When fitted to flexTPC-generated data with adequate high-temperature sampling, flexTPC recovered $TPC_{max}$ with relatively narrow confidence intervals. However, flexTPC did not consistently improve upper-limit estimation under model mismatch. When fitted to Thomas-generated data, flexTPC produced broad confidence intervals even when the highest temperature treatment was retained, and uncertainty increased further as high-temperature observations were removed. Across generating models, flexTPC confidence interval widths ranged from 1.40–14.09 under Max 35 sampling, 1.40–46.49 under Max 33 sampling, and 31.09–48.80 under Max 31 sampling. Under reduced high-temperature sampling, flexTPC estimates were also sometimes influenced by the imposed upper search limit. For example, upper confidence limits reached 80.00 for Brière-generated data under Max 33 sampling and for Thomas- and flexTPC-generated data under Max 31 sampling. Thus, the additional flexibility of flexTPC did not remove the need for strong empirical constraint on the descending limb of the curve (Figure S10C, D).

The Norberg-Thomas model showed intermediate performance. It generally produced accurate and precise $TPC_{max}$ estimates when high-temperature data were available, and both uncertainty and failure rates increased under limited sampling (Figure S10B-D). This pattern was central to our model choice. For Thomas-generated data, Norberg-Thomas estimates remained close to the true $TPC_{max}$ even when the maximum temperature sampled was reduced, but confidence intervals widened as high-temperature coverage declined. For example, CI width increased from 0.34 under Max 35 sampling to 0.83 under Max 33 and 2.41 °C under Max 31. Norberg-Thomas also performed well when fitted to flexTPC-generated data, although uncertainty increased under the sparsest sampling scenario. For flexTPC-generated data, CI width increased from 0.31 °C under Max 35 sampling to 1.00 under Max 33 and 4.36 under Max 31. In contrast, Norberg-Thomas model convergence was particularly sensitive to data generated from the Sharpe-Schoolfield model—where performance exhibits a steep decline after $T_{opt}$—as where fits either failed or produced extremely wide, biologically unrealistic confidence intervals.

In contrast, the Sharpe-Schoolfield model exhibited strong sensitivity to the absence of data on the declining limb of the thermal performance curve. Sharpe-Schoolfield recovered $TPC_{max}$ well when fitted to Schoolfield-generated data with adequate high-temperature sampling, but failure rates increased sharply when the descending limb was poorly sampled and when the model was fitted to data generated by other curve forms (Figure S10B-D). For example, failure rates reached 0.95 for Thomas-generated data under Max 31 sampling, 0.94 for Brière-generated data under Max 33–31 sampling, and 1.00 for flexTPC-generated data under Max 31 sampling. These results highlight that the Sharpe-Schoolfield model requires dense data coverage, particularly at high temperatures, to yield reliable estimates of $TPC_{max}$.

Together, these results demonstrate that the most useful model for our empirical pipeline was not necessarily the model that always returned the most precise $TPC_{max}$ estimate, but the model that performed well when data were informative and reflected increasing uncertainty when high-temperature sampling was limited. While Brière II and flexTPC may output finite estimates of $TPC_{max}$ under data-poor conditions, these estimates are not necessarily well constrained by the data. The Norberg-Thomas model generally performed well when high-temperature data were available and, importantly, produced wider confidence intervals or fitting failures as high-temperature sampling became insufficient. This diagnostic behavior is preferable for our QC pipeline to identify and exclude low-confidence $TPC_{max}$ estimates before downstream analyses. Further, the Norberg-Thomas model can accommodate a wide range of curve shapes, including asymmetrical and nearly symmetrical curves, providing an inclusive framework for fitting empirical data. Accordingly, we used the Norberg-Thomas model for all fitting of empirical data. Across all models, the decline in accuracy and precision under sparse sampling underscores the importance of implementing rigorous quality control to exclude biologically unrealistic or poorly constrained $TPC_{max}$ estimates.

**iv.c. Sensitivity to model choice (empirical comparison).**

We evaluated whether our biological inferences based on $TPC_{max}$ were robust to our choice of TPC model in two distinct steps. First, we evaluated relative model support across our full paired TPC-$CT_{max}$ dataset (N = 410), which includes all empirical TPCs with associated species $CT_{max}$ estimates prior to quality control (QC) filtering (see data pipeline Figure S1). We compared model fit and $TPC_{max}$ estimates from the Norberg-Thomas model, the Briere II model, the flexTPC model, and Sharpe-Schoolfield model. We fitted all four models to all empirical datasets (N = 410) as described above in SI Methods A.ii and determined the best-supported model for each dataset using AIC. Then, we tested that our biological inferences remained consistent by testing for trait correlations between absolute $TPC_{max}$ and the dataset rank in $TPC_{max}$ produced by each model within our $TPC_{max}$ hypothesis-testing dataset (N = 159; Figure S1).

When comparing model fits across our full paired dataset (N = 410), the best-supported model varied among empirical TPCs, indicating that no single candidate model universally provided the best empirical fit, consistent with (3). While some models frequently achieved a lower AIC for particular datasets, goodness-of-fit (AIC) does not always equate to accurate or precise estimates of $TPC_{max}$.

$TPC_{max}$ estimates were generally consistent between the Norberg-Thomas model and the Brière II and Sharpe-Schoolfield models, but less consistent between Norberg-Thomas and flexTPC (Figure S11). For the N = 159 datasets used in our $TPC_{max}$ hypothesis-testing analyses, Norberg-Thomas $TPC_{max}$ estimates were strongly correlated with Sharpe-Schoolfield estimates (Pearson's r = 0.98; Spearman's ρ = 0.97) and moderately to strongly correlated with Brière II estimates (Pearson's r = 0.80; Spearman's ρ = 0.86). In contrast, Norberg-Thomas and flexTPC estimates showed weaker agreement, particularly for absolute $TPC_{max}$ values (Pearson's r = 0.34; Spearman's ρ = 0.74). This weaker agreement reflected several empirical curves for which flexTPC produced very high $TPC_{max}$ estimates relative to the other fitted models, indicating that $TPC_{max}$ was not consistently well constrained by this model in our empirical dataset. These empirical results are consistent with the simulation results above, in which flexTPC estimates were sensitive to model mismatch and sparse high-temperature sampling. These results also suggest that our downstream inference is robust to the use of Brière II or Sharpe-Schoolfield for $TPC_{max}$ estimation, but less robust to flexTPC because flexTPC produced divergent and sometimes biologically implausible upper-limit estimates for some empirical curves. Accordingly, we retained the Norberg-Thomas model as the primary model for empirical trait estimation.

**iv.d. Sensitivity analysis of $CT_{max}$ endpoints.**

Because studies differed in the endpoint used to define $CT_{max}$, we retained all reported measures of upper thermal tolerance to maximize taxonomic coverage. Plant studies included endpoints such as Tmax, LT50, LT100, and Tcrit, whereas animal studies included loss of equilibrium (LOE), loss of righting response (LRR), onset of spasms, paralysis, response to prodding, opercular flaring, and death. To evaluate the potential influence of endpoint choice, we conducted a sensitivity analysis using the $T_{opt}$-$CT_{max}$ dataset because this analysis retained the largest number of TPCs (N = 204) and species (N = 102). To evaluate the potential influence of endpoint choice, we first examined species for which multiple $CT_{max}$ endpoints were reported. Of the 102 species in this dataset, only 38 had multiple endpoints of $CT_{max}$, and all 38 species were animals. Among the 38 species with more than one endpoint, 15 species showed differences in mean $CT_{max}$ greater than 3°C across endpoints (Figure S12).

We evaluated the robustness of our primary results by refitting the $T_{opt}$-$CT_{max}$ meta-regression models using (1) all $CT_{max}$ endpoint estimates, (2) non-lethal endpoints only, and (3) loss of righting response/loss of equilibrium (LRR/LOE) endpoints only. We refit the $T_{opt}$-$CT_{max}$ meta-regression models for each endpoint definition, including taxonomic class as a fixed effect and species as a random intercept. To distinguish effects of endpoint definition from changes in sample composition, we also compared each restricted endpoint model against the all-endpoint model using the same matched set of species.

Across endpoint definitions, slope estimates were similar and 95% confidence intervals overlapped substantially (Figure S12), indicating that the $T_{opt}$-$CT_{max}$ relationship was robust to variation in $CT_{max}$ endpoint definitions. This suggests that retaining all reported $CT_{max}$ endpoints in the primary analyses increased taxonomic coverage without strongly influencing the estimated relationship between $T_{opt}$ and $CT_{max}$.

**v. Supplementary statistical analyses**

Model fit and heterogeneity metrics

We assessed model fit using AIC, the Omnibus Test of Moderators (QM), and a *pseudo*-$R^2$ metric. We interpreted lower AIC relative to a null model including taxonomic Class as a fixed effect and species as a random intercept as improved fit. The QM statistic tests whether model predictors collectively explain heterogeneity; a significant QM test ($p < 0.05$) indicates that the predictors explain significant variation in the response. We calculated *pseudo*-$R^2$ as the proportional reduction in species-level random-effect variance between the null and predictor models. We evaluated AIC, QM, and pseudo-$R^2$ jointly to assess model fit and variance structure.

Log-log scaling analyses

Primary analyses were conducted on the Celsius scale because $CT_{max}$, $T_{opt}$, and $TPC_{max}$ are measured in the same units, allowing direct interpretation of absolute thermal differences. As a complementary analysis, we repeated the TPC trait–$CT_{max}$ models using log-transformed absolute temperatures (Kelvin) to evaluate multiplicative, rather than additive, scaling relationships. Whereas the Celsius models estimate the expected change in a TPC trait associated with a 1°C change in $CT_{max}$, log–log Kelvin models estimate proportional scaling across absolute temperatures. The log-log (Kelvin-scale) analyses produced qualitatively similar results to the primary analyses for the $TPC_{max}$–$CT_{max}$ relationship but indicated sub-linear scaling of $T_{opt}$ with $CT_{max}$ (Tables S13 and S14). We therefore focus interpretation on Celsius-scale models because they directly represent the temperature scale on which these traits were measured and provide the clearest and most intuitive biological interpretation of trait differences. By contrast, log transformation of absolute temperature compresses variation across the observed range and makes effect sizes less directly interpretable in terms of organismal thermal biology.

Piecewise threshold analyses

To evaluate whether relationships between TPC-derived thermal traits and $CT_{max}$ exhibited nonlinear or threshold behavior, we conducted exploratory piecewise meta-regression analyses using the same weighting and random-effects structure as the primary models. Models included taxonomic Class as a fixed effect and species as a random intercept to account for non-independence among observations. We conducted profile-likelihood grid searches across candidate breakpoints ($\psi$), evaluating values from 15 to 65 °C in 0.5 °C increments for $T_{opt}$ and from 21.05 to 65 °C in 0.1 °C increments for $TPC_{max}$. At each candidate breakpoint, we constructed pre- and post-breakpoint linear segments and refit the model, selecting the breakpoint that maximized model log-likelihood. Piecewise analyses provided no evidence for a high-temperature threshold in the $T_{opt}$–$CT_{max}$ relationship. The $TPC_{max}$ analysis identified a significant slope change at a low $CT_{max}$ value ($\psi$ = 27.05 °C), but this estimate was informed by only 20 observations from seven species below the breakpoint. We therefore interpret this result cautiously and retain continuous linear models as the primary representation of both relationships (Table S15).

$T_{opt}$–$TPC_{max}$ scaling relationship

As a descriptive analysis, we evaluated the relationship between $T_{opt}$ and $TPC_{max}$ within the SOR dataset, where both traits were estimated from the same fitted thermal performance curves. $T_{opt}$ and $TPC_{max}$ were strongly positively correlated (Pearson's $r = 0.86$, $p < 0.001$). A linear model indicated that $TPC_{max}$ scaled approximately 1:1 with $T_{opt}$ (slope = 0.99, 95% CI = 0.89, 1.09, $p < 0.001$).

Because both $T_{opt}$ and $TPC_{max}$ are estimated traits and share uncertainty within fitted curves, we do not interpret this descriptive regression as a full measurement-error model. Instead, we use it to evaluate whether the observed $CT_{max}$ relationships could simply reflect constrained sublinear scaling between $T_{opt}$ and $TPC_{max}$. The descriptive relationship provides no support for this explanation. We separately propagated the bootstrap variances and covariance between $T_{opt}$ and $TPC_{max}$ in the multivariate meta-analytic model comparing their relationships with $CT_{max}$ (see main text).

Leave-one-out sensitivity analyses

We conducted leave-one-out sensitivity analyses for the primary $T_{opt}$–$CT_{max}$, $TPC_{max}$–$CT_{max}$, and SOR–$CT_{max}$ models to assess whether any trait relationships were driven by a single influential observation. For each model, we sequentially removed one observation, refit the model, and compared the resulting $CT_{max}$ slope to the full-model estimate and 95% confidence interval. Leave-one-out slope estimates were highly consistent with the full-model estimates and remained well within the corresponding 95% confidence intervals for all three traits: $T_{opt}$–$CT_{max}$ full-model slope = 0.81 [0.61, 1.02], leave-one-out range = 0.78–0.90; $TPC_{max}$–$CT_{max}$ full-model slope = 0.95 [0.69, 1.20], leave-one-out range = 0.91–1.00; SOR–$CT_{max}$ full-model slope = 0.26 [0.01, 0.52], leave-one-out range = 0.21–0.31. These results indicate that the observed trait–$CT_{max}$ relationships were not strongly influenced by any single observation.

#### 2. Supplementary Lists

**i.** Categorization of TPC traits into Response Type categories.

Traits classified as bioenergetic processes:

- Adjusted daily food consumption
- Aerobic scope / Scope for activity
- Bacterial killing ability
- Development time
- Energy / Feed intake (Includes standardized rates)
- Gene expression (Mean log)
- Growth efficiency
- Growth rate (Includes specific, weight-based, and tadpole)
- Heart rate (bpm)
- HSP 70 synthesis
- Mass-specific growth rate
- Photosynthesis rate (Includes net rates)
- Respiration rate
- Rate of recovery
- Spleen weight (growth)
- Total live length
- Weight gain

Traits classified as locomotor processes:

- Burst swim speed (max and unit variants)
- Distance jumped / Jumping distance
- Distance moved in 40ms
- Distance running / Endurance running
- Foraging efficiency
- Jump speed / Velocity jumping
- Maximum angular speed
- Maximum speed
- Maximum sustained swimming speed
- Median migration rate
- Muscle contraction
- Running speed / Sprint speed
- Stamina
- Swimming speed
- Swimming stroke length
- Walking speed

Traits classified as life history processes:

- Colony weight
- Egg to adult viability

- Emergence
- Fecundity
- Mean percentage of fertilized eggs

**ii. Categorization of TPC traits into Exposure Duration categories.**

Exposure duration was categorized by consulting the main text of the publication associated with each thermal performance dataset. TPC assays lasting more than 30 minutes were categorized as prolonged, whereas assays lasting less than 30 minutes were categorized as acute. After consolidating differences in trait naming and measurement units, the categories were as follows.

Traits classified as acute:

- Aerobic scope / Scope for activity
- Bacterial killing ability
- Burst swimming speed (includes maximum and unit variants)
- Chill-coma recovery
- Choline concentration (proxy for protein synthesis)
- Critical swimming speed
- Distance jumped / Jumping distance
- Distance moved in 40 ms
- Distance running
- Heart rate
- HSP70 synthesis
- Jump speed / Jumping velocity
- Maximum acceleration
- Maximum angular speed
- Maximum oxygen consumption
- Maximum speed
- Muscle contraction
- Photosynthesis rate (includes net rates)
- Respiration rate
- Running speed / Sprint speed
- Spontaneous swimming activity
- Stamina
- Swimming speed
- Swimming stroke length
- Walking speed

Traits classified as prolonged:

- Adjusted daily food consumption
- Colony weight
- Consumption probability
- Critical holding velocity
- Development time

- Egg-to-adult viability
- Emergence
- Endurance running
- Energy / Feed intake (includes standardized rates)
- Fecundity
- Food-mass conversion efficiency
- Foraging efficiency
- Gene expression (mean log; *cg10910*, *mag*, and *mur29b*)
- Growth by weight
- Growth efficiency
- Growth rate (includes mass-specific, specific, standardized weight-specific, and tadpole growth rates)
- Maximum sustained swimming speed
- Mean percentage of fertilized eggs
- Median migration rate
- Spleen weight relative to body weight
- Total live length
- Weight gain

**iii.** List of search terms used, with “temperature OR thermal OR thermos*”, when querying the Web of Science Core Collection for our systematic literature search.

Topic: (temperature OR thermal OR thermo*) AND

Topic: ("undulation" OR "gill beat" OR "gill particle transport" OR "growth rate" OR "population voluntary activity" OR "population voluntary movement" OR "strike rate" OR "wing beat" OR "fecundity" OR "mortality" OR "population density" OR "population growth rate" OR "filtration rate" OR "consumption rate" OR "foraging rate" OR "handling rate" OR "parasitization rate" OR "ammonia excretion rate" OR "development rate" OR "metabolic rate" OR "oxygen consumption" OR "intrinsic growth rate" OR "juvenile development rate") AND

All fields: (insect OR fly OR flies OR mussel OR clam OR snail OR isopod OR zooplankton OR rotifer OR invertebrate OR gastropod OR mollusc* OR isopod OR pest OR beetle OR echinoderm* OR annelid OR daphnia OR arthropod* OR crustacea* OR nematode* OR roundworm* OR worm*).

#### Part B: Supplementary Tables

**Table S1. Results from a multi-level regression model evaluating the relationships between $T_{opt}$ and $CT_{max}$ (N = 204 TPCs, 102 species).**

$CT_{max}$ was standardized to a 15 °C acclimation temperature using phylum-specific acclimation response ratios (ARRs) and centered (mean $CT_{max}$ = 37.42°C). Class was included as fixed effect covariate to account for taxonomic differences and species was included as a random intercept to account for non-independent sampling of TPCs. Slopes and 95% confidence intervals describe the scaling relationships of $T_{opt}$ with $CT_{max}$.

| **Term** | **Estimate [95% CI]** | **SE** | **Z statistic** | **P-value** |
|---|---|---|---|---|
| Intercept | 23.86 [21.35, 26.38] | 1.28 | 18.61 | <0.0001 |
| Mean $CT_{max}$ | 0.82 [0.61, 1.02] | 0.1 | 7.8 | <0.0001 |
| Class: Amphibia | 2.48 [-0.33, 5.29] | 1.43 | 1.73 | 0.08 |
| Class: Arachnida | -1.52 [-6.30, 3.25] | 2.44 | -0.63 | 0.53 |
| Class: Bivalvia | 5.90 [-2.82, 14.62] | 4.45 | 1.33 | 0.18 |
| Class: Gastropoda | 16.04 [7.29, 24.79] | 4.46 | 3.59 | <0.001 |
| Class: Insecta | -0.26 [-5.18, 4.67] | 2.51 | -0.1 | 0.92 |
| Class: Liliopsida | -4.14 [-11.94, 3.66] | 3.98 | -1.04 | 0.30 |
| Class: Magnoliopsida | -13.32 [-20.07, -6.57] | 3.44 | -3.87 | 0.0001 |
| Class: Reptilia | 8.11 [4.69, 11.53] | 1.75 | 4.65 | <0.0001 |
| Species variance | 18.97 | | | |
| *QM test (moderators)* | *269.43 (df = 9)* | | | *<0.0001* |
| *QE test (residual heterogeneity)* | *45037.03 (df = 194)* | | | *<0.0001* |
| *Pseudo R2* | *39.40%* | | | |
| *Residual heterogeneity explained* | *42.80%* | | | |
| *delta AIC (moderated - null)* | *-50.61* | | | |

**Table S2. Results from a multi-level regression model evaluating the relationships between $TPC_{max}$ and $CT_{max}$ (N = 159 TPCs, 79 species).**

$CT_{max}$ was standardized to a 15°C acclimation temperature using phylum-specific acclimation response ratios (ARRs) and centered (mean $CT_{max}$ = 38.12℃). Class was included as fixed effect covariate to account for taxonomic differences and species was included as a random intercept to account for non-independent sampling of TPCs. Slopes and 95% confidence intervals describe the scaling relationships of $TPC_{max}$ with $CT_{max}$.

| Term | Estimate [95% CI] | SE | Z statistic | P-value |
|---|---|---|---|---|
| Intercept | 36.38 [33.25, 39.51] | 1.6 | 22.8 | <0.0001 |
| Mean $CT_{max}$ | 0.95 [0.69, 1.20] | 0.13 | 7.24 | <0.0001 |
| Class: Amphibia | 0.22 [-3.18, 3.62] | 1.73 | 0.13 | 0.90 |
| Class: Arachnida | -2.65 [-7.99, 2.70] | 2.73 | -0.97 | 0.33 |
| Class: Bivalvia | 1.61 [-7.84, 11.07] | 4.82 | 0.33 | 0.734 |
| Class: Gastropoda | 13.91 [4.45, 23.37] | 4.83 | 2.88 | < 0.01 |
| Class: Insecta | -2.08 [-7.34, 3.19] | 2.69 | -0.77 | 0.44 |
| Class: Liliopsida | 2.84 [-5.99, 11.66] | 4.5 | 0.63 | 0.53 |
| Class: Magnoliopsida | -10.50 [-18.46, -2.55] | 4.06 | -2.59 | 0.01 |
| Class: Reptilia | 6.19 [2.01, 10.37] | 2.13 | 2.9 | <0.001 |
| Species variance | 21.88 | | | |
| *QM test (moderators)* | *201.28 (df = 9)* | | | *<0.0001* |
| *QE test (residual heterogeneity)* | *23995.24 (df = 149)* | | | *<0.0001* |
| *Pseudo R2* | *42.60%* | | | |
| *Residual heterogeneity explained* | *50.30%* | | | |
| *delta AIC (moderated - null)* | *-43.11* | | | |

701

**Table S3. Intraspecific scaling relationships between warm-end TPC-derived traits and $CT_{max}$ across Actinopterygii, Amphibia, and Reptilia.**

$CT_{max}$ was standardized to a 15°C acclimation temperature using phylum-specific acclimation response ratios (ARRs) and centered on the mean. Class was included as a fixed effect, and species as a random intercept, to account for taxonomic differences and non-independent sampling of TPCs. The full model tested a $CT_{max}$ × Class interaction to assess whether scaling slopes differed among the three best-represented Classes in the dataset: Actinopterygii, Amphibia, and Reptilia. Because the interaction was not significant, the additive model is also presented, with class-specific intercepts and a common slope. Slopes and 95% confidence intervals describe the scaling relationship between the TPC-derived trait ($T_{opt}$ or $TPC_{max}$) and $CT_{max}$.

**$T_{opt}$ (N = 141 TPCs, 80 species):**

*Interactive:*

| **Term** | **Estimate [95% CI]** | **SE** | **Z statistic** | **P-value** |
|---|---|---|---|---|
| Intercept | 24.54 [22.08, 27.00] | 1.26 | 19.54 | <0.0001 |
| Mean $CT_{max}$ | 0.89 [0.67, 1.11] | 0.11 | 7.82 | <0.0001 |
| Class: Amphibia | 1.89 [-0.96, 4.75] | 1.46 | 1.3 | 0.19 |
| Class: Reptilia | 9.10 [5.57, 12.64] | 1.8 | 5.05 | <0.0001 |
| Mean $CT_{max}$ : Amphibia | -0.01 [-0.56, 0.54] | 0.28 | -0.03 | 0.97 |
| Mean $CT_{max}$ : Reptilia | -0.66 [-1.43, 0.10] | 0.39 | -1.69 | 0.09 |
| Species variance | 13.56 | | | |
| *QM test (moderators)* | *348.59 (df = 5)* | | | *<0.0001* |
| *QE test (residual heterogeneity)* | *34809.42 (df = 135)* | | | *<0.0001* |
| *Pseudo R2* | *48.30%* | | | |
| *Residual heterogeneity explained* | *51.20%* | | | |
| *delta AIC (moderated - null)* | *-61.77* | | | |

*Additive*:

| **Term** | **Estimate [95% CI]** | **SE** | **Z statistic** | **P-value** |
|---|---|---|---|---|
| Intercept | 24.10 [21.81, 26.39] | 1.17 | 20.62 | <0.0001 |
| Mean $CT_{max}$ | 0.84 [0.64, 1.04] | 0.1 | 8.35 | <0.0001 |
| Class: Amphibia | 2.28 [-0.22, 4.78] | 1.28 | 1.79 | 0.07 |
| Class: Reptilia | 7.79 [4.68, 10.90] | 1.59 | 4.91 | <0.0001 |
| Species variance | 13.73 | | | |
| *QM test (moderators)* | *341.50 (df = 3)* | | | *<0.0001* |
| *QE test (residual heterogeneity)* | *37465.09 (df = 137)* | | | *<0.0001* |
| *Pseudo R2* | *47.70%* | | | |

| | |
|---|---|
| *Residual heterogeneity explained* | *47.50%* |
| *delta AIC (moderated - null)* | *-53.19* |

**$TPC_{max}$ (N = 101 TPCs, 56 species):**

*Interactive:*

| Term | Estimate [95% CI] | SE | Z statistic | P-value |
|---|---|---|---|---|
| Intercept | 37.97 [33.90, 42.03] | 2.07 | 18.3 | <0.0001 |
| Mean $CT_{max}$ | 1.13 [0.74, 1.53] | 0.2 | 5.6 | <0.0001 |
| Class: Amphibia | -2.04 [-6.94, 2.87] | 2.5 | -0.81 | 0.41 |
| Class: Reptilia | 6.21 [0.94, 11.47] | 2.69 | 2.31 | 0.02 |
| Mean $CT_{max}$ : Amphibia | -0.47 [-1.33, 0.39] | 0.44 | -1.06 | 0.29 |
| Mean $CT_{max}$ : Reptilia | -0.88 [-2.03, 0.27] | 0.59 | -1.5 | 0.13 |
| Species variance | 22.61 | | | |
| *QM test (moderators)* | *140.30 (df = 5)* | | | *<0.0001* |
| *QE test (residual heterogeneity)* | *18248.57 (df = 95)* | | | *<0.0001* |
| *Pseudo R2* | *37.60%* | | | |
| *Residual heterogeneity explained* | *52.30%* | | | |
| *delta AIC (moderated - null)* | *-38.18* | | | |

*Additive*:

| Term | Estimate [95% CI] | SE | Z statistic | P-value |
|---|---|---|---|---|
| Intercept | 36.49 [32.81, 40.17] | 1.88 | 19.45 | <0.0001 |
| Mean $CT_{max}$ | 0.96 [0.62, 1.30] | 0.17 | 5.57 | <0.0001 |
| Class: Amphibia | 0.14 [-3.57, 3.86] | 1.9 | 0.08 | 0.94 |
| Class: Reptilia | 6.05 [1.19, 10.91] | 2.48 | 2.44 | 0.01 |
| Species variance | 23.02 | | | |
| *QM test (moderators)* | *134.98 (df = 3)* | | | *<0.0001* |
| *QE test (residual heterogeneity)* | *18475.19 (df = 97)* | | | *<0.0001* |
| *Pseudo R2* | *36.50%* | | | |
| *Residual heterogeneity explained* | *51.70%* | | | |
| *delta AIC (moderated - null)* | *-28.63* | | | |

**Table S4. Response Type Hypothesis test.**

Results from multi-level meta-regression models testing the hypothesis that relationships between thermal performance curve (TPC) traits and $CT_{max}$ depend on the responses captured by the TPC (bioenergetic or locomotor processes). $CT_{max}$ was standardized to a 15°C acclimation temperature using phylum-specific acclimation response ratios (ARRs) and centered ($T_{opt}$ dataset mean $CT_{max}$ = 37.42℃; $TPC_{max}$ dataset mean $CT_{max}$ = 38.12℃). Class was included as fixed effect covariate to account for taxonomic differences and species was included as a random intercept to account for non-independent sampling of TPCs. Slopes and 95% confidence intervals describe the scaling relationships of TPC traits with $CT_{max}$. For each analysis, we first present the model including the TPC trait × response-type interaction. When the interaction was not significant, we present the additive model below; coefficients are interpreted from this additive model. Slopes and 95% confidence intervals describe the scaling relationships between TPC traits and $CT_{max}$.

**$T_{opt}$ (N = 171 TPCs, 91 species):**

*Interactive:*

| **Term** | **Estimate [95% CI]** | **SE** | **Z statistic** | **P-value** |
|---|---|---|---|---|
| Intercept | 25.40 [22.76, 28.03] | 1.35 | 18.87 | <0.0001 |
| Mean $CT_{max}$ | 0.95 [0.71, 1.18] | 0.12 | 8.01 | <0.0001 |
| Response Type (Locomotion) | -1.56 [-3.37, 0.25] | 0.92 | -1.69 | 0.09 |
| Class: Amphibia | 2.09 [-0.49, 4.67] | 1.32 | 1.58 | 0.11 |
| Class: Arachnida | -1.59 [-5.96, 2.78] | 2.23 | -0.71 | 0.48 |
| Class: Bivalvia | 5.60 [-2.13, 13.34] | 3.95 | 1.42 | 0.16 |
| Class: Gastropoda | 15.20 [7.42, 22.98] | 3.97 | 3.83 | < 0.001 |
| Class: Insecta | -2.96 [-8.03, 2.12] | 2.59 | -1.14 | 0.25 |
| Class: Reptilia | 7.95 [4.73, 11.17] | 1.64 | 4.84 | <0.0001 |
| Mean $CT_{max}$ : Response Type (Locomotion) | -0.09 [-0.29, 0.11] | 0.1 | -0.89 | 0.37 |
| Species variance | 14.87 | | | |
| *QM test (moderators)* | *340.34 (df = 9,NA)* | | | <0.0001 |
| *QE test (residual heterogeneity)* | *37292.51 (df = 161)* | | | <0.0001 |
| *Pseudo R2* | *55.60%* | | | |
| *Residual heterogeneity explained* | *56.70%* | | | |
| *delta AIC (moderated - null)* | *-112.3* | | | |

*Additive:*

| **Term** | **Estimate [95% CI]** | **SE** | **Z statistic** | **P-value** |
|---|---|---|---|---|
| Intercept | 24.87 [22.52, 27.22] | 1.2 | 20.75 | <0.0001 |
| Mean $CT_{max}$ | 0.89 [0.70, 1.08] | 0.1 | 9.01 | <0.0001 |

| | | | | |
|---|---|---|---|---|
| Response Type (Locomotion) | -0.90 [-2.07, 0.26] | 0.59 | -1.52 | 0.1291 |
| Class: Amphibia | 2.18 [-0.35, 4.71] | 1.29 | 1.69 | 0.091 |
| Class: Arachnida | -1.80 [-6.09, 2.49] | 2.19 | -0.82 | 0.4103 |
| Class: Bivalvia | 5.60 [-2.02, 13.22] | 3.89 | 1.44 | 0.1495 |
| Class: Gastropoda | 15.43 [7.79, 23.08] | 3.9 | 3.96 | <0.0001 |
| Class: Insecta | -2.67 [-7.64, 2.29] | 2.53 | -1.06 | 0.29 |
| Class: Reptilia | 7.76 [4.61, 10.91] | 1.61 | 4.83 | <0.0001 |
| Species variance | 14.42 | | | |
| *QM test (moderators)* | *349.79 (df = 8)* | | | *<0.0001* |
| *QE test (residual heterogeneity)* | *38573.97 (df = 162)* | | | *<0.0001* |
| *Pseudo R2* | *57.00%* | | | |
| *Residual heterogeneity explained* | *55.20%* | | | |
| *delta AIC (moderated - null)* | *-109.26* | | | |

**$TPC_{max}$ (N = 128, 67 species):**

*Interactive:*

| **Term** | **Estimate [95% CI]** | **SE** | **Z statistic** | **P-value** |
|---|---|---|---|---|
| Intercept | 32.29 [29.05, 35.54] | 1.66 | 19.51 | <0.0001 |
| Mean $CT_{max}$ | 0.74 [0.46, 1.01] | 0.14 | 5.26 | <0.0001 |
| Response Type (Locomotion) | 6.82 [4.01, 9.64] | 1.44 | 4.75 | <0.0001 |
| Class: Amphibia | 0.50 [-2.36, 3.36] | 1.46 | 0.34 | 0.73 |
| Class: Arachnida | -5.31 [-9.90, -0.72] | 2.34 | -2.27 | 0.02 |
| Class: Bivalvia | 3.61 [-4.03, 11.25] | 3.9 | 0.93 | 0.35 |
| Class: Gastropoda | 16.76 [9.06, 24.45] | 3.92 | 4.27 | <0.0001 |
| Class: Insecta | 3.09 [-2.36, 8.54] | 2.78 | 1.11 | 0.27 |
| Class: Reptilia | 3.53 [-0.23, 7.30] | 1.92 | 1.84 | 0.07 |
| Mean $CT_{max}$ : Response Type (Locomotion) | 0.16 [-0.09, 0.41] | 0.13 | 1.27 | 0.20 |
| Species variance | 14.17 | | | |
| *QM test (moderators)* | *287.20 (df = 9)* | | | *<0.0001* |
| *QE test (residual heterogeneity)* | *12341.09 (df = 118)* | | | *<0.0001* |
| *Pseudo R2* | *60.30%* | | | |
| *Residual heterogeneity explained* | *65.50%* | | | |
| *delta AIC (moderated - null)* | *-72.95* | | | |

*Additive:*

| **Term** | **Estimate [95% CI]** | **SE** | **Z statistic** | **P-value** |
|---|---|---|---|---|
| Intercept | 33.30 [30.46, 36.14] | 1.45 | 22.99 | <0.0001 |
| Mean $CT_{max}$ | 0.83 [0.59, 1.07] | 0.12 | 6.89 | <0.0001 |
| Response Type (Locomotion) | 5.40 [3.64, 7.16] | 0.9 | 6.02 | <0.0001 |

| | | | | |
|---|---|---|---|---|
| Class: Amphibia | 0.20 [-2.61, 3.02] | 1.44 | 0.14 | 0.89 |
| Class: Arachnida | -4.79 [-9.29, -0.28] | 2.3 | -2.08 | 0.04 |
| Class: Bivalvia | 3.52 [-4.10, 11.14] | 3.89 | 0.91 | 0.36 |
| Class: Gastropoda | 16.29 [8.66, 23.93] | 3.9 | 4.18 | <0.0001 |
| Class: Insecta | 2.33 [-2.98, 7.65] | 2.71 | 0.86 | 0.39 |
| Class: Reptilia | 4.11 [0.46, 7.76] | 1.86 | 2.21 | 0.03 |
| Species variance | 14.09 | | | |
| *QM test (moderators)* | *287.08 (df = 8)* | | | *<0.0001* |
| *QE test (residual heterogeneity)* | *13465.97 (df = 119)* | | | *<0.0001* |
| *Pseudo R2* | *60.50%* | | | |
| *Residual heterogeneity explained* | *62.40%* | | | |
| *delta AIC (moderated - null)* | *-69.04* | | | |

**Table S5. Biological Scale Hypothesis test.**

Results from a multi-level meta-regression model testing the hypothesis that relationships between thermal performance curve (TPC) optimum temperature ($T_{opt}$) and $CT_{max}$ depend on biological scale (organismal vs. sub-organismal). $CT_{max}$ was standardized to a 15°C acclimation temperature using phylum-specific acclimation response ratios (ARRs) and centered (mean $CT_{max}$ = 37.42°C). Class was included as fixed effect covariate to account for taxonomic differences and species was included as a random intercept to account for non-independent sampling of TPCs. Slopes and 95% confidence intervals describe the scaling relationships of $T_{opt}$ with $CT_{max}$. For models where interaction terms were non-significant, results are also reported for the additive model. For these models, the organismal scale is the reference group and Actinopterygii is the reference class. Note: we did not test this hypothesis for $TPC_{max}$-$CT_{max}$ relationships.

**$T_{opt}$ (N = 202 TPCs, 101 species):**

*Interactive:*

| Term | Estimate [95% CI] | SE | Z statistic | P-value |
|---|---|---|---|---|
| Intercept | 24.30 [21.77, 26.84] | 1.29 | 18.81 | <0.0001 |
| Mean $CT_{max}$ | 0.84 [0.62, 1.06] | 0.11 | 7.37 | <0.0001 |
| Level of Response (Sub-Organismal) | -3.64 [-4.19, -3.09] | 0.28 | -12.92 | <0.0001 |
| Class: Amphibia | 2.41 [-0.39, 5.21] | 1.43 | 1.69 | 0.09 |
| Class: Arachnida | -2.02 [-6.79, 2.75] | 2.43 | -0.83 | 0.41 |
| Class: Bivalvia | 5.67 [-2.95, 14.29] | 4.4 | 1.29 | 0.20 |
| Class: Gastropoda | 15.72 [7.08, 24.36] | 4.41 | 3.57 | <0.001 |
| Class: Insecta | -0.73 [-5.88, 4.41] | 2.62 | -0.28 | 0.78 |
| Class: Liliopsida | -2.76 [-10.47, 4.95] | 3.93 | -0.7 | 0.48 |
| Class: Magnoliopsida | -10.10 [-17.13, -3.08] | 3.58 | -2.82 | <0.01 |
| Class: Reptilia | 7.60 [4.14, 11.06] | 1.77 | 4.3 | <0.0001 |
| Mean $CT_{max}$ : Level of Response (Sub-Organismal) | -0.06 [-0.32, 0.20] | 0.13 | -0.45 | 0.66 |
| Species variance | 18.50 | | | |
| *QM test (moderators)* | *448.56 (df = 11)* | | | *<0.0001* |
| *QE test (residual heterogeneity)* | *44176.64 (df = 190)* | | | *<0.0001* |
| *Pseudo R2* | *41.30%* | | | |
| *Residual heterogeneity explained* | *43.60%* | | | |
| *delta AIC (moderated - null)* | *-226.44* | | | |

*Additive :*

| Term | Estimate [95% CI] | SE | Z statistic | P-value |
|---|---|---|---|---|
| Intercept | 24.19 [21.72, 26.66] | 1.26 | 19.18 | <0.0001 |
| Mean $CT_{max}$ | 0.82 [0.62, 1.02] | 0.1 | 7.94 | <0.0001 |
| Level of Response (Sub-Organismal) | -3.61 [-4.15, -3.07] | 0.27 | -13.15 | <0.0001 |
| Class: Amphibia | 2.49 [-0.27, 5.26] | 1.41 | 1.77 | 0.078 |
| Class: Arachnida | -1.86 [-6.56, 2.84] | 2.4 | -0.78 | 0.44 |
| Class: Bivalvia | 5.59 [-2.99, 14.16] | 4.37 | 1.28 | 0.20 |
| Class: Gastropoda | 15.72 [7.12, 24.32] | 4.39 | 3.58 | <0.001 |
| Class: Insecta | -0.66 [-5.77, 4.45] | 2.61 | -0.25 | 0.80 |
| Class: Liliopsida | -2.71 [-10.39, 4.96] | 3.92 | -0.69 | 0.49 |
| Class: Magnoliopsida | -10.60 [-17.25, -3.95] | 3.39 | -3.12 | <0.01 |
| Class: Reptilia | 7.77 [4.41, 11.14] | 1.72 | 4.53 | <0.0001 |
| Species variance | 18.34 | | | |
| *QM test (moderators)* | *450.84 (df = 10)* | | | *<0.0001* |
| *QE test (residual heterogeneity)* | *44179.02 (df = 191)* | | | *<0.0001* |
| *Pseudo R2* | *41.90%* | | | |
| *Residual heterogeneity explained* | *43.60%* | | | |
| *delta AIC (moderated - null)* | *-223.3* | | | |

**Table S6. Exposure Duration Hypothesis test.**

Results from a multi-level meta-regression model testing the hypothesis that relationships between thermal performance curve (TPC) traits ($T_{opt}$, $TPC_{max}$) and $CT_{max}$ depend on the the duration of temperature exposure in the TPC assay (acute vs prolonged). $CT_{max}$ was standardized to a 15 °C acclimation temperature using phylum-specific acclimation response ratios (ARRs) and centered ($T_{opt}$ dataset mean $CT_{max}$ = 37.42°C; $TPC_{max}$ dataset mean $CT_{max}$ = 38.12°C). Class was included as fixed effect covariate to account for taxonomic differences and species was included as a random intercept to account for non-independent sampling of TPCs. Slopes and 95% confidence intervals describe the scaling relationships of TPC traits with $CT_{max}$. For models where interaction terms were non-significant, results are also reported for the additive model. For all models, the acute exposure duration category is the reference group and Actinopterygii is the reference class.

**$T_{opt}$ (N = 204 TPCs, 102 species):**

*Interactive:*

| Term | Estimate [95% CI] | SE | Z statistic | P-value |
|---|---|---|---|---|
| Intercept | 21.81 [19.00, 24.62] | 1.43 | 15.2 | <0.0001 |
| Mean $CT_{max}$ | 0.79 [0.54, 1.04] | 0.13 | 6.1 | <0.0001 |
| Exposure Duration (Prolonged) | 6.38 [5.37, 7.39] | 0.51 | 12.4 | <0.0001 |
| Class: Amphibia | 2.93 [-0.20, 6.06] | 1.6 | 1.83 | 0.07 |
| Class: Arachnida | 0.59 [-4.71, 5.88] | 2.7 | 0.22 | 0.83 |
| Class: Bivalvia | 3.79 [-5.98, 13.56] | 4.98 | 0.76 | 0.45 |
| Class: Gastropoda | 17.95 [8.22, 27.68] | 4.96 | 3.62 | <0.001 |
| Class: Insecta | -2.80 [-8.27, 2.68] | 2.79 | -1 | 0.32 |
| Class: Liliopsida | -1.78 [-10.49, 6.94] | 4.45 | -0.4 | 0.69 |
| Class: Magnoliopsida | -10.78 [-18.47, -3.09] | 3.92 | -2.75 | <0.01 |
| Class: Reptilia | 9.67 [5.88, 13.46] | 1.93 | 5 | <0.0001 |
| Mean $CT_{max}$ : Exposure Duration (Prolonged) | 0.26 [-0.06, 0.58] | 0.16 | 1.6 | 0.12 |
| *Species variance* | *23.24* | | | |
| *QM test (moderators)* | *378.68 (df = 11)* | | | *<0.0001* |
| *QE test (residual heterogeneity)* | *36545.86 (df = 192)* | | | *<0.0001* |
| *Pseudo R2* | *25.80%* | | | |
| *Residual heterogeneity explained* | *53.60%* | | | |
| *delta AIC (moderated - null)* | *-212.22* | | | |

*Additive:*

| Term | Estimate [95% CI] | SE | Z statistic | P-value |
|---|---|---|---|---|
| Intercept | 21.60 [18.79, 24.42] | 1.44 | 15.04 | <0.0001 |
| Mean $CT_{max}$ | 0.88 [0.65, 1.11] | 0.12 | 7.58 | <0.0001 |
| Exposure Duration (Prolonged) | 6.15 [5.18, 7.11] | 0.49 | 12.48 | <0.0001 |
| Class: Amphibia | 3.22 [0.09, 6.34] | 1.59 | 2.02 | 0.04 |

| | | | | |
|---|---|---|---|---|
| Class: Arachnida | 0.59 [-4.74, 5.91] | 2.72 | 0.22 | 0.83 |
| Class: Bivalvia | 2.63 [-7.08, 12.35] | 4.96 | 0.53 | 0.59 |
| Class: Gastropoda | 18.65 [8.91, 28.39] | 4.97 | 3.75 | <0.001 |
| Class: Insecta | -2.47 [-7.96, 3.01] | 2.8 | -0.88 | 0.38 |
| Class: Liliopsida | -2.81 [-11.47, 5.86] | 4.42 | -0.63 | 0.53 |
| Class: Magnoliopsida | -12.31 [-19.80, -4.81] | 3.83 | -3.22 | <0.01 |
| Class: Reptilia | 9.69 [5.88, 13.50] | 1.94 | 4.99 | <0.0001 |
| *Species variance* | *23.49* | | | |
| *QM test (moderators)* | *374.04 (df = 10)* | | | *<0.0001* |
| *QE test (residual heterogeneity)* | *36639.93 (df = 193)* | | | *<0.0001* |
| *Pseudo R2* | *25.00%* | | | |
| *Residual heterogeneity explained* | *53.50%* | | | |
| *delta AIC (moderated - null)* | *-206.17* | | | |

**$TPC_{max}$ (N = 159, 79 species):**

*Interactive:*

| **Term** | **Estimate [95% CI]** | **SE** | **Z statistic** | **P-value** |
|---|---|---|---|---|
| Intercept | 38.00 [34.78, 41.22] | 1.64 | 23.13 | <0.0001 |
| Mean $CT_{max}$ | 0.93 [0.64, 1.22] | 0.15 | 6.25 | <0.0001 |
| Exposure Duration (Prolonged) | -4.05 [-7.11, -0.98] | 1.57 | -2.58 | 0.01 |
| Class: Amphibia | 0.43 [-2.89, 3.75] | 1.69 | 0.26 | 0.80 |
| Class: Arachnida | -4.24 [-9.51, 1.02] | 2.69 | -1.58 | 0.11 |
| Class: Bivalvia | 3.02 [-6.21, 12.26] | 4.71 | 0.64 | 0.52 |
| Class: Gastropoda | 12.21 [2.98, 21.44] | 4.71 | 2.59 | 0.01 |
| Class: Insecta | -1.33 [-6.42, 3.77] | 2.6 | -0.51 | 0.61 |
| Class: Liliopsida | 1.40 [-7.38, 10.18] | 4.48 | 0.31 | 0.75 |
| Class: Magnoliopsida | -11.88 [-20.08, -3.67] | 4.19 | -2.84 | 0.05 |
| Class: Reptilia | 4.60 [0.41, 8.79] | 2.14 | 2.15 | 0.03 |
| Mean $CT_{max}$ : Exposure Duration (Prolonged) | -0.09 [-0.49, 0.31] | 0.2 | -0.43 | 0.67 |
| Species variance | 20.213 | | | |
| *QM test (moderators)* | *225.21 (df = 11)* | | | *<0.0001* |
| *QE test (residual heterogeneity)* | *19994.85 (df = 147)* | | | *<0.0001* |
| *Pseudo R2* | *47.00%* | | | |
| *Residual heterogeneity explained* | *58.50%* | | | |
| *delta AIC (moderated - null)* | *-57.23* | | | |

*Additive:*

| **Term** | **Estimate [95% CI]** | **SE** | **Z statistic** | **P-value** |
|---|---|---|---|---|
| Intercept | 37.97 [34.77, 41.16] | 1.63 | 23.28 | <0.0001 |
| Mean $CT_{max}$ | 0.90 [0.65, 1.15] | 0.13 | 7.13 | <0.0001 |
| Exposure Duration (Prolonged) | -3.72 [-6.38, -1.06] | 1.36 | -2.74 | 0.01 |
| Class: Amphibia | 0.30 [-2.94, 3.55] | 1.66 | 0.18 | 0.85 |

| | | | | |
|---|---|---|---|---|
| Class: Arachnida | -4.16 [-9.38, 1.06] | 2.66 | -1.56 | 0.12 |
| Class: Bivalvia | 3.27 [-5.84, 12.38] | 4.65 | 0.7 | 0.48 |
| Class: Gastropoda | 12.04 [2.90, 21.18] | 4.66 | 2.58 | 0.01 |
| Class: Insecta | -1.35 [-6.41, 3.71] | 2.58 | -0.52 | 0.60 |
| Class: Liliopsida | 1.86 [-6.61, 10.33] | 4.32 | 0.43 | 0.67 |
| Class: Magnoliopsida | -11.24 [-18.87, -3.62] | 3.89 | -2.89 | <0.01 |
| Class: Reptilia | 4.71 [0.57, 8.84] | 2.11 | 2.23 | 0.03 |
| Species variance | 19.957 | | | |
| *QM test (moderators)* | *227.86 (df = 10)* | | | *<0.0001* |
| *QE test (residual heterogeneity)* | *19997.18 (df = 148)* | | | *<0.0001* |
| *Pseudo R2* | *47.70%* | | | |
| *Residual heterogeneity explained* | *58.50%* | | | |
| *delta AIC (moderated - null)* | *-53.61* | | | |

**Table S7. Environmental Realm Hypothesis test.**

Results from a multi-level meta-regression model testing the hypothesis that relationships between thermal performance curve (TPC) traits ($T_{opt}$, $TPC_{max}$) and $CT_{max}$ depend on the environmental realm (aquatic vs terrestrial) of the species sampled. $CT_{max}$ was standardized to a 15 °C acclimation temperature using phylum-specific acclimation response ratios (ARRs) and centered ($T_{opt}$ dataset mean $CT_{max}$ = 37.42°C; $TPC_{max}$ dataset mean $CT_{max}$ = 38.12°C). Class was included as fixed effect covariate to account for taxonomic differences and species was included as a random intercept to account for non-independent sampling of TPCs. Slopes and 95% confidence intervals describe the scaling relationships of TPC traits with $CT_{max}$. For models where interaction terms were non-significant, results are also reported for the additive model. For these models, the aquatic environmental realm is the reference group and Actinopterygii is the reference class.

**$T_{opt}$ (N = 204 TPCs, 102 species)**

*Interactive:*

| **Term** | **Estimate [95% CI]** | **SE** | **Z statistic** | **P-value** |
|---|---|---|---|---|
| Intercept | 24.65 [21.80, 27.51] | 1.46 | 16.92 | <0.0001 |
| Mean $CT_{max}$ | 0.90 [0.65, 1.16] | 0.13 | 6.92 | <0.0001 |
| Realm (Terrestrial) | 0.87 [-2.37, 4.11] | 1.65 | 0.53 | 0.60 |
| Class: Amphibia | 0.98 [-2.86, 4.82] | 1.96 | 0.5 | 0.62 |
| Class: Arachnida | -2.59 [-7.63, 2.45] | 2.57 | -1.01 | 0.31 |
| Class: Bivalvia | 5.96 [-2.77, 14.69] | 4.45 | 1.34 | 0.18 |
| Class: Gastropoda | 15.73 [6.96, 24.50] | 4.48 | 3.51 | <0.001 |
| Class: Insecta | -1.02 [-6.92, 4.89] | 3.01 | -0.34 | 0.74 |
| Class: Liliopsida | -3.69 [-12.52, 5.14] | 4.5 | -0.82 | 0.41 |
| Class: Magnoliopsida | -12.03 [-20.52, -3.54] | 4.33 | -2.78 | 0.01 |
| Class: Reptilia | 6.90 [2.19, 11.60] | 2.4 | 2.87 | <0.01 |
| Mean $CT_{max}$ : Realm (Terrestrial) | -0.25 [-0.67, 0.18] | 0.22 | -1.15 | 0.25 |
| Species variance | 19.01 | | | |
| *QM test (moderators)* | *270.64 (df = 11)* | | | *<0.0001* |
| *QE test (residual heterogeneity)* | *44417.52 (df = 192)* | | | *<0.0001* |
| *Pseudo R2* | *39.30%* | | | |
| *Residual heterogeneity explained* | *43.60%* | | | |
| *delta AIC (moderated - null)* | *-59.08* | | | |

*Additive:*

| **Term** | **Estimate [95% CI]** | **SE** | **Z statistic** | **P-value** |
|---|---|---|---|---|
| Intercept | 23.86 [21.34, 26.38] | 1.29 | 18.55 | <0.0001 |
| Mean $CT_{max}$ | 0.81 [0.61, 1.02] | 0.1 | 7.77 | <0.0001 |
| Realm (Terrestrial) | 1.07 [-2.16, 4.30] | 1.65 | 0.65 | 0.51 |

| | | | | |
|---|---|---|---|---|
| Class: Amphibia | 1.72 [-1.91, 5.35] | 1.85 | 0.93 | 0.35 |
| Class: Arachnida | -1.74 [-6.57, 3.10] | 2.47 | -0.7 | 0.48 |
| Class: Bivalvia | 5.90 [-2.85, 14.65] | 4.46 | 1.32 | 0.19 |
| Class: Gastropoda | 16.04 [7.27, 24.81] | 4.48 | 3.58 | <0.001 |
| Class: Insecta | -1.33 [-7.22, 4.57] | 3.01 | -0.44 | 0.66 |
| Class: Liliopsida | -5.21 [-13.67, 3.24] | 4.31 | -1.21 | 0.23 |
| Class: Magnoliopsida | -14.39 [-21.88, -6.90] | 3.82 | -3.77 | <0.001 |
| Class: Reptilia | 7.04 [2.33, 11.74] | 2.4 | 2.93 | <0.01 |
| Species variance | 19.09 | | | |
| QM test (moderators) | *268.27 (df = 10)* | | | *<0.0001* |
| QE test (residual heterogeneity) | *44835.21 (df = 193)* | | | *<0.0001* |
| Pseudo R2 | *39.10%* | | | |
| Residual heterogeneity explained | *43.10%* | | | |
| delta AIC (moderated - null) | *-54.47* | | | |

**$TPC_{max}$ (N = 159 TPCs, 79 species)**

*Interactive:*

| **Term** | **Estimate [95% CI]** | **SE** | **Z statistic** | **P-value** |
|---|---|---|---|---|
| Intercept | 38.02 [34.10, 41.94] | 2 | 19.02 | <0.0001 |
| Mean $CT_{max}$ | 1.14 [0.76, 1.52] | 0.19 | 5.91 | <0.0001 |
| Realm (Terrestrial) | -0.63 [-4.83, 3.57] | 2.14 | -0.29 | 0.77 |
| Class: Amphibia | -1.07 [-5.89, 3.75] | 2.46 | -0.44 | 0.66 |
| Class: Arachnida | -4.37 [-10.31, 1.57] | 3.03 | -1.44 | 0.15 |
| Class: Bivalvia | 1.92 [-7.55, 11.39] | 4.83 | 0.4 | 0.69 |
| Class: Gastropoda | 13.43 [3.93, 22.92] | 4.85 | 2.77 | 0.01 |
| Class: Insecta | -2.43 [-9.07, 4.21] | 3.39 | -0.72 | 0.47 |
| Class: Liliopsida | 3.86 [-5.93, 13.66] | 5 | 0.77 | 0.44 |
| Class: Magnoliopsida | -8.63 [-17.92, 0.66] | 4.74 | -1.82 | 0.07 |
| Class: Reptilia | 5.55 [-0.32, 11.42] | 3 | 1.85 | 0.06 |
| Mean $CT_{max}$ : Realm (Terrestrial) | -0.35 [-0.86, 0.15] | 0.26 | -1.36 | 0.17 |
| Species variance | 21.91 | | | |
| QM test (moderators) | *202.85 (df = 11)* | | | *<0.0001* |
| QE test (residual heterogeneity) | *23706.21 (df = 147)* | | | *<0.0001* |
| Pseudo R2 | *42.60%* | | | |
| Residual heterogeneity explained | *50.90%* | | | |
| delta AIC (moderated - null) | *-52.14* | | | |

*Additive*:

| **Term** | **Estimate [95% CI]** | **SE** | **Z statistic** | **P-value** |
|---|---|---|---|---|
| Intercept | 36.38 [33.23, 39.53] | 1.61 | 22.64 | <0.0001 |
| Mean $CT_{max}$ | 0.95 [0.69, 1.20] | 0.13 | 7.18 | <0.0001 |
| Realm (Terrestrial) | 0.03 [-4.09, 4.15] | 2.1 | 0.02 | 0.99 |
| Class: Amphibia | 0.20 [-4.29, 4.69] | 2.29 | 0.09 | 0.93 |
| Class: Arachnida | -2.65 [-8.09, 2.79] | 2.77 | -0.96 | 0.34 |
| Class: Bivalvia | 1.61 [-7.91, 11.14] | 4.86 | 0.33 | 0.74 |
| Class: Gastropoda | 13.91 [4.38, 23.44] | 4.86 | 2.86 | <0.01 |
| Class: Insecta | -2.11 [-8.78, 4.56] | 3.4 | -0.62 | 0.53 |
| Class: Liliopsida | 2.80 [-6.94, 12.55] | 4.97 | 0.56 | 0.57 |
| Class: Magnoliopsida | -10.54 [-19.47, -1.60] | 4.56 | -2.31 | 0.02 |
| Class: Reptilia | 6.16 [0.31, 12.00] | 2.98 | 2.07 | 0.04 |
| Species variance | 22.21 | | | |
| *QM test (moderators)* | *198.37 (df = 10)* | | | *<0.0001* |
| *QE test (residual heterogeneity)* | *23936.12 (df = 148)* | | | *<0.0001* |
| *Pseudo R2* | *41.80%* | | | |
| *Residual heterogeneity explained* | *50.40%* | | | |
| *delta AIC (moderated - null)* | *-46.78* | | | |

**Table S8. Climatic realm (temperate vs tropical)**

Results from a multi-level meta-regression model testing the hypothesis that relationships between thermal performance curve (TPC) traits ($T_{opt}$, $TPC_{max}$) and $CT_{max}$ depend on the climatic realm (temperate vs tropical) of the species sampled. $CT_{max}$ was standardized to a 15 °C acclimation temperature using phylum-specific acclimation response ratios (ARRs) and centered ($T_{opt}$ dataset mean $CT_{max}$ = 37.42°C; $TPC_{max}$ dataset mean $CT_{max}$ = 38.12°C). Class was included as fixed effect covariate to account for taxonomic differences and species was included as a random intercept to account for non-independent sampling of TPCs. Slopes and 95% confidence intervals describe the scaling relationships of TPC traits with $CT_{max}$. For models where interaction terms were non-significant, results are also reported for the additive model. For all models, the temperate climate category is the reference group and Actinopterygii is the reference class.

**$T_{opt}$ (N = 204 TPCs, 102 species)**

*Interactive:*

| **Term** | **Estimate [95% CI]** | **SE** | **Z statistic** | **P-value** |
|---|---|---|---|---|
| Intercept | 23.47 [20.86, 26.08] | 1.33 | 17.62 | <0.0001 |
| Mean $CT_{max}$ | 0.78 [0.57, 0.99] | 0.11 | 7.13 | <0.0001 |
| Climate (Tropical) | -0.47 [-0.90, -0.04] | 0.22 | -2.14 | 0.03 |
| Class: Amphibia | 2.85 [0.02, 5.68] | 1.44 | 1.97 | 0.05 |
| Class: Arachnida | -1.05 [-5.91, 3.80] | 2.48 | -0.43 | 0.67 |
| Class: Bivalvia | 5.95 [-2.79, 14.69] | 4.46 | 1.33 | 0.18 |
| Class: Gastropoda | 16.24 [7.47, 25.01] | 4.47 | 3.63 | <0.001 |
| Class: Insecta | -0.06 [-5.01, 4.89] | 2.53 | -0.02 | 0.98 |
| Class: Liliopsida | -3.27 [-11.23, 4.68] | 4.06 | -0.81 | 0.42 |
| Class: Magnoliopsida | -12.26 [-19.27, -5.26] | 3.57 | -3.43 | <0.001 |
| Class: Reptilia | 8.37 [4.91, 11.83] | 1.76 | 4.74 | <0.0001 |
| Mean $CT_{max}$ : Climate (Tropical) | 0.66 [-0.07, 1.39] | 0.37 | 1.77 | 0.08 |
| Species variance | 19.04 | | | |
| *QM test (moderators)* | *310.08 (df = 11)* | | | *<0.0001* |
| *QE test (residual heterogeneity)* | *40025.82 (df = 192)* | | | *<0.0001* |
| *Pseudo R2* | *39.20%* | | | |
| *Residual heterogeneity explained* | *49.20%* | | | |
| *delta AIC (moderated - null)* | *-93.55* | | | |

*Additive:*

| **Term** | **Estimate [95% CI]** | **SE** | **Z statistic** | **P-value** |
|---|---|---|---|---|
| Intercept | 24.10 [21.55, 26.64] | 1.3 | 18.54 | <0.0001 |
| Mean $CT_{max}$ | 0.83 [0.63, 1.04] | 0.11 | 7.87 | <0.0001 |
| Climate (Tropical) | -0.79 [-1.04, -0.54] | 0.13 | -6.2 | <0.0001 |
| Class: Amphibia | 2.60 [-0.25, 5.45] | 1.45 | 1.79 | 0.0736 |

| | | | | |
|---|---|---|---|---|
| Class: Arachnida | -1.80 [-6.64, 3.04] | 2.47 | -0.73 | 0.47 |
| Class: Bivalvia | 5.85 [-2.99, 14.68] | 4.51 | 1.3 | 0.19 |
| Class: Gastropoda | 15.91 [7.04, 24.77] | 4.52 | 3.52 | 0.0004 |
| Class: Insecta | -0.42 [-5.41, 4.57] | 2.55 | -0.17 | 0.87 |
| Class: Liliopsida | -4.63 [-12.54, 3.27] | 4.03 | -1.15 | 0.25 |
| Class: Magnoliopsida | -13.91 [-20.75, -7.07] | 3.49 | -3.99 | <0.0001 |
| Class: Reptilia | 7.95 [4.48, 11.41] | 1.77 | 4.49 | <0.0001 |
| *Species variance* | *19.49* | | | |
| *QM test (moderators)* | *300.84 (df = 10)* | | | *<0.0001* |
| *QE test (residual heterogeneity)* | *40039.97 (df = 193)* | | | *<0.0001* |
| *Pseudo R2* | *37.80%* | | | |
| *Residual heterogeneity explained* | *49.20%* | | | |
| *delta AIC (moderated - null)* | *-87.75* | | | |

**$TPC_{max}$ (N = 159 TPCs, 79 species)**

*Interactive:*

| **Term** | **Estimate [95% CI]** | **SE** | **Z statistic** | **P-value** |
|---|---|---|---|---|
| Intercept | 36.08 [32.83, 39.34] | 1.66 | 21.74 | <0.0001 |
| Mean $CT_{max}$ | 0.92 [0.65, 1.19] | 0.14 | 6.78 | <0.0001 |
| Climate (Tropical) | 0.33 [-0.74, 1.40] | 0.54 | 0.61 | 0.54 |
| Class: Amphibia | 0.43 [-3.02, 3.88] | 1.76 | 0.24 | 0.81 |
| Class: Arachnida | -2.31 [-7.76, 3.14] | 2.78 | -0.83 | 0.41 |
| Class: Bivalvia | 1.66 [-7.83, 11.15] | 4.84 | 0.34 | 0.73 |
| Class: Gastropoda | 14.06 [4.55, 23.57] | 4.85 | 2.9 | <0.01 |
| Class: Insecta | -1.91 [-7.22, 3.40] | 2.71 | -0.7 | 0.48 |
| Class: Liliopsida | 3.44 [-5.59, 12.47] | 4.61 | 0.75 | 0.45 |
| Class: Magnoliopsida | -9.77 [-18.03, -1.50] | 4.22 | -2.32 | 0.02 |
| Class: Reptilia | 6.31 [2.10, 10.53] | 2.15 | 2.94 | <0.01 |
| Mean $CT_{max}$ : Climate (Tropical) | 0.32 [-0.55, 1.19] | 0.44 | 0.71 | 0.48 |
| Species variance | 22.03 | | | |
| *QM test (moderators)* | *200.55 (df = 11)* | | | *<0.0001* |
| *QE test (residual heterogeneity)* | *22987.22 (df = 147)* | | | *<0.0001* |
| *Pseudo R2* | *42.30%* | | | |
| *Residual heterogeneity explained* | *52.30%* | | | |
| *delta AIC (moderated - null)* | *-45.31* | | | |

*Additive:*

| **Term** | **Estimate [95% CI]** | **SE** | **Z statistic** | **P-value** |
|---|---|---|---|---|
| Intercept | 36.39 [33.27, 39.52] | 1.6 | 22.81 | <0.0001 |
| Mean $CT_{max}$ | 0.95 [0.69, 1.20] | 0.13 | 7.24 | <0.0001 |
| Climate (Tropical) | -0.04 [-0.33, 0.24] | 0.15 | -0.3 | 0.76 |

| | | | | |
|---|---|---|---|---|
| Class: Amphibia | 0.23 [-3.17, 3.62] | 1.73 | 0.13 | 0.89 |
| Class: Arachnida | -2.66 [-8.01, 2.68] | 2.73 | -0.98 | 0.33 |
| Class: Bivalvia | 1.61 [-7.84, 11.06] | 4.82 | 0.33 | 0.74 |
| Class: Gastropoda | 13.90 [4.44, 23.36] | 4.83 | 2.88 | <0.01 |
| Class: Insecta | -2.09 [-7.35, 3.18] | 2.69 | -0.78 | 0.44 |
| Class: Liliopsida | 2.81 [-6.02, 11.63] | 4.5 | 0.62 | 0.53 |
| Class: Magnoliopsida | -10.54 [-18.50, -2.58] | 4.06 | -2.6 | <0.01 |
| Class: Reptilia | 6.18 [2.00, 10.36] | 2.13 | 2.9 | <0.01 |
| Species variance | 21.867 | | | |
| *QM test (moderators)* | *201.51 (df = 10)* | | | *<0.0001* |
| *QE test (residual heterogeneity)* | *23644.62 (df = 148)* | | | *<0.0001* |
| *Pseudo R2* | *42.70%* | | | |
| *Residual heterogeneity explained* | *51.00%* | | | |
| *delta AIC (moderated - null)* | *-41.92* | | | |

**Table S9. Relationship between the supra-optimal range of performance (SOR) and $CT_{max}$ across species.**

Results from a multi-level regression model evaluating the relationship between the supra-optimal range (SOR, representing $TPC_{max}$-$T_{opt}$) and $CT_{max}$, accounting for Class effects as a random intercept (N = 136, 68 species). $CT_{max}$ was standardized to a 15°C acclimation temperature using phylum-specific acclimation response ratios (ARRs). Slopes and 95% confidence intervals describe the relationships.

| **Term** | **Estimate [95% CI]** | **SE** | **Z statistic** | **P-value** |
|---|---|---|---|---|
| Intercept | 1.99 [-5.89, 9.87] | 4.02 | 0.49 | 0.62 |
| Mean $CT_{max}$ | 0.26 [0.01, 0.52] | 0.13 | 2.04 | 0.04 |
| Class: Amphibia | -1.94 [-5.18, 1.30] | 1.65 | -1.17 | 0.24 |
| Class: Arachnida | -1.17 [-5.97, 3.64] | 2.45 | -0.48 | 0.63 |
| Class: Bivalvia | -3.03 [-11.14, 5.09] | 4.14 | -0.73 | 0.46 |
| Class: Gastropoda | -1.27 [-9.38, 6.83] | 4.14 | -0.31 | 0.75 |
| Class: Insecta | -5.69 [-11.19, -0.20] | 2.8 | -2.03 | 0.04 |
| Class: Liliopsida | 4.46 [-3.64, 12.55] | 4.13 | 1.08 | 0.28 |
| Class: Magnoliopsida | 0.58 [-7.34, 8.49] | 4.04 | 0.14 | 0.89 |
| Class: Reptilia | -2.12 [-6.03, 1.79] | 2 | -1.06 | 0.29 |
| Species variance | 15.76 | | | |
| *QM test (moderators)* | *38.16 (df = 9)* | | | *<0.0001* |
| *QE test (residual heterogeneity)* | *11369.77 (df = 126)* | | | *<0.0001* |
| *Pseudo R2* | *5.40%* | | | |
| *Residual heterogeneity explained* | *18.00%* | | | |
| *delta AIC (moderated - null)* | *-7.39* | | | |

**Table S10. Joint analysis of $T_{opt}$ and $TPC_{max}$ relationships with $CT_{max}$ using a trait interaction model.**

Results from a multi-level meta-regression jointly modeling thermal performance curve (TPC) traits ($T_{opt}$ and $TPC_{max}$) to test whether their relationships with $CT_{max}$ differ. Trait type ($T_{opt}$ or $TPC_{max}$), $CT_{max}$, their interaction, and class were included as fixed effects, with species included as a random intercept to account for non-independent sampling of paired trait estimates. $CT_{max}$ was standardized to a 15°C acclimation temperature using phylum-specific acclimation response ratios (ARRs). The interaction between trait type and $CT_{max}$ tests whether $T_{opt}$ and $TPC_{max}$ exhibit different scaling relationships with $CT_{max}$. The null model used for model comparison included trait type and class as fixed effects and species as a random intercept (N = 136 TPCs from 68 species). Slopes and 95% confidence intervals describe the relationships between each TPC trait and $CT_{max}$, with $T_{opt}$ and Actinopterygii serving as the reference categories.

| **Term** | **Estimate [95% CI]** | **SE** | **Z statistic** | **P-value** |
|---|---|---|---|---|
| Intercept | 0.66 [-7.02, 8.33] | 3.92 | 0.17 | 0.8665 |
| Mean $CT_{max}$ | 0.68 [0.43, 0.93] | 0.13 | 5.39 | <0.0001 |
| TPC trait type : mean $CT_{max}$ | -1.65 [-1.87, -1.42] | 0.11 | -14.55 | <0.0001 |
| Class: Amphibia | -0.61 [-3.76, 2.54] | 1.61 | -0.38 | 0.70 |
| Class: Arachnida | -3.39 [-8.09, 1.31] | 2.4 | -1.42 | 0.16 |
| Class: Bivalvia | 2.17 [-5.78, 10.11] | 4.06 | 0.53 | 0.59 |
| Class: Gastropoda | 13.27 [5.31, 21.22] | 4.06 | 3.27 | <0.01 |
| Class: Insecta | -2.11 [-7.45, 3.23] | 2.72 | -0.78 | 0.44 |
| Class: Liliopsida | -2.36 [-10.16, 5.43] | 3.98 | -0.59 | 0.55 |
| Class: Magnoliopsida | -11.62 [-19.33, -3.90] | 3.94 | -2.95 | <0.01 |
| Class: Reptilia | 6.26 [2.46, 10.06] | 1.94 | 3.23 | <0.01 |
| TPC trait type : mean $CT_{max}$ (mean $TPC_{max}$) | 0.31 [0.30, 0.32] | 0.003 | 94.54 | <0.0001 |
| Species variance | 15.21 | | | |
| *QM test (moderators)* | *105674.08 (df = 11)* | | | *<0.0001* |
| *QE test (residual heterogeneity)* | *110939.98 (df = 260)* | | | *<0.0001* |
| *Pseudo R2* | *40.20%* | | | |
| *Residual heterogeneity explained* | *48.40%* | | | |
| *delta AIC (moderated - null)* | *-8970.32* | | | |

**Table S11. Unstandardized and standardized $CT_{max}$ correlation coefficients across taxonomic summaries**.

Correlation coefficients among species' $CT_{max}$ values that were not ARR-standardized ("None") and those standardized using ARR means of different taxonomic hierarchies.

| | None | Family | Order | Class | Phylum | Kingdom |
|---|---|---|---|---|---|---|
| **None** | 1 | | | | | |
| **Family** | 0.88 | 1 | | | | |
| **Order** | 0.89 | 1 | 1 | | | |
| **Class** | 0.89 | 1 | 1 | 1 | | |
| **Phylum** | 0.88 | 0.99 | 1 | 1 | 1 | |
| **Kingdom** | 0.88 | 0.99 | 0.89 | 1 | 1 | 1 |

**Table S12. Model comparison of alternative random-intercept structures accounting for phylogenetic relatedness.**

We compared multilevel meta-regressions (rma) models that included different hierarchical groupings as random intercepts to evaluate how phylogenetic history influences variance structure. We selected Class as the preferred random-intercept level because it provided a strong balance between model fit and parsimony (AIC), while minimizing categories represented by a single species, which can inflate variance estimates.

**$T_{opt}$:**

| Model | df | AIC | Categories represented by a single species |
|---|---|---|---|
| ~ 1 | 1 | 225025.3 | NA |
| ~ 1 + (1\|Kingdom) | 2 | 220495.8 | 0 |
| ~ 1 + (1\|Class) | 2 | 78838.8 | 1 |
| ~ 1 + (1\|Order) | 2 | 43214.1 | 7 |
| ~ 1 + (1\|Family) | 2 | 8844.3 | 15 |

**$TPC_{max}$:**

| Model | df | AIC | Categories represented by a single species |
|---|---|---|---|
| ~ 1 | 1 | 216598.7 | NA |
| ~ 1 + (1\|Kingdom) | 2 | 136800.4 | 0 |
| ~ 1 + (1\|Class) | 2 | 48346.6 | 1 |
| ~ 1 + (1\|Order) | 2 | 19261.1 | 6 |
| ~ 1 + (1\|Family) | 2 | 10939.9 | 10 |

**Table S13. Results from a multi-level regression model evaluating the log-log scaling relationship between $T_{opt}$ and $CT_{max}$ expressed in Kelvin (N = 204 TPCs, 102 species).**

Temperatures were converted from °C to Kelvin prior to log transformation. $CT_{max}$ was standardized to a 15°C acclimation temperature using phylum-specific acclimation response ratios (ARRs). Class was included as a fixed-effect covariate to account for taxonomic differences, and species was included as a random intercept to account for non-independent sampling of TPCs. Slopes and 95% confidence intervals describe the scaling relationship between log-transformed $T_{opt}$ and log-transformed $CT_{max}$. Here, a slope estimate and CIs overlapping 1 indicates proportional scaling.

| **Term** | **Estimate [95% CI]** | **SE** | **Z statistic** | **P-value** |
|---|---|---|---|---|
| Intercept | 5.70 [5.69, 5.71] | 0 | 1298.15 | <0.0001 |
| Log Mean $CT_{max}$ | 0.71 [0.46, 0.95] | 0.13 | 5.57 | <0.0001 |
| Class: Amphibia | 0.00 [-0.01, 0.02] | 0.01 | 0.93 | 0.35 |
| Class: Arachnida | -0.01 [-0.02, 0.01] | 0.01 | -1.07 | 0.28 |
| Class: Bivalvia | 0.01 [-0.01, 0.04] | 0.01 | 0.89 | 0.37 |
| Class: Gastropoda | 0.05 [0.02, 0.07] | 0.01 | 4.26 | <0.0001 |
| Class: Insecta | 0.00 [-0.01, 0.02] | 0.01 | 0.58 | 0.56 |
| Class: Liliopsida | 0.03 [0.00, 0.05] | 0.01 | 1.98 | 0.05 |
| Class: Magnoliopsida | -0.04 [-0.06, -0.01] | 0.01 | -3.06 | <0.01 |
| Class: Reptilia | 0.02 [0.01, 0.04] | 0.01 | 4.19 | <0.0001 |
| Species variance | 0.0001 | | | |
| *QM test (moderators)* | *185.97 (df = 9)* | | | *<0.0001* |
| *QE test (residual heterogeneity)* | *246.63 (df = 194)* | | | *0.0063* |
| *Pseudo R2* | *38.70%* | | | |
| *Residual heterogeneity explained* | *53.10%* | | | |
| *delta AIC (moderated - null)* | *-18.27* | | | |

**Table S14. Results from a multi-level regression model evaluating the log-log scaling relationship between $TPC_{max}$ and $CT_{max}$ expressed in Kelvin (N = 159 TPCs, 79 species).**

Temperatures were converted from °C to Kelvin prior to log transformation. $CT_{max}$ was standardized to a 15°C acclimation temperature using phylum-specific acclimation response ratios (ARRs). Class was included as a fixed-effect covariate to account for taxonomic differences, and species was included as a random intercept to account for non-independent sampling of TPCs. Slopes and 95% confidence intervals describe the scaling relationship between log-transformed $T_{opt}$ and log-transformed $CT_{max}$. Here, a slope estimate and CIs overlapping 1 indicates proportional scaling.

| Term | Estimate [95% CI] | SE | Z statistic | P-value |
|---|---|---|---|---|
| Intercept | 5.74 [5.73, 5.75] | 0.01 | 1084.76 | <0.0001 |
| Log Mean $CT_{max}$ | 0.95 [0.69, 1.22] | 0.14 | 6.98 | <0.0001 |
| Class: Amphibia | -0.00 [-0.02, 0.01] | 0.01 | -0.71 | 0.48 |
| Class: Arachnida | -0.01 [-0.03, 0.00] | 0.01 | -1.36 | 0.17 |
| Class: Bivalvia | 0.00 [-0.03, 0.03] | 0.01 | 0.17 | 0.86 |
| Class: Gastropoda | 0.04 [0.02, 0.07] | 0.01 | 3.28 | <0.01 |
| Class: Insecta | -0.01 [-0.03, 0.01] | 0.01 | -1.15 | 0.25 |
| Class: Liliopsida | 0.01 [-0.02, 0.04] | 0.02 | 0.85 | 0.39 |
| Class: Magnoliopsida | -0.03 [-0.06, -0.01] | 0.01 | -2.45 | 0.01 |
| Class: Reptilia | 0.01 [-0.00, 0.03] | 0.01 | 1.79 | 0.07 |
| Species variance | 0.0001 | | | |
| *QM test (moderators)* | *185.40 (df = 9)* | | | *<0.0001* |
| *QE test (residual heterogeneity)* | *344.68 (df = 149)* | | | *<0.0001* |
| *Pseudo R2* | *51.80%* | | | |
| *Residual heterogeneity explained* | *52.40%* | | | |
| *delta AIC (moderated - null)* | *-29.04* | | | |

**Table S15. Results of a profile-likelihood piecewise regression testing for a breakpoint in the $T_{opt}$–$CT_{max}$ relationship.**

We conducted profile-likelihood grid searches across candidate breakpoints (ψ), evaluating values from 15 to 65 °C in 0.5 °C increments for $T_{opt}$ and from 21.05 to 65 °C in 0.1 °C increments for $TPC_{max}$. We fitted meta-regression models using the same variance weighting and model structure as the primary analyses, including taxonomic Class as a fixed effect and species as a random intercept. Reported results correspond to the breakpoint that maximized model log-likelihood. The pre-ψ slope represents the $CT_{max}$ slope below the estimated breakpoint, and the post-ψ slope change represents the difference in slope above versus below that breakpoint. For the $T_{opt}$–$CT_{max}$ relationship, only one observation exceeded the estimated breakpoint (ψ = 59.5 °C), and the slope change was not significant. For the $TPC_{max}$–$CT_{max}$ relationship, the estimated slope change was significant; however, only 20 observations representing seven species occurred below the selected breakpoint (ψ = 27.05 °C), limiting inference about the relationship in this portion of the $CT_{max}$ range.

| **Model** | **N** | **ψ** | **Pre-ψ slope** | **Post-ψ slope change** | **z** | **p** |
|---|---|---|---|---|---|---|
| $T_{opt}$ ~ $CT_{max}$ seg1 + $CT_{max}$ seg2 + class + (1\|species) | 204 | 59.5 | 0.90 [0.70, 1.1] | –0.22 [–1.11, 0.65] | −0.50 | 0.62 |
| $TPC_{max}$ ~ $CT_{max}$ seg1 + $CT_{max}$ seg2 + class + (1\|species) | 159 | 27.05 | 0.10 [–1.01, 1.21] | 1.04 [0.77, 1.31] | 7.65 | < 0.001 |

### Part C: Supplementary Figures

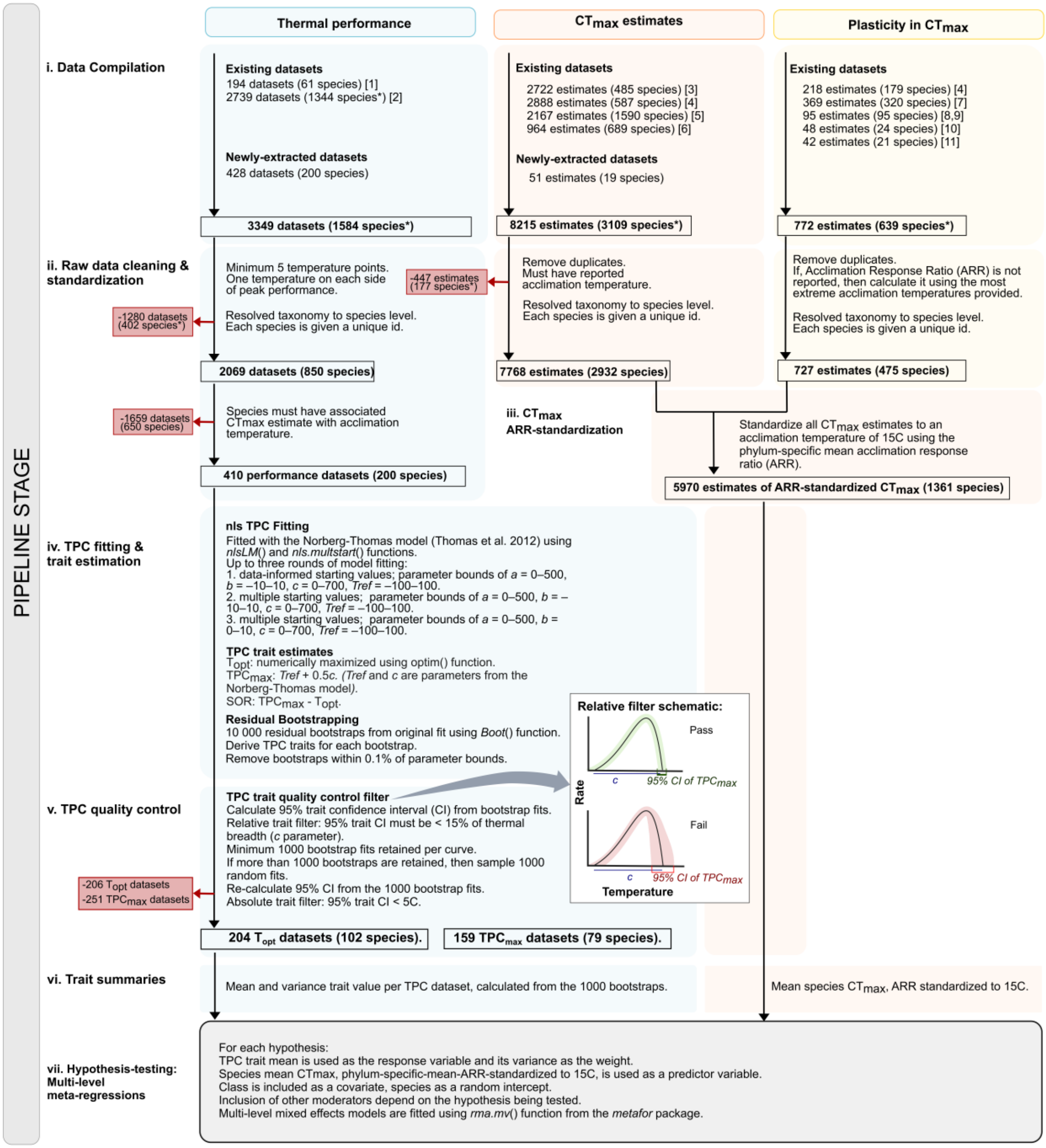

**Figure S1. Data pipeline from data synthesis and compilation to hypothesis-testing models.** We compiled data (i) on thermal performance, $CT_{max}$ estimates, and plasticity in $CT_{max}$ (acclimation response ratio, ARR) from published sources and through literature searches and data extractions. Initial data cleaning (ii) included resolving taxonomy from kingdom to species level for all species units and ensuring necessary overlap between performance data and $CT_{max}$ estimates, as well as $CT_{max}$ estimates and plasticity data via acclimation temperatures. Species

$CT_{max}$ values were standardized (iii) to an acclimation temperature of 15°C using phylum-level average ARRs. Performance data underwent thermal performance curve (TPC) fitting (iv) via the Norberg-Thomas model (21), from which we estimated TPC traits, including temperature of peak performance ($T_{opt}$), maximum temperature of performance above zero ($TPC_{max}$), and the sura-optimal range of performance (SOR, not shown as this trait was not used in hypothesis-testing). Uncertainty in trait estimates was quantified via residual bootstrapping. We then imposed strict quality control of fits (v) using both relative and absolute thresholds of trait confidence interval (CI) width. TPC traits were summarized by dataset as means and variances based on 1000 bootstrap parameter estimates that passed quality control, and standardized $CT_{max}$ was summarized to species means (vi). Our hypothesis-testing multi-level mixed effects models (vii) used mean TPC traits as response variables, weighted by their variance, and species mean standardized $CT_{max}$ as predictor variables. Class was included as a fixed effect and species was included as a random intercept across all models, while additional covariates varied depending on the hypothesis being tested.
Species* indicates species units with unresolved taxonomy. Red arrows to red text show the number of datasets filtered out of our pipeline. References from data compilation: [1] (4), [2] (3), [3] (8), [4] (10), [5] (11), [6] (9), [7] (16), [8] (29), [9] (17), [10] (19), [11] (18).

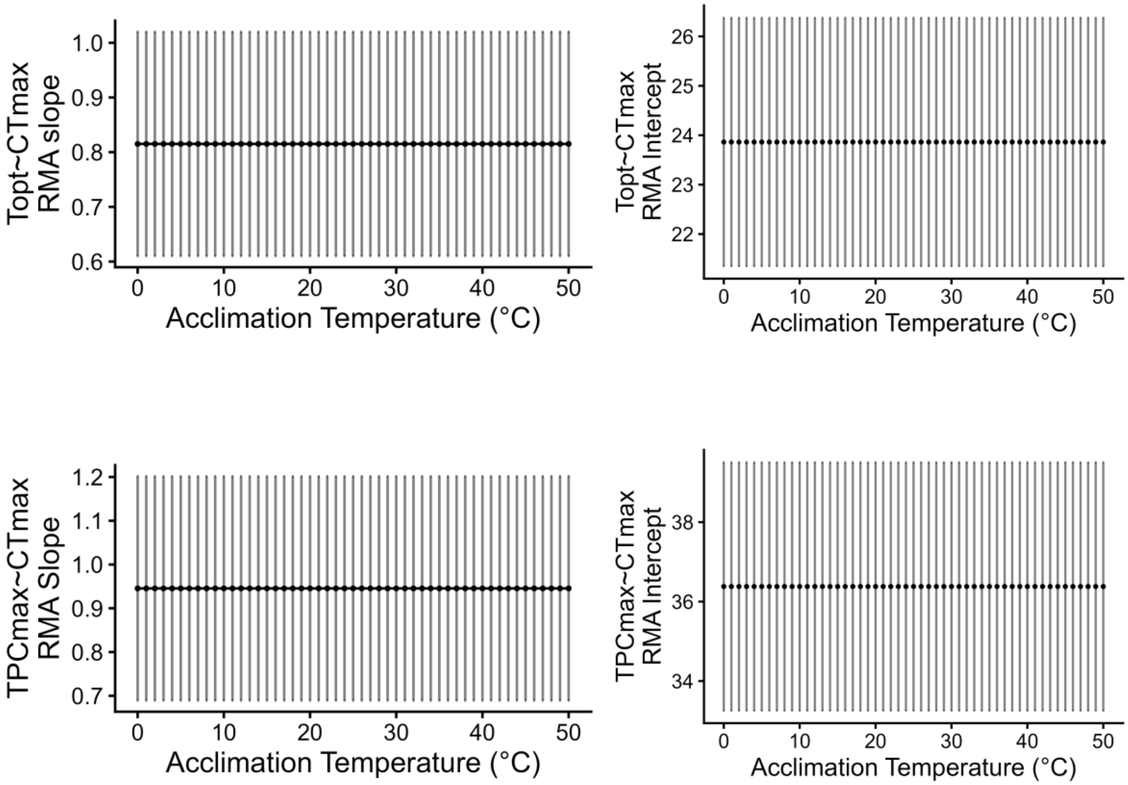

**Figure S2. Sensitivity of TPC-$CT_{max}$ relationships to acclimation temperature choice.** We evaluated the sensitivity of TPC–$CT_{max}$ relationships to the acclimation temperature used to standardize $CT_{max}$ (i.e., *n* in Equation S2) by first using phylum-averaged acclimation response ratios (ARR) to standardize $CT_{max}$ values to a range of possible acclimation temperatures (0–50°C, in 1°C increments). For each possible acclimation temperature, ARR-standardized $CT_{max}$ values were centered on the mean and used as a fixed effect in a meta-regression model with Class as a fixed effect, species as a random intercept, and either $T_{opt}$ (top) or $TPC_{max}$ (bottom) as the response variable.

For both the top ($T_{opt}$) and bottom ($TPC_{max}$) panels, the figure on the left shows the sensitivity of estimated slope coefficients to the acclimation temperature used to standardize $CT_{max}$, showing the intercept (expected TPC trait at the mean scaled standardized $CT_{max}$). Panel on the right show the sensitivity of the regression intercept coefficient to the standardization temperature. Points represent coefficient estimates from each meta-regression, with error bars indicating 95% confidence intervals.

For both traits ($T_{opt}$ and $TPC_{max}$), slope and intercept estimates showed complete overlap in their 95% confidence intervals across the range of acclimation temperatures, indicating that inferred TPC–$CT_{max}$ relationship was largely insensitive to the acclimation temperature used for $CT_{max}$ ARR standardization.

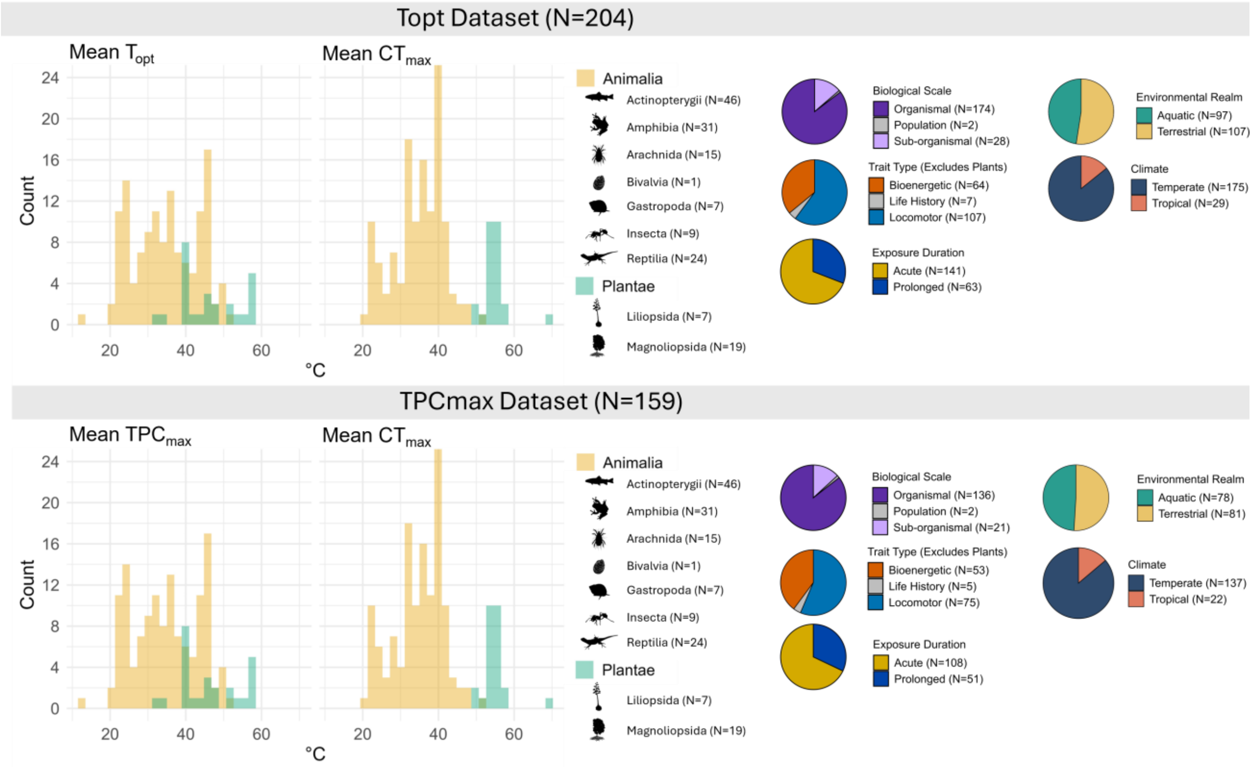

**Figure S3. Data summaries across Classes and TPC trait categories.**

The upper and lower panels summarize the thermal optimum, $T_{opt}$ (N = 204 TPCs) and upper limit of performance, $TPC_{max}$ (N = 159 TPCs) datasets, respectively. Histograms show the distributions of thermal trait estimates and the corresponding mean $CT_{max}$ values for animals (yellow) and plants (green). Taxonomic symbols identify the classes represented and their sample sizes. Pie charts summarize observations by biological scale, trait response category, thermal exposure duration, environmental realm, and climatic zone; trait response category summaries exclude plants. All sample sizes represent TPC-level observations. $CT_{max}$ is the species mean estimate, standardized to ana acclimation temperature of 15°C using phylum-specific acclimation response ratios.

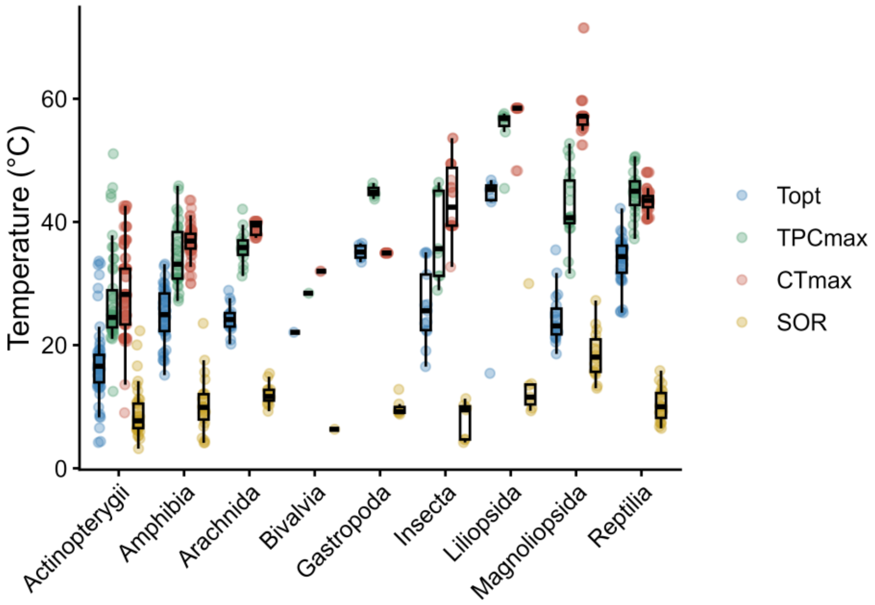

**Figure S4. Distributions of the four thermal traits used in the analyses ($T_{opt}$, $TPC_{max}$, $CT_{max}$, and SOR) across taxonomic classes.**

Colored points represent individual observations, and black boxplots summarize the distribution of each trait within each class; boxes show the interquartile range, and the central line denotes the median. $CT_{max}$ was ARR-standardized to 15℃.

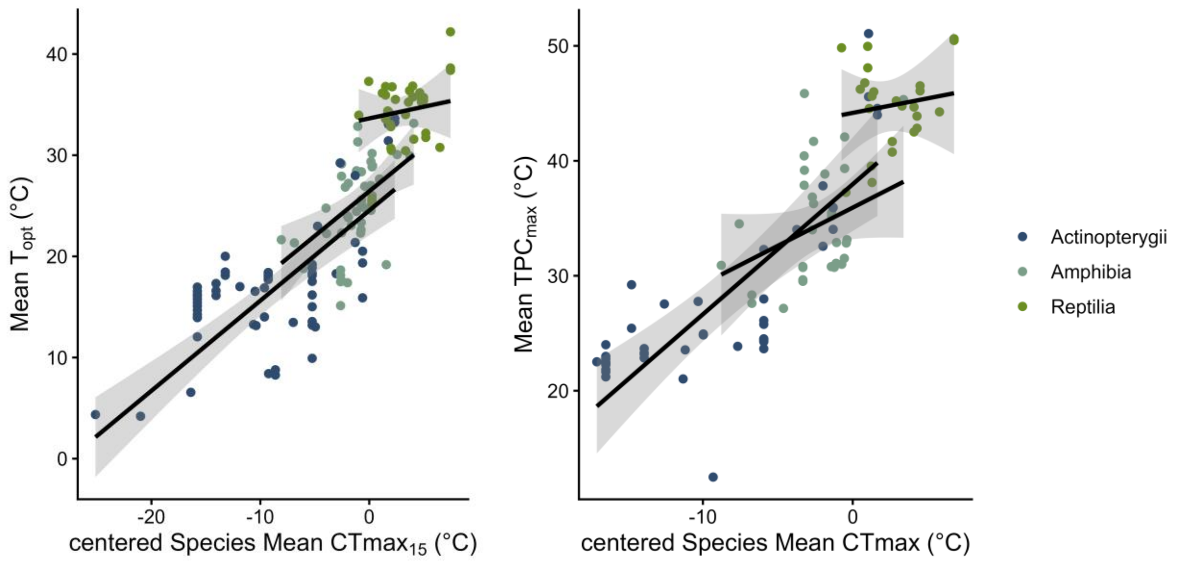

**Figure S5. Class-specific relationships between thermal performance curve traits and critical thermal maximum ($CT_{max}$) in the three best-represented vertebrate classes.**

Points represent trait estimates from individual thermal performance curves for Actinopterygii (blue), Amphibia (teal), and Reptilia (green). Lines and shaded bands show class-specific predictions and 95% confidence intervals from models containing class-by-$CT_{max}$ interactions, with species included as a random effect. Panels show relationships of $CT_{max}$ with (A) thermal optimum ($T_{opt}$) and (B) the upper limit of performance ($TPC_{max}$). The class-by-$CT_{max}$ interactions were not significant, providing no evidence that scaling slopes differed among classes. Species-mean $CT_{max}$ values, standardized to an acclimation temperature of 15°C, were centered separately within each dataset; zero on the x-axis corresponds to $CT_{max}$ = 37.42°C for $T_{opt}$ and 38.12°C for $TPC_{max}$. Only these three classes had sufficient representation to evaluate class-specific scaling relationships.

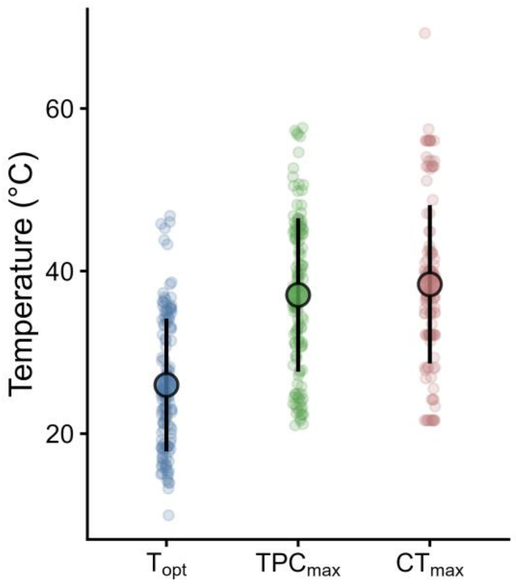

**Figure S6. Distribution and variability of thermal traits in the paired-trait dataset.** $T_{opt}$, $TPC_{max}$, and $CT_{max}$ varied across the observations, with $TPC_{max}$ and $CT_{max}$ generally occurring at higher temperatures than $T_{opt}$. Points show individual TPC-level observations, large outlined points show means, and vertical bars show ±1 SD. The dataset comprised 136 thermal performance curves from 68 species for which $T_{opt}$ and $TPC_{max}$ were estimated from the same fitted curve and a corresponding species-mean $CT_{max}$ was available. $CT_{max}$ values were standardized to an acclimation temperature of 15°C. $T_{opt}$, thermal optimum; $TPC_{max}$, upper limit of performance; $CT_{max}$, critical thermal maximum.

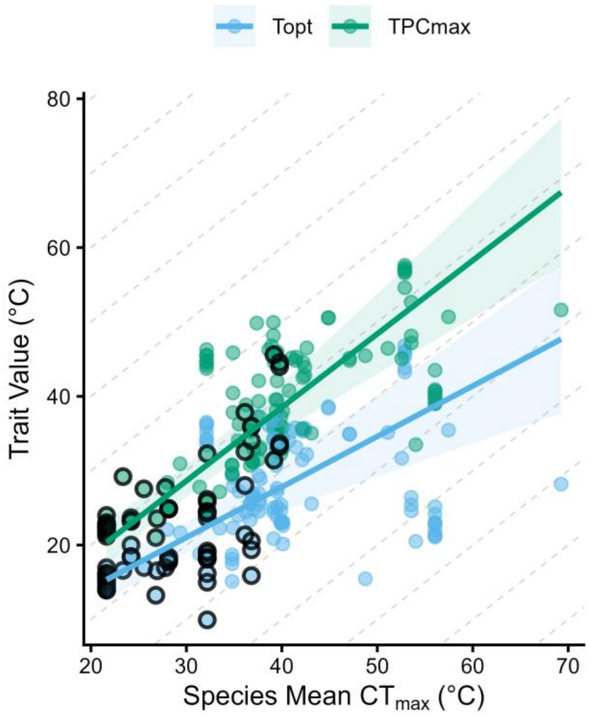

**Figure S7. Scaling relationships between critical thermal maximum ($CT_{max}$) and thermal performance curve traits estimated jointly.**

Each fitted thermal performance curve contributes estimates of thermal optimum ($T_{opt}$, blue) and the upper limit of performance ($TPC_{max}$, green). Points outlined in black represent observations from the reference taxonomic group, Actinopterygii; unoutlined points represent the other classes included in the model. Lines show model-predicted relationships for Actinopterygii. The multi-variate meta-regression jointly modeled $T_{opt}$ and $TPC_{max}$, allowing their relationship with $CT_{max}$ to be compared directly while accounting for taxonomic class, repeated observations within species, and the sampling variance and covariance of both trait estimates. $TPC_{max}$ increased more steeply with $CT_{max}$ than did $T_{opt}$ (slopes = 0.99 and 0.68, respectively; $CT_{max}$-by-trait type interaction: $p < 0.001$). Consequently, the absolute difference between the two traits (the supra-optimal range) widened with $CT_{max}$ by about 0.31°C for every 1°C increase in $CT_{max}$.

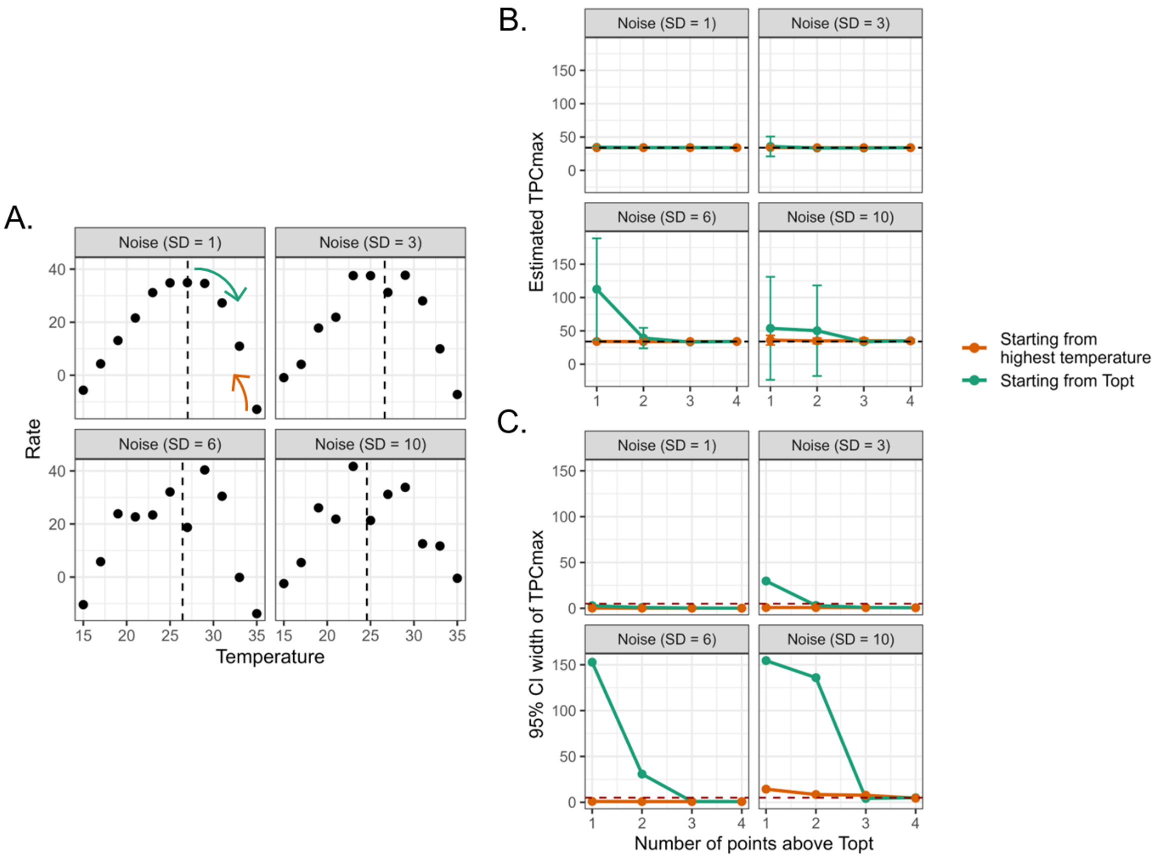

**Figure S8. Model-generated simulation results indicate that the number of observations above $T_{opt}$ influences the precision and accuracy of $TPC_{max}$ estimates.**

We generated four datasets from one Norberg-Thomas model parameter set, each with increasing levels of observation noise (SD = 1, 3, 6, 10). (A) Simulated thermal performance curves under each noise level. The dashed line indicates $T_{opt}$ derived from the generating parameter set. Each dataset was subsequently subsampled to vary both the number and position of sampled temperatures between $T_{opt}$ and the maximum temperature sampled, by including point either starting from $T_{opt}$ (green) or from the highest temperatures (orange). (B) $TPC_{max}$ estimates (points) from 1,000 bootstrap refits with 95% confidence intervals (error bars), shown across noise levels (panels), sampling strategy (colors), and sample sizes (i.e. number of points above $T_{opt}$). Increasing the number of sampled points generally reduces uncertainty in $TPC_{max}$. Sampling temperatures close to $TPC_{max}$ (orange) yields more accurate estimates, with confidence intervals overlapping the true $TPC_{max}$ (dashed black line). (C) Width of the 95% CI across noise levels (panels), sampling strategy (colors), and sample sizes above $T_{opt}$. The dashed red line indicates the most stringent threshold used in our quality control pipeline; estimates exceeding this threshold would be excluded from downstream analyses due to high uncertainty.

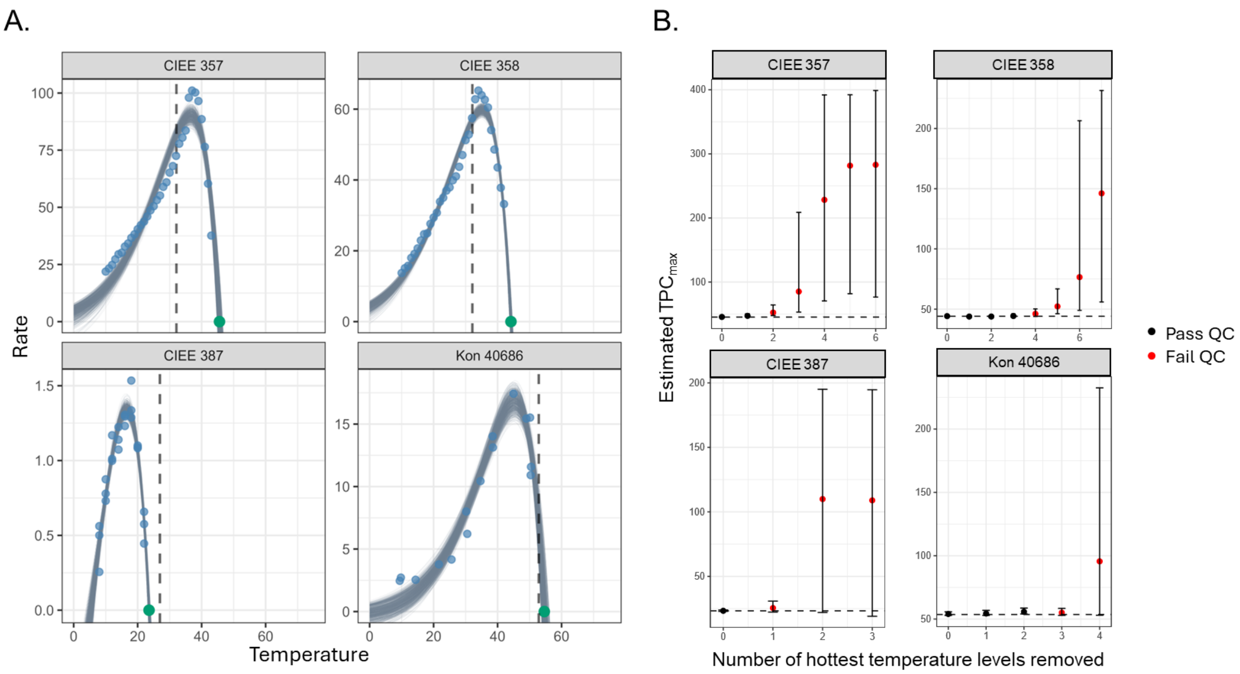

**Figure S9. Sensitivity of TPC$_{max}$ estimates to sampling above T$_{opt}$.**

For four well-sampled thermal performance curves (CIEE 357, CIEE 358, CIEE 387, and Kon 40688), we systematically excluded the hottest temperature observation and refit the Norberg-Thomas model to evaluate how sampling density beyond T$_{opt}$ affects TPC$_{max}$ estimation. A) Empirical observations and TPC fits of the included datasets. Blue points denote raw observed data. Light blue lines denote the TPC bootstrap fits where each line is one of 1000 bootstraps. The black dashed line indicates species CT$_{max}$ and the green point indicates TPC-derived estimated TPC$_{max}$. B) Results of resampled fits, estimates, and quality control filters demonstrating that, across all datasets, increasingly reduced datasets lead to increased uncertainty in TPC$_{max}$ estimates. Importantly, these results also demonstrate the effectiveness of our strict quality control filters in identifying and excluding low-confidence estimates from poor fits. Points represent mean TPC$_{max}$ from 1000 residual bootstrap fits and error bars represent 95% confidence intervals. The black dashed line denotes the TPC$_{max}$ estimated using the complete dataset. Red points indicate that the resampled curve failed our quality control filters and would have been excluded from downstream analyses; black points indicate that the resampled curve would have passed our quality control filters and been included in downstream analyses.

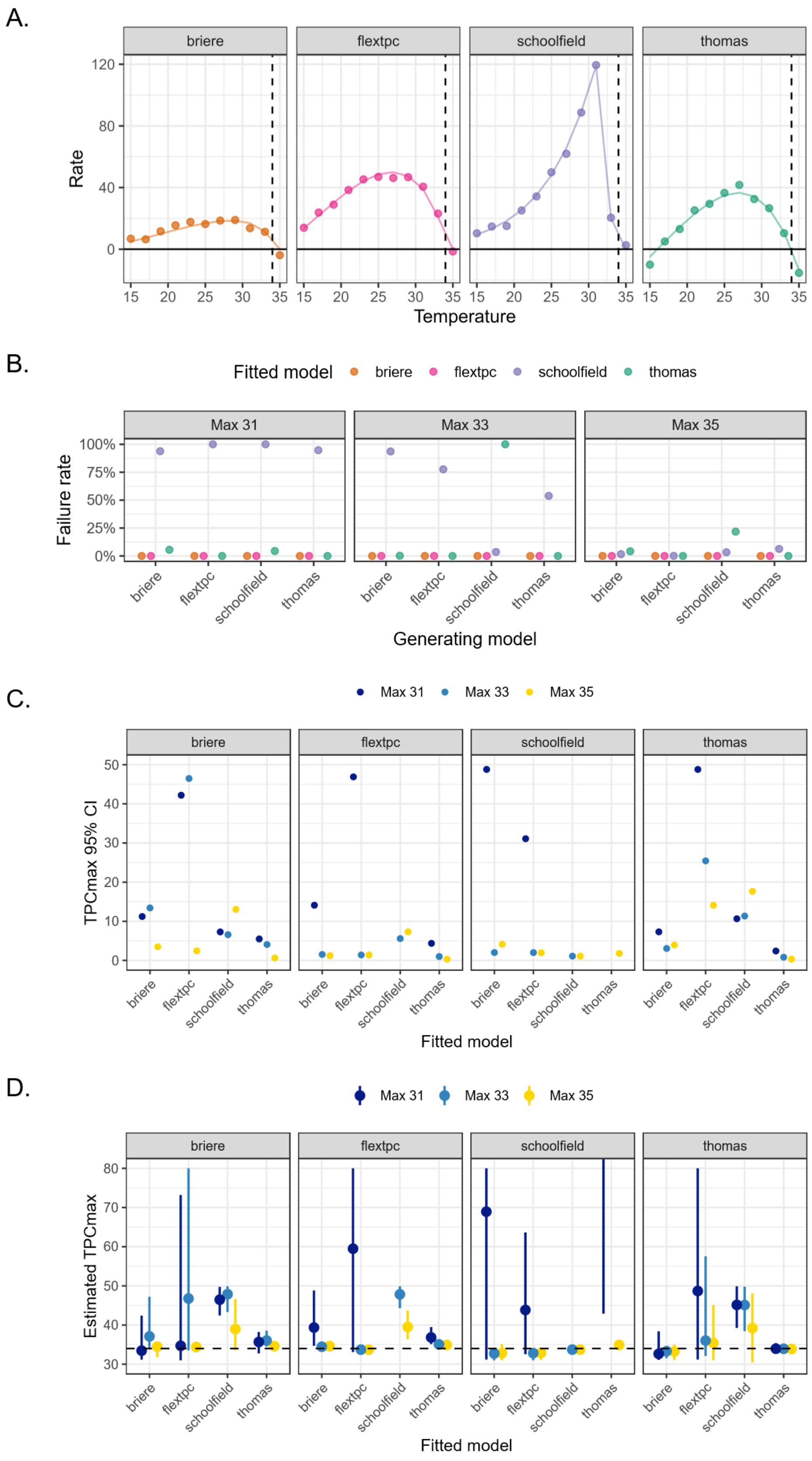

A.
briere
flextpc
schoolfield
thomas
Rate
120
80
40
0
15
20
25
30
35
Temperature
B.
Fitted model
briere
flextpc
schoolfield
thomas
Max 31
Max 33
Max 35
Failure rate
100%
75%
50%
25%
0%
Generating model
C.
Max 31
Max 33
Max 35
TPCmax 95% CI
50
40
30
20
10
0
Fitted model
D.
Max 31
Max 33
Max 35
Estimated TPCmax
80
70
60
50
40
30
Fitted model

**Figure S10. Effects of TPC fitting model on $TPC_{max}$ estimates using model-generated data.** Results from our model-generated simulation analysis demonstrating the effect of TPC model choice on estimation of $TPC_{max}$ under different data sampling densities above $T_{opt}$. (A) Thermal performance curves generated using the Brière II, flexTPC, Sharpe-Schoolfield, and Norberg-Thomas models with known parameters. Points show simulated datasets generated from each curve with Gaussian error (SD = 2). Dashed vertical lines indicate the true $TPC_{max}$ used for each generating model, which was approximately 34°C for all models. (B) Proportion of model fits that failed to converge during fitting for each generating model, fitted model, and sampling scenario, where the maximum sampled temperature above $T_{opt}$ was 31, 33, or 35°C. (C) Width of the 95% confidence interval (CI) for $TPC_{max}$ estimated from 1,000 residual bootstrap fits, showing how uncertainty changed as high-temperature sampling was reduced. (D) Estimated $TPC_{max}$ values across all generating models and sampling scenarios, with 95% CIs from residual bootstrapping. Horizontal dashed lines indicate the true $TPC_{max}$ for each generating model. Missing points indicate model convergence failures. The y-axis in panel D is truncated at 80°C to show variation among biologically interpretable estimates; one poorly constrained Norberg-Thomas estimate from the Schoolfield-generated, Max 31 scenario exceeded this range.

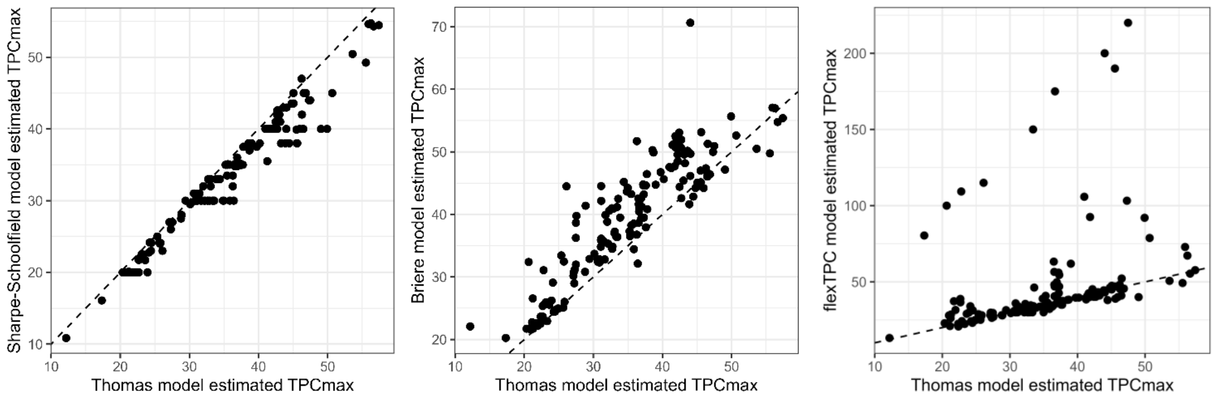

**Figure S11. Comparison of $TPC_{max}$ estimates across thermal performance curve models.** $TPC_{max}$ values estimated from different model fits (Brière II, Sharpe–Schoolfield, and flexTPC) are shown as pairwise comparisons with estimates derived from Norberg-Thomas fitted TPCs. Each point represents a dataset $TPC_{max}$ estimate under two models. The dashed line indicates the 1:1 relationship.

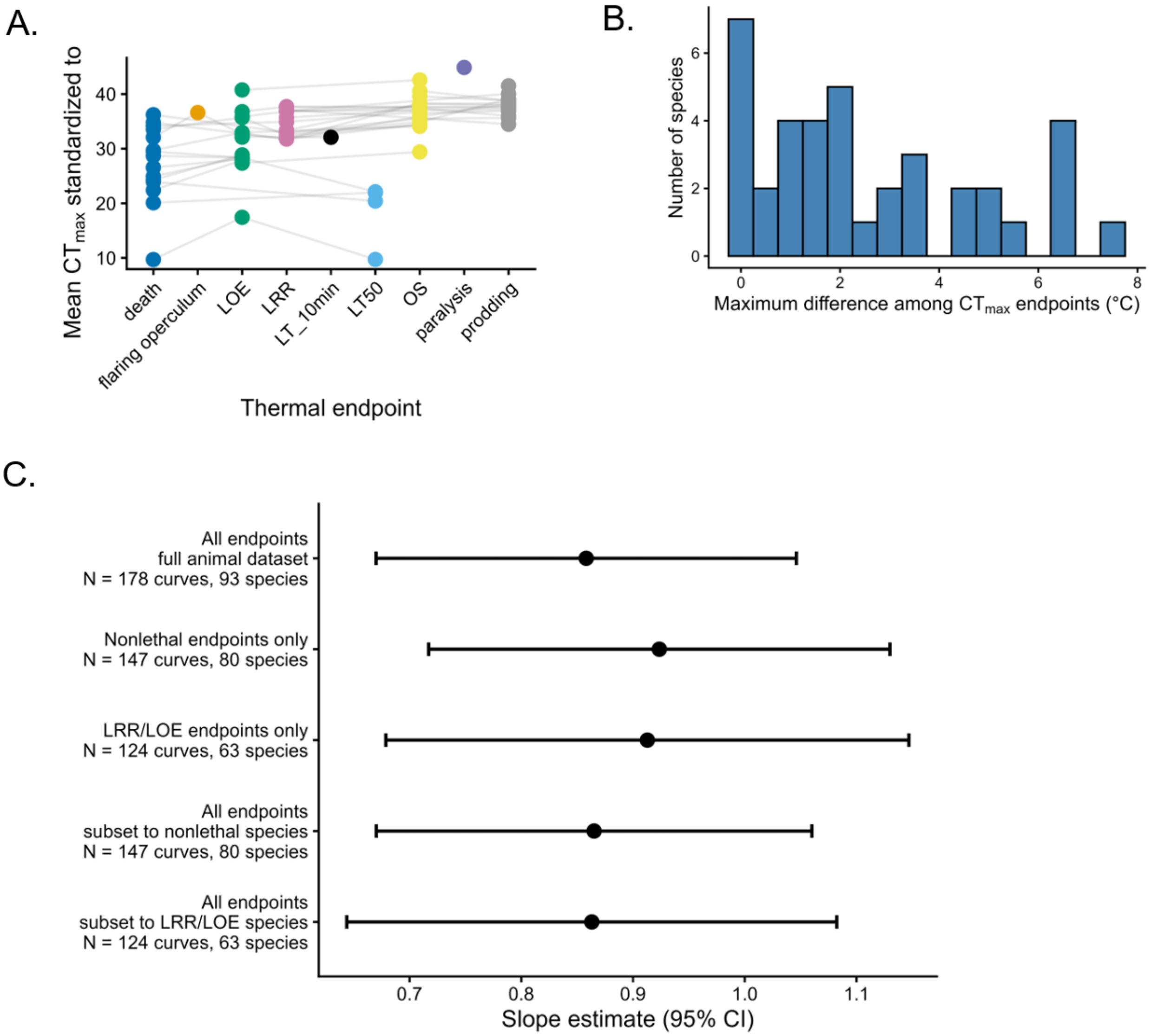

**Figure S12. Sensitivity of $CT_{max}$ to thermal endpoint definitions.**

(A) Mean $CT_{max}$ standardized to 15°C using phylum-level acclimation response ratios, estimated separately for each thermal endpoint definition in species with measurements from multiple endpoints. Lines connect estimates from the same species to illustrate within-species differences among endpoint definitions. (B) Distribution of the maximum within-species difference in mean standardized $CT_{max}$ among endpoint definitions, calculated as the range in mean $CT_{max}$ across all endpoint types reported for each species. (C) Comparison of meta-regression slope estimates relating $T_{opt}$ to $CT_{max}$ under alternative endpoint treatments. Models compare the full animal dataset using all endpoints, datasets restricted to non-lethal endpoints or LRR/LOE endpoints, and all-endpoint models restricted to the same species represented in those endpoint subsets. Points show estimated slopes and error bars denote 95% confidence intervals. Across analyses, endpoint treatment had little influence on the estimated $T_{opt}$-$CT_{max}$ relationship.

#### Part D: TPC Fits and trait estimates

**i. $T_{opt}$ dataset**

Below are the bootstrapped thermal performance curves (TPCs) for each empirical dataset (facets), fitted using the Norberg-Thomas model and used to estimate the temperature of peak performance ($T_{opt}$; N = 204). Thin blue-grey lines represent individual bootstrap fits of parameters (N = 1000) used to summarize $T_{opt}$ and its variance for use in hypothesis-testing multi-level regression models. Empirical observations of performance are shown as small blue points. The estimated $T_{opt}$ are summarized by large blue points (mean) and horizontal bars indicating 95% confidence intervals. These points are placed on the x-axis. Dark grey dashed lines denote the mean species $CT_{max}$, ARR-corrected to a standardized temperature of 15°C. Curves are plotted across a standardized temperature range (0-75°C), and panels are scaled independently along the y-axis.

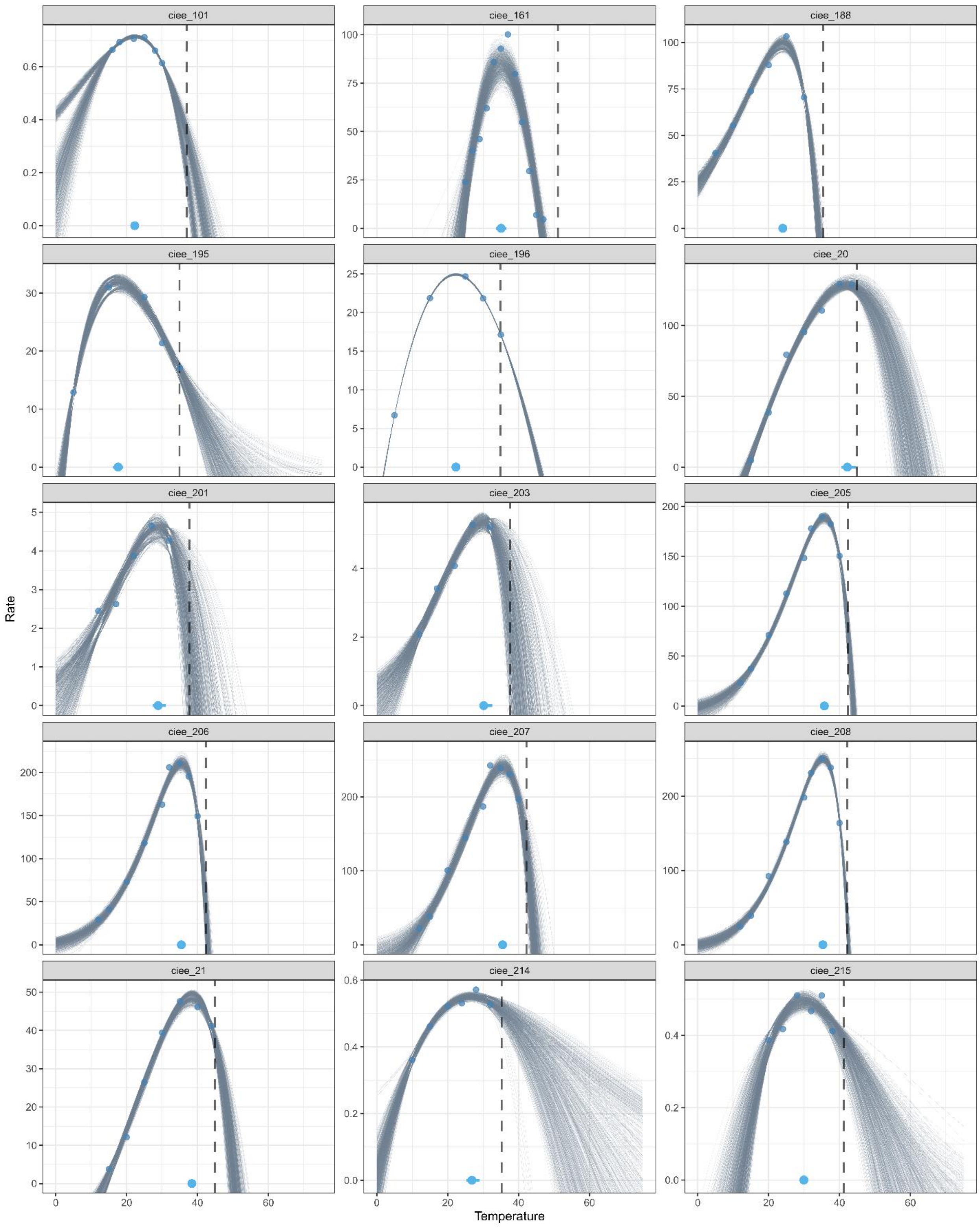
ciee_101
ciee_161
ciee_188
ciee_195
ciee_196
ciee_20
ciee_201
ciee_203
ciee_205
ciee_206
ciee_207
ciee_208
ciee_21
ciee_214
ciee_215
Rate
Temperature

1108

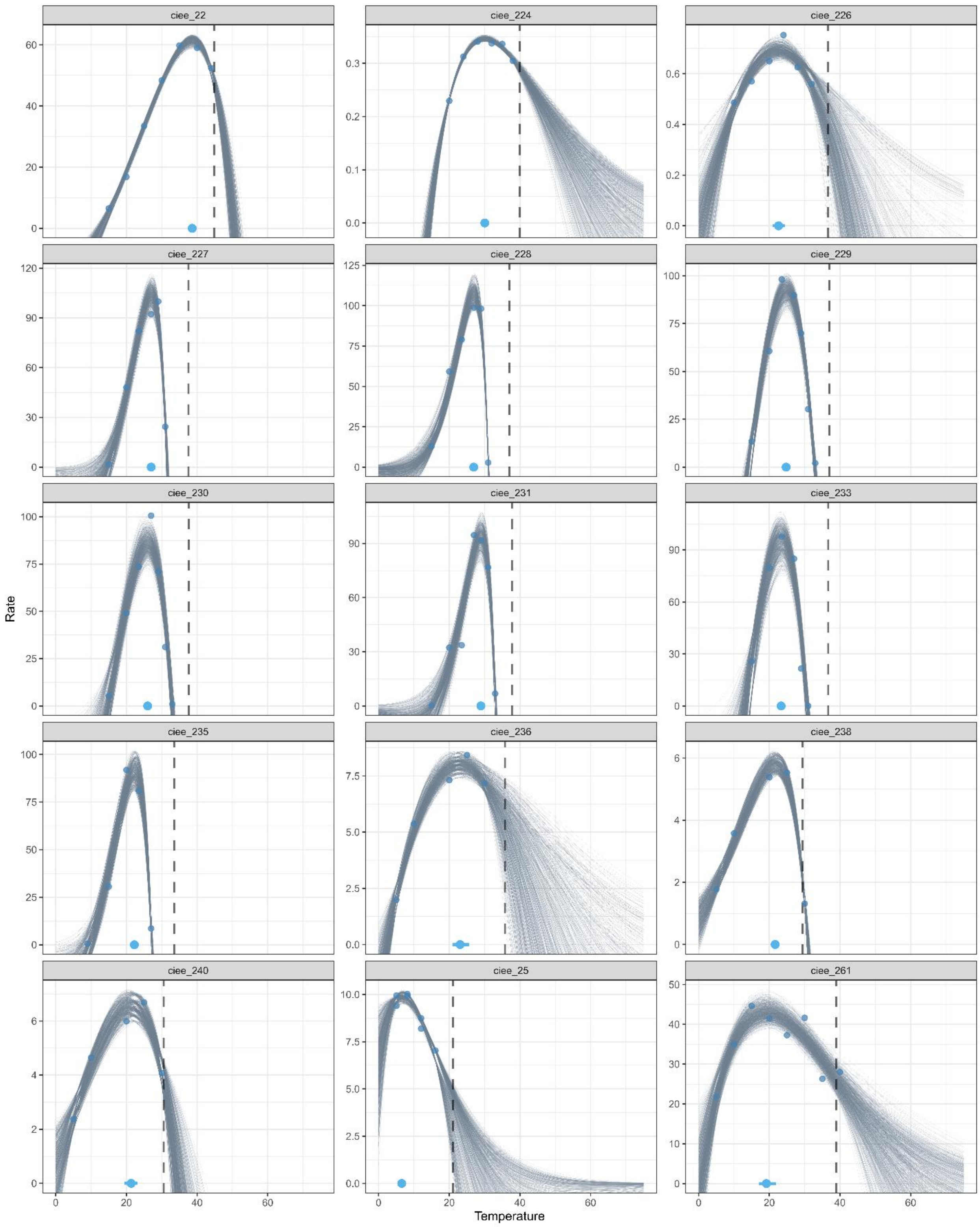

ciee_22
ciee_224
ciee_226
ciee_227
ciee_228
ciee_229
ciee_230
ciee_231
ciee_233
ciee_235
ciee_236
ciee_238
ciee_240
ciee_25
ciee_261
Rate
Temperature

1109

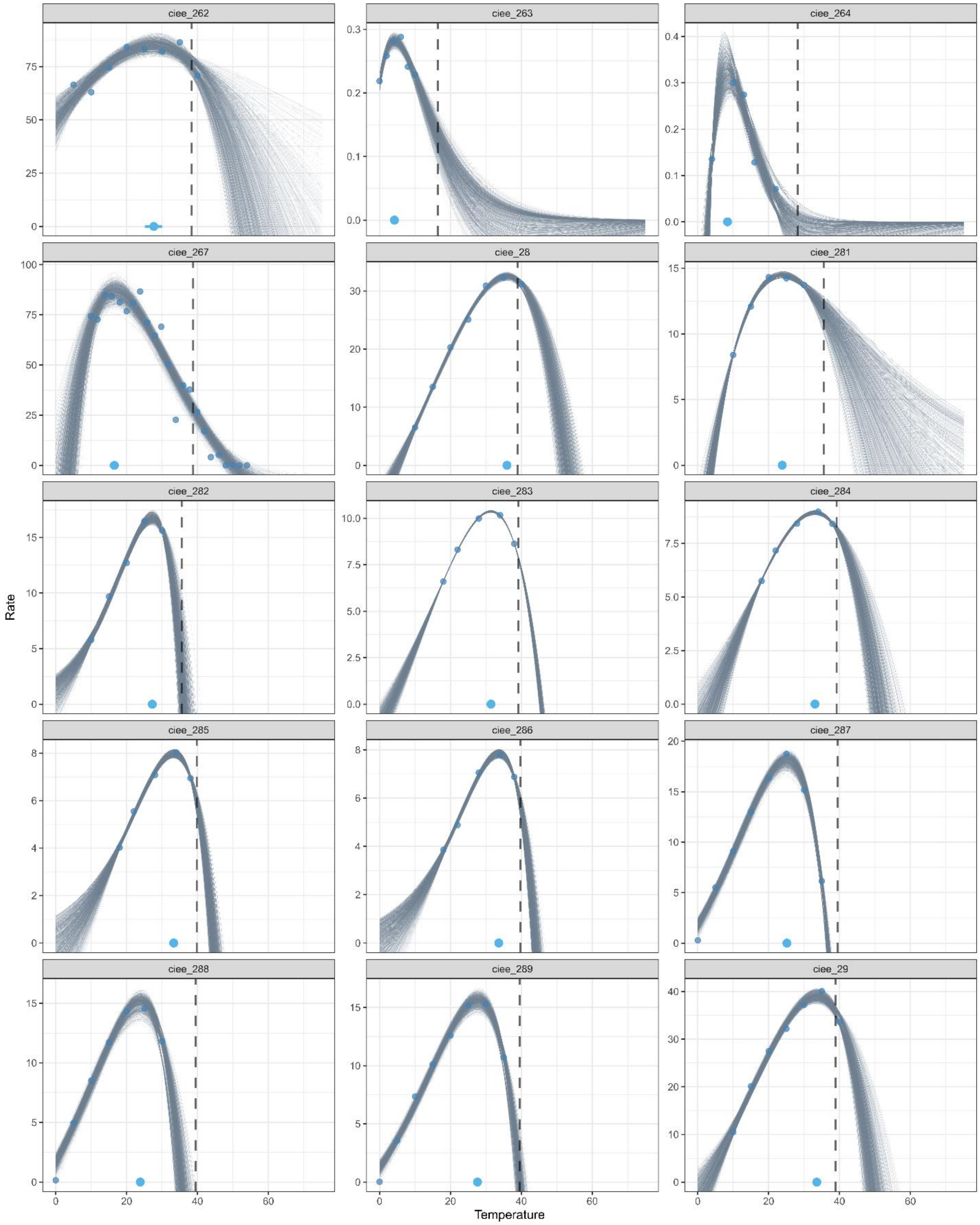

ciee_262
ciee_263
ciee_264
ciee_267
ciee_28
ciee_281
ciee_282
ciee_283
ciee_284
ciee_285
ciee_286
ciee_287
ciee_288
ciee_289
ciee_29
Rate
Temperature

1110

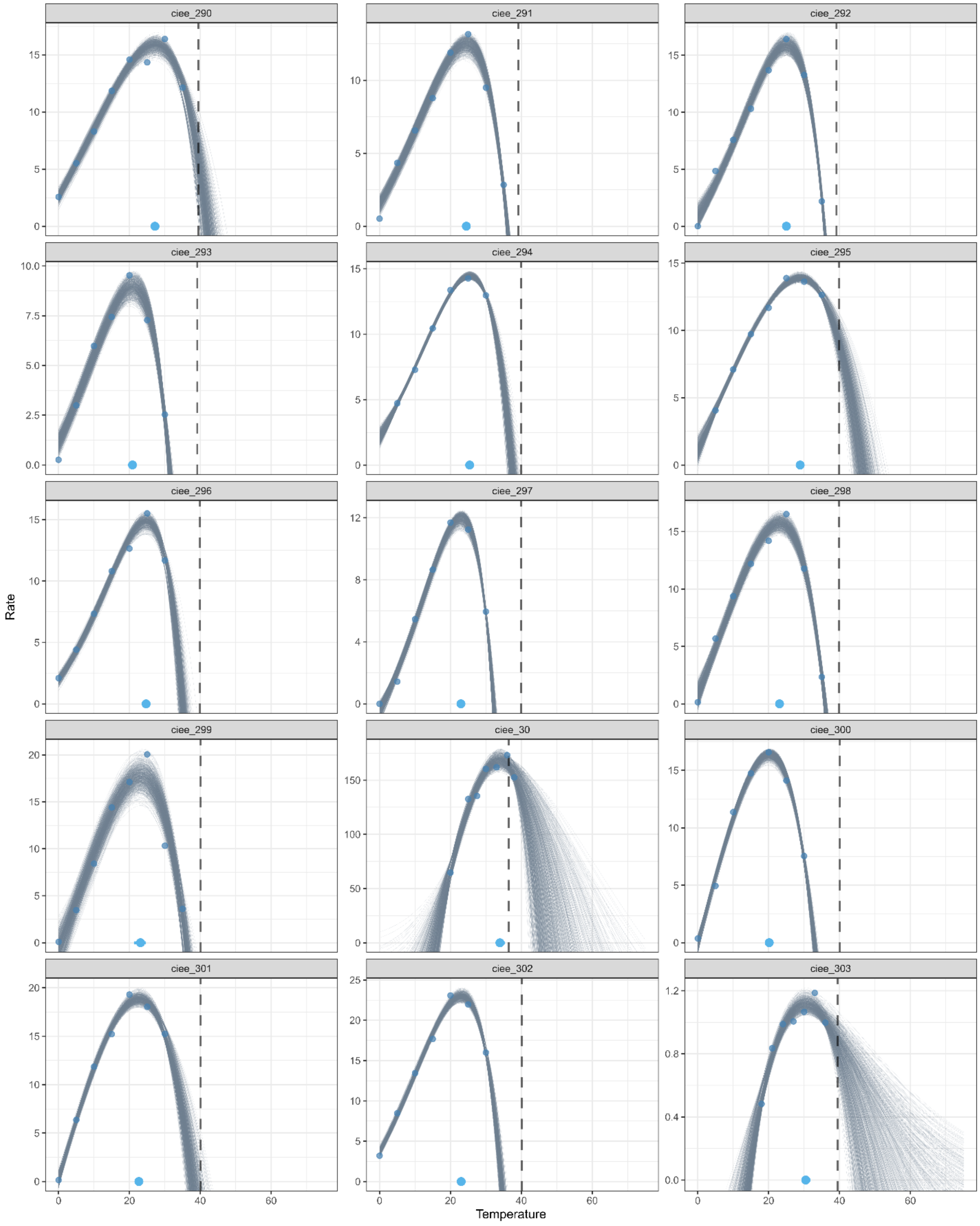

ciee_290
ciee_291
ciee_292
ciee_293
ciee_294
ciee_295
ciee_296
ciee_297
ciee_298
ciee_299
ciee_30
ciee_300
ciee_301
ciee_302
ciee_303
Rate
Temperature

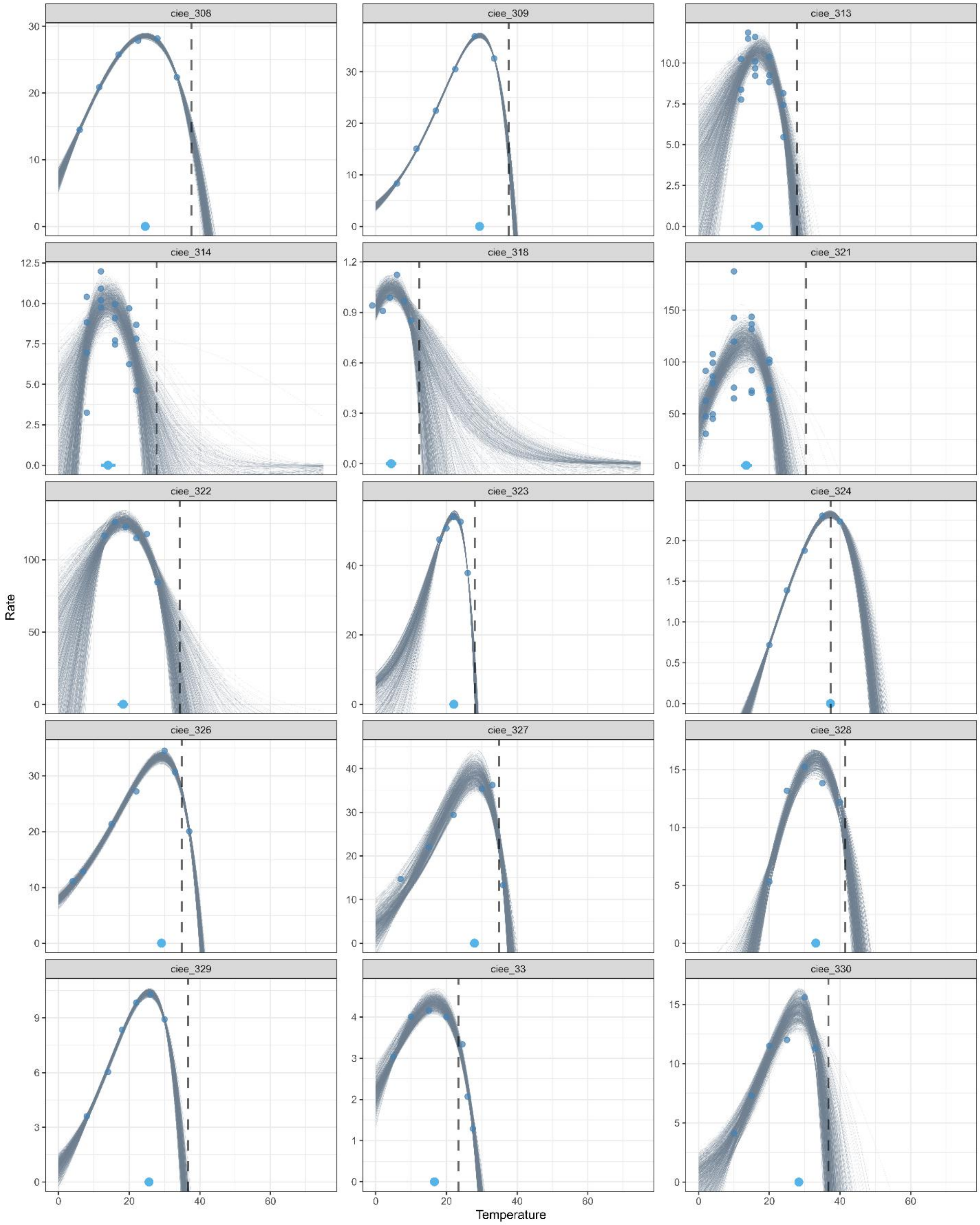
ciee_308
ciee_309
ciee_313
ciee_314
ciee_318
ciee_321
ciee_322
ciee_323
ciee_324
ciee_326
ciee_327
ciee_328
ciee_329
ciee_33
ciee_330
Rate
Temperature

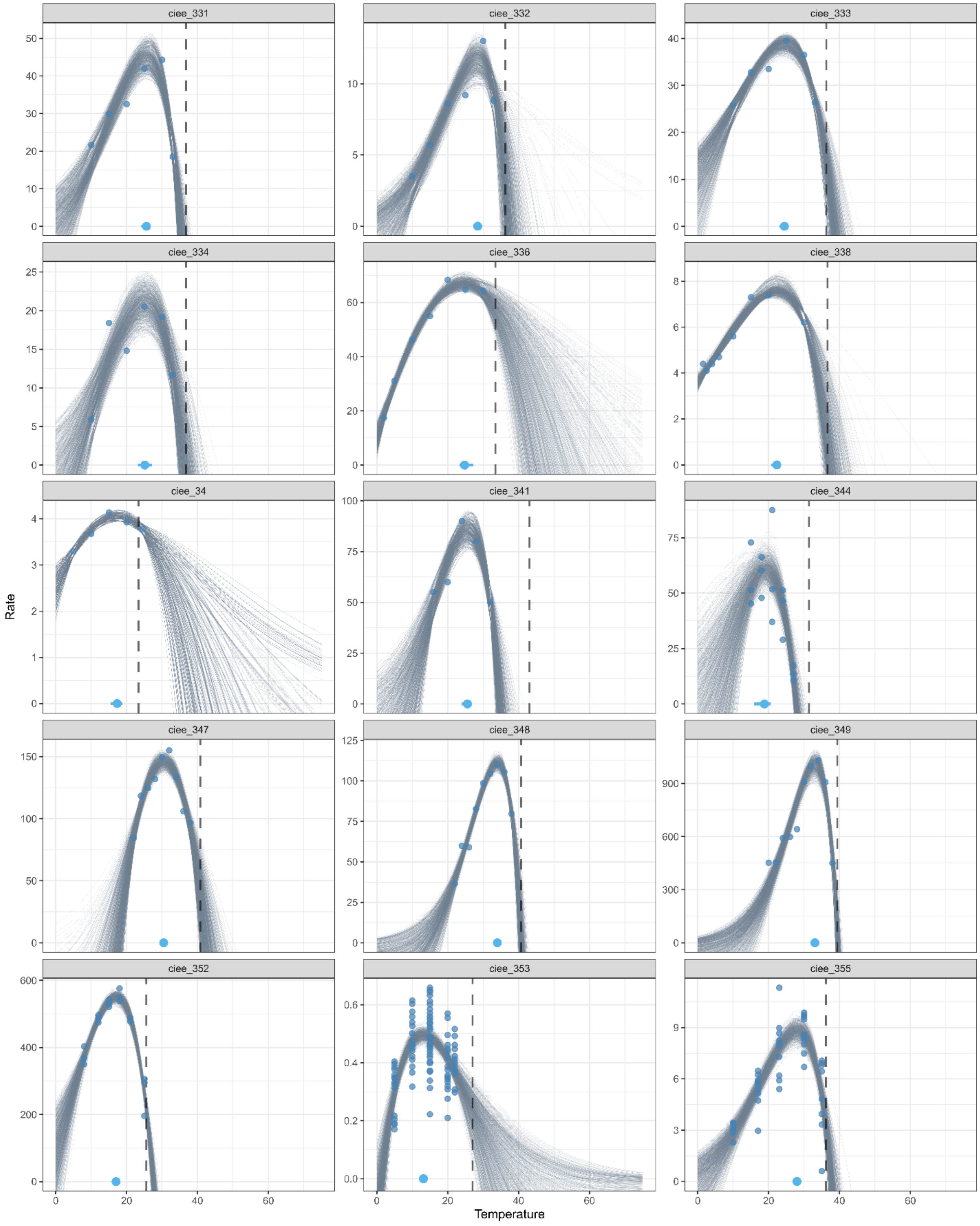
ciee_331
ciee_332
ciee_333
ciee_334
ciee_336
ciee_338
ciee_34
ciee_341
ciee_344
ciee_347
ciee_348
ciee_349
ciee_352
ciee_353
ciee_355
Rate
Temperature

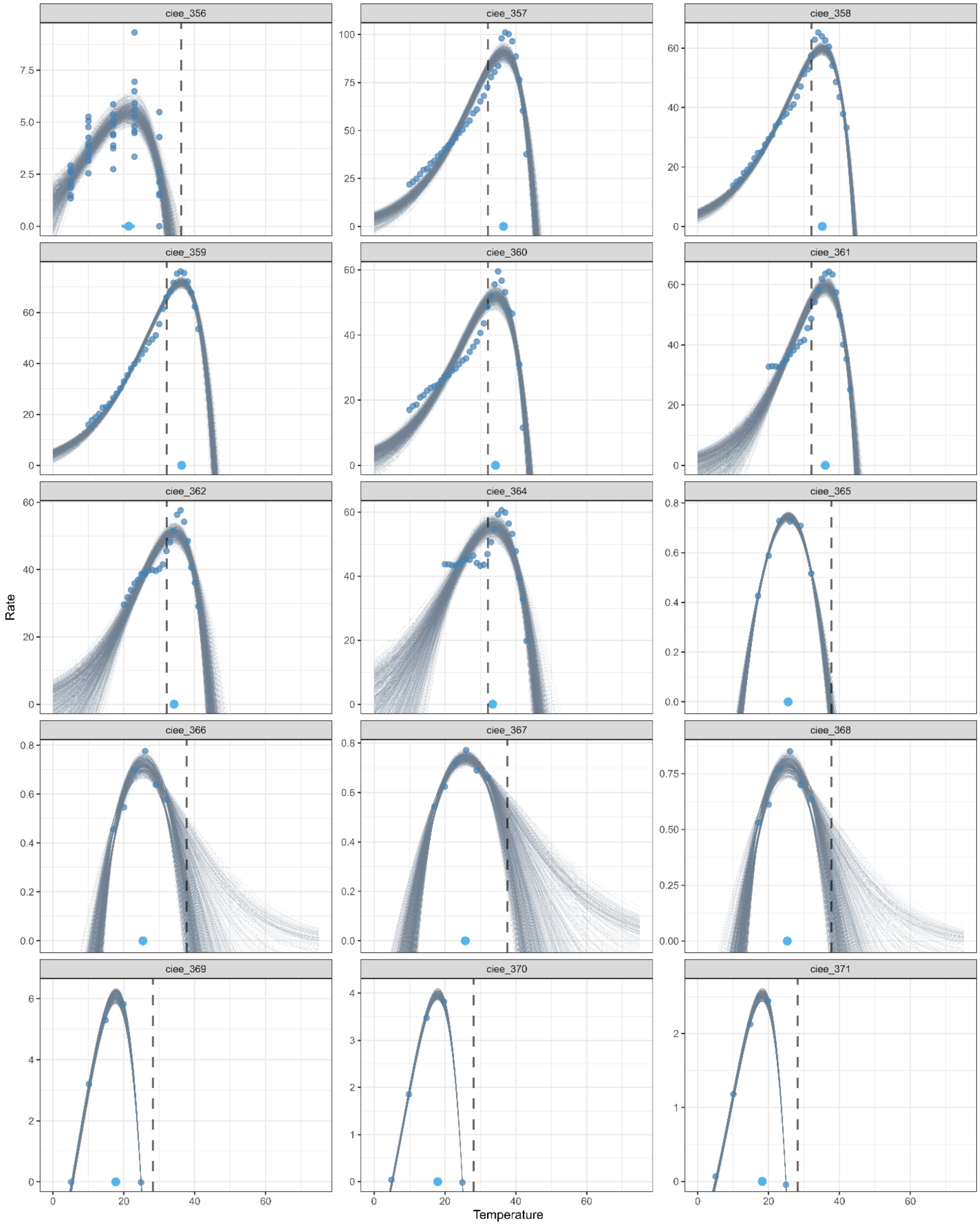
ciee_356
ciee_357
ciee_358
ciee_359
ciee_360
ciee_361
ciee_362
ciee_364
ciee_365
ciee_366
ciee_367
ciee_368
ciee_369
ciee_370
ciee_371
Rate
Temperature

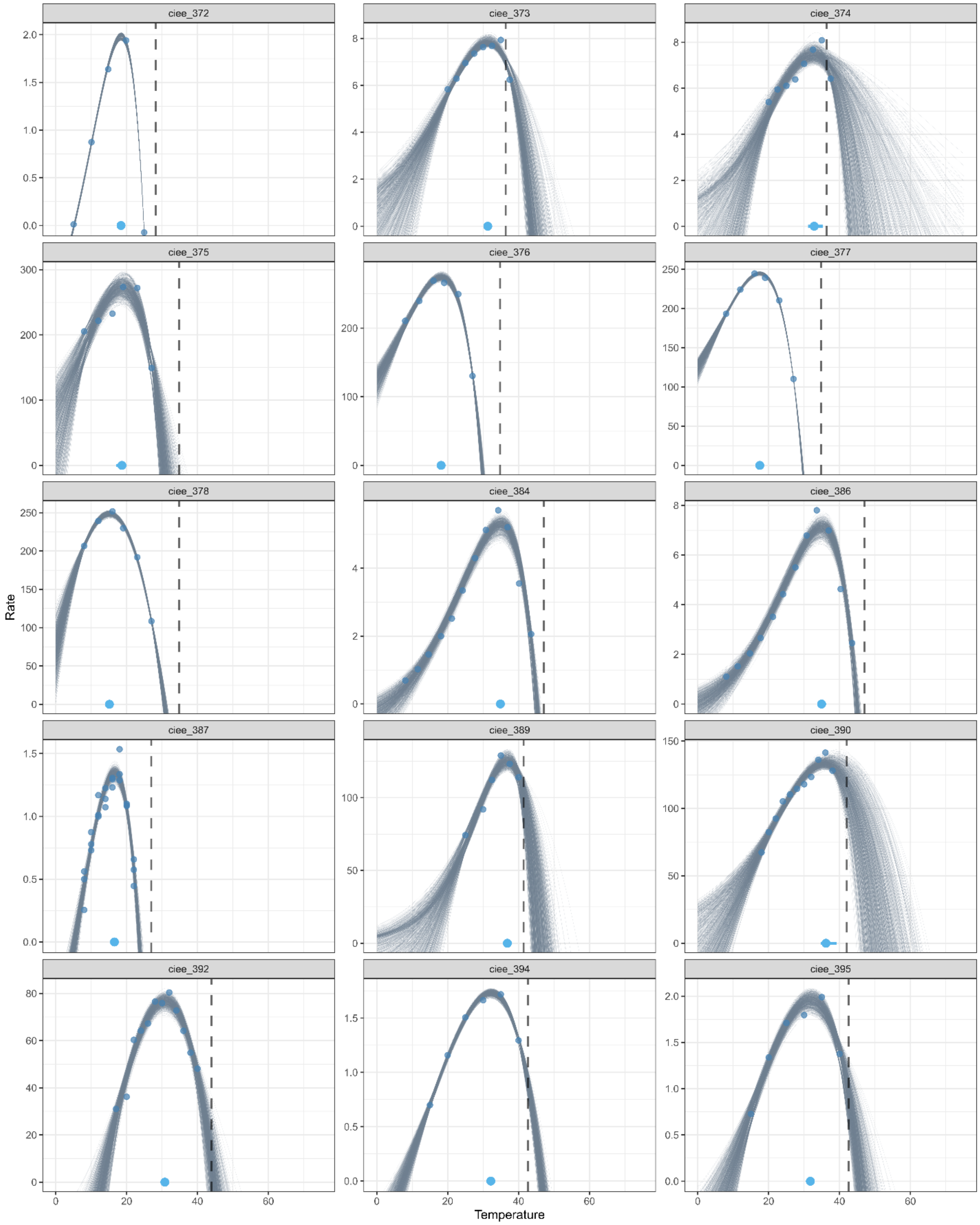
ciee_372
ciee_373
ciee_374
ciee_375
ciee_376
ciee_377
ciee_378
ciee_384
ciee_386
ciee_387
ciee_389
ciee_390
ciee_392
ciee_394
ciee_395
Rate
Temperature

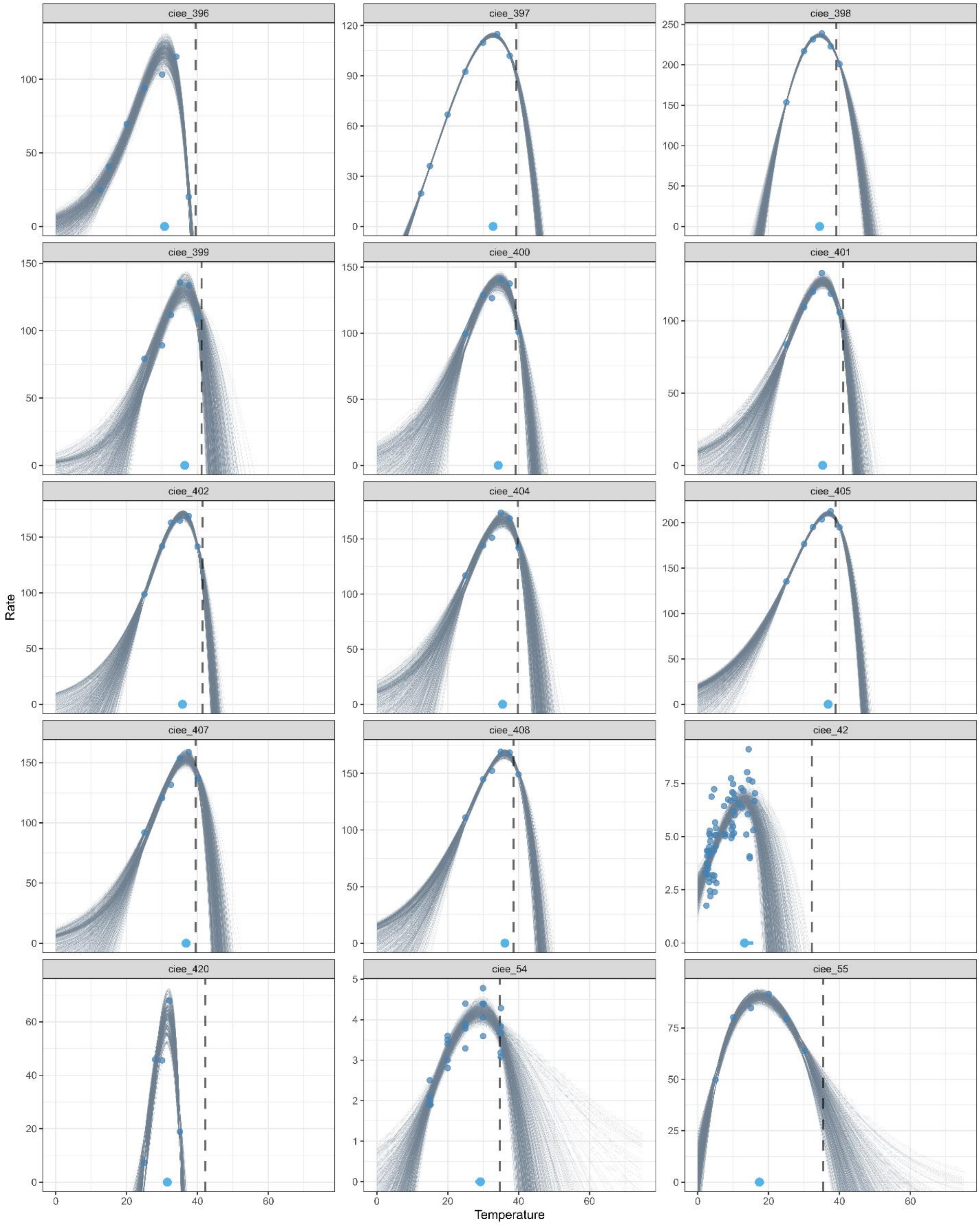
ciee_396
ciee_397
ciee_398
ciee_399
ciee_400
ciee_401
ciee_402
ciee_404
ciee_405
ciee_407
ciee_408
ciee_42
ciee_420
ciee_54
ciee_55
Rate
Temperature

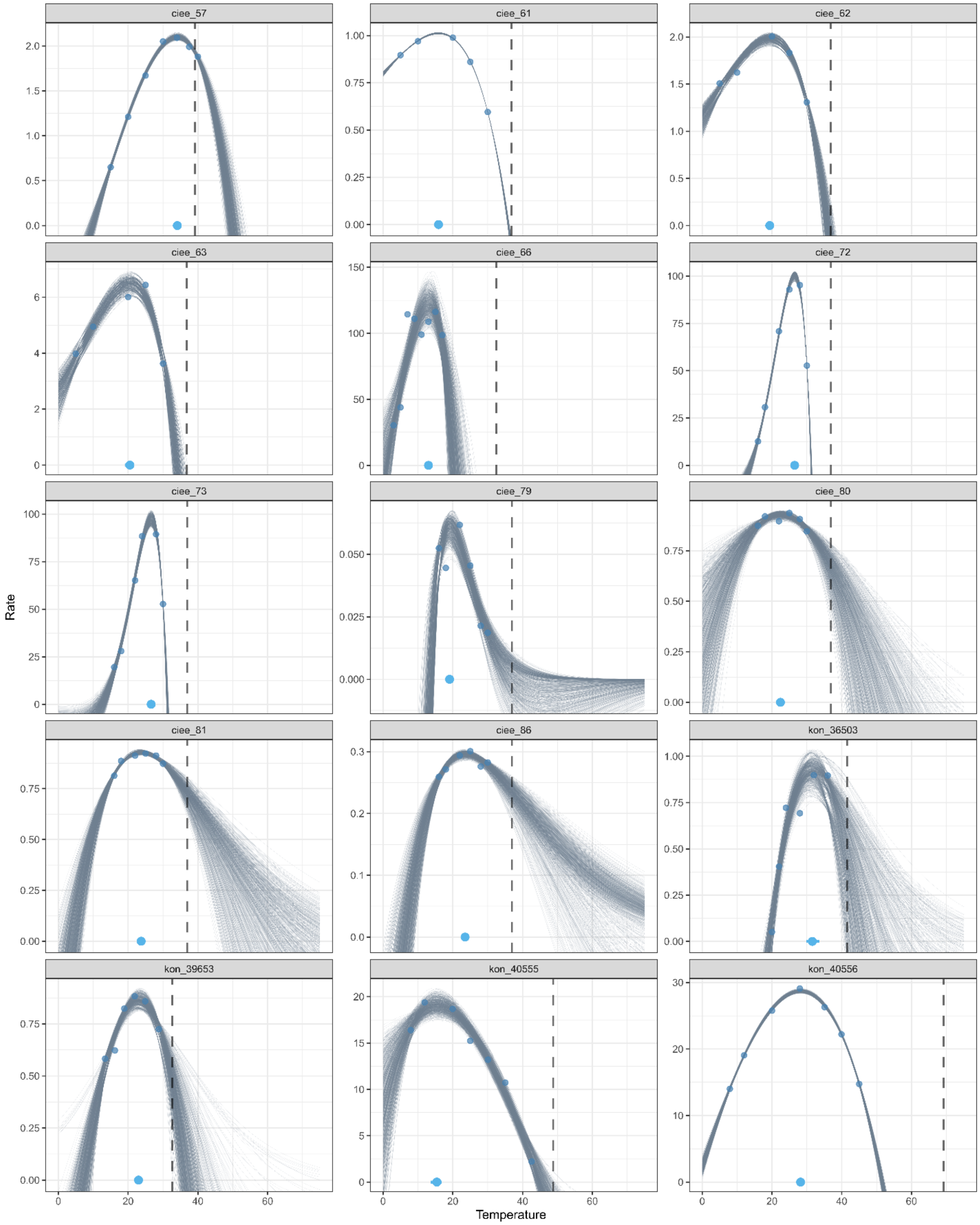

ciee_57
ciee_61
ciee_62
ciee_63
ciee_66
ciee_72
ciee_73
ciee_79
ciee_80
ciee_81
ciee_86
kon_36503
kon_39653
kon_40555
kon_40556
Rate
Temperature

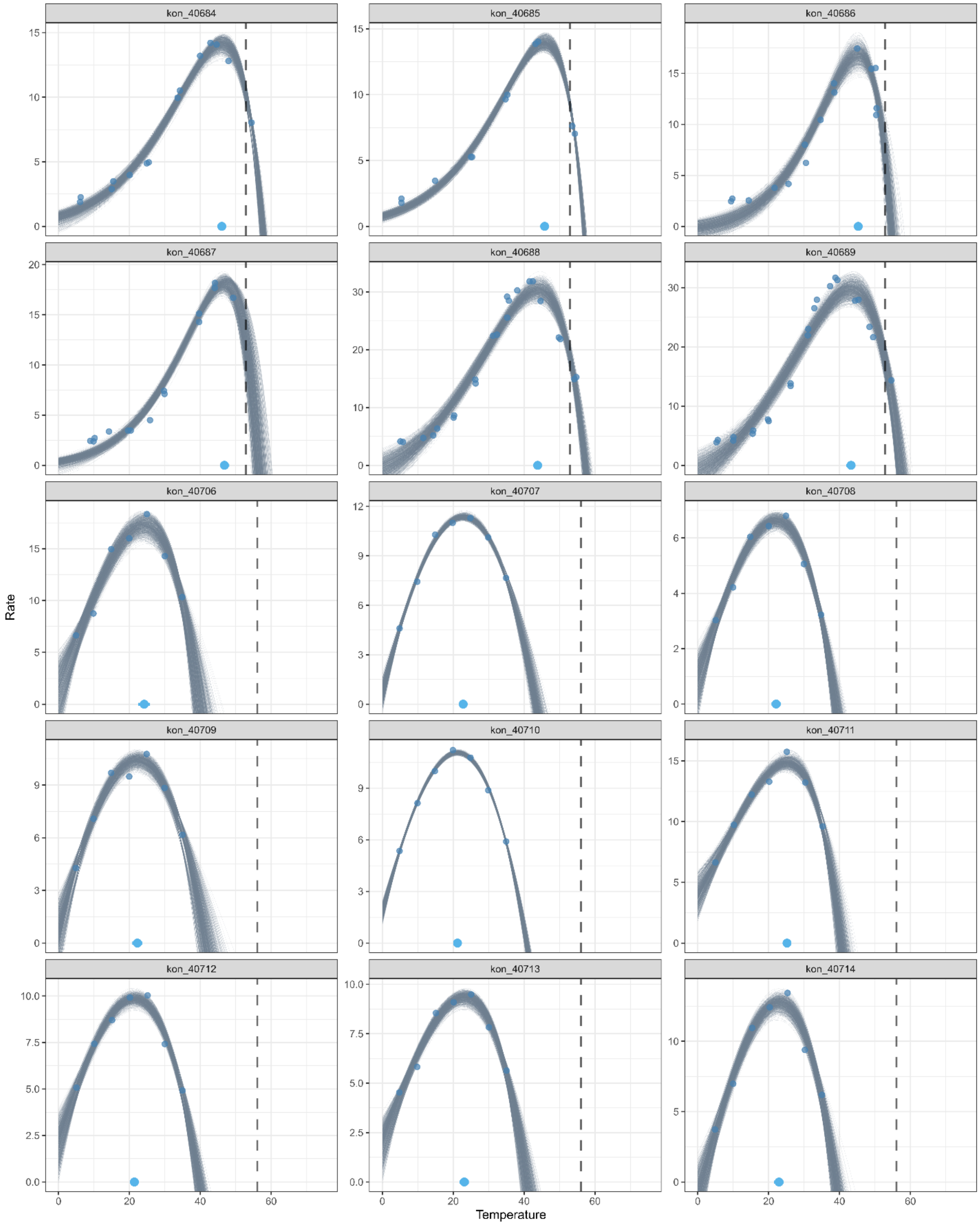
kon_40684
kon_40685
kon_40686
kon_40687
kon_40688
kon_40689
kon_40706
kon_40707
kon_40708
kon_40709
kon_40710
kon_40711
kon_40712
kon_40713
kon_40714
Rate
Temperature

1117

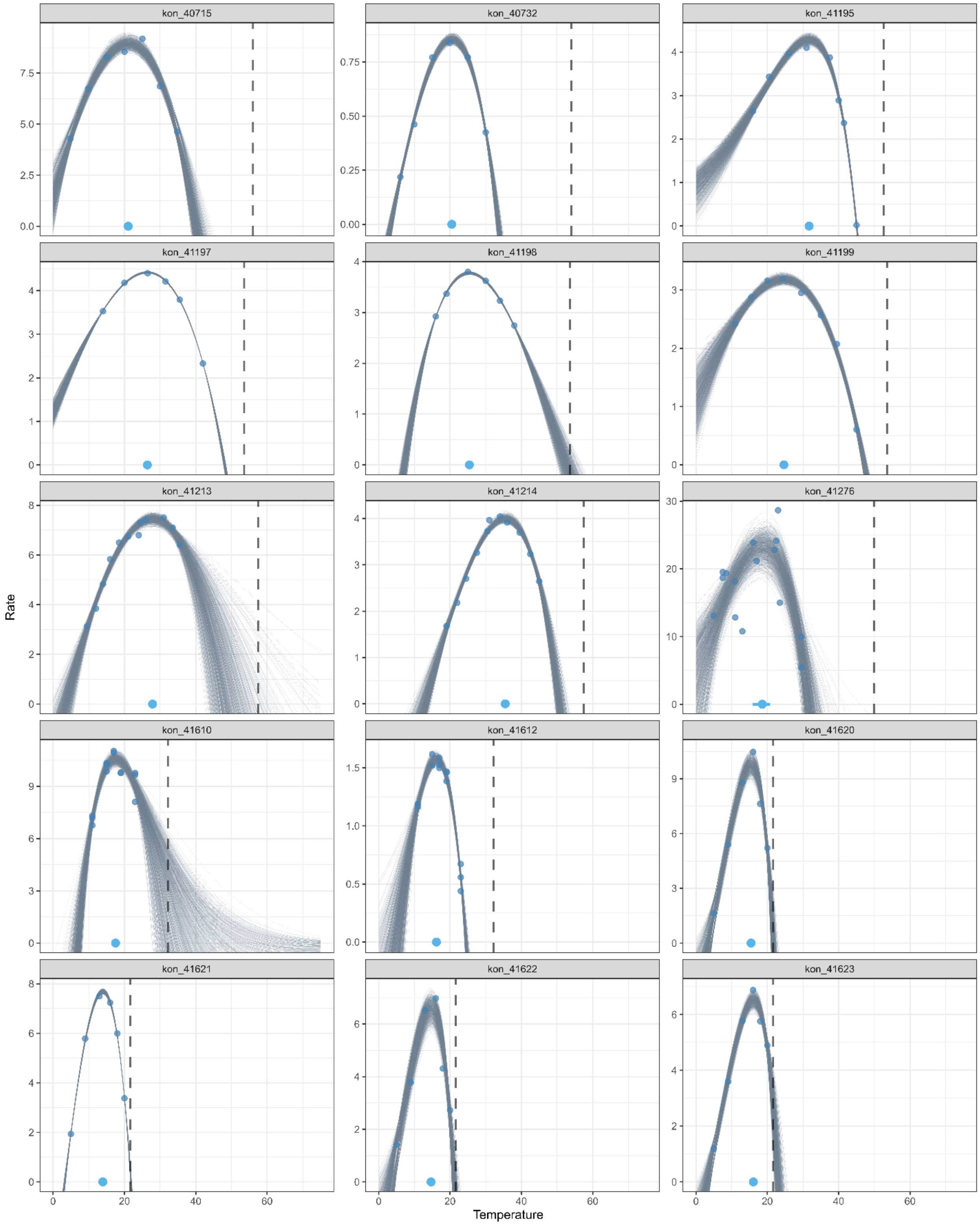
kon_40715
kon_40732
kon_41195
kon_41197
kon_41198
kon_41199
kon_41213
kon_41214
kon_41276
kon_41610
kon_41612
kon_41620
kon_41621
kon_41622
kon_41623
Rate
Temperature
0
20
40
60

1118

1119

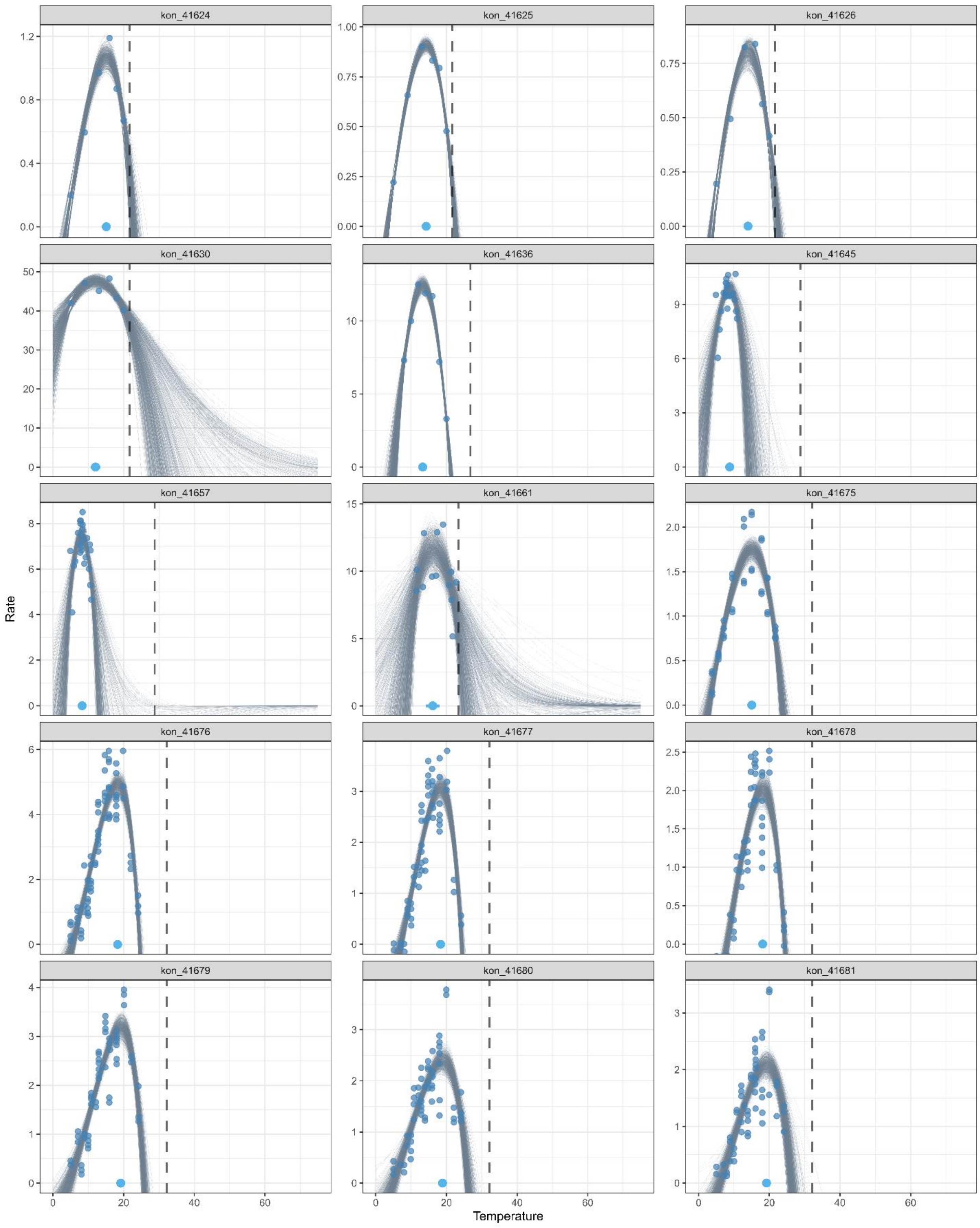

kon_41624
kon_41625
kon_41626
kon_41630
kon_41636
kon_41645
kon_41657
kon_41661
kon_41675
kon_41676
kon_41677
kon_41678
kon_41679
kon_41680
kon_41681
Rate
Temperature

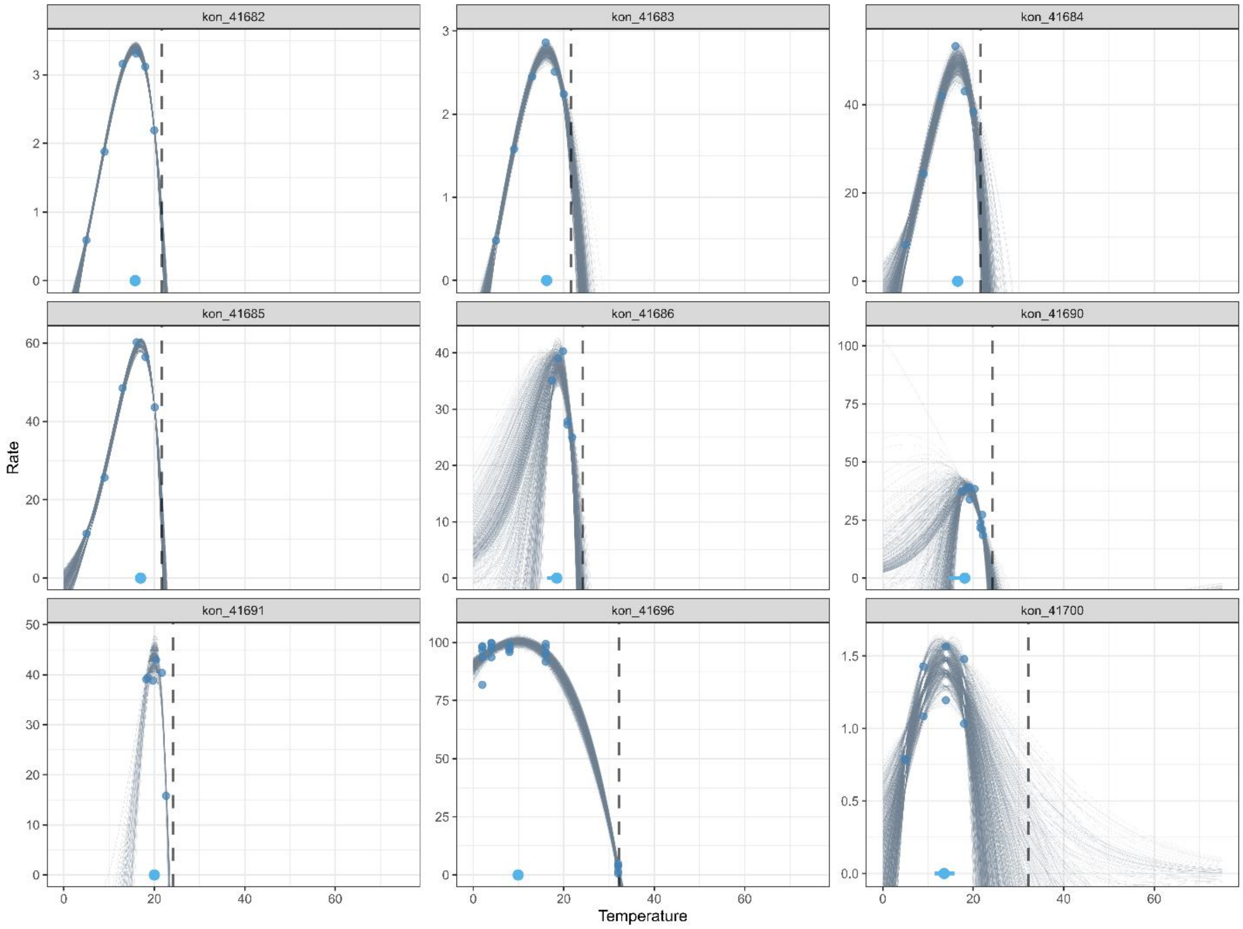
kon_41682
kon_41683
kon_41684
kon_41685
kon_41686
kon_41690
kon_41691
kon_41696
kon_41700
Rate
Temperature

**ii. $TPC_{max}$ dataset**

Below are the bootstrapped thermal performance curves (TPCs) for each empirical dataset (facets), fitted using the Norberg-Thomas model and used to estimate the upper limit of performance ($TPC_{max}$; N = 159). Thin blue-grey lines represent individual bootstrap fits of parameters (N = 1000) used to summarize $TPC_{max}$ and its variance for use in multi-level regression models. Empirical observations of performane are shown as small blue points. The estimates of $TPC_{max}$ are summarized by green points (mean) and horizontal bars indicating 95% confidence intervals, located on the x-axis. Dark grey dashed lines denote the mean species $CT_{max}$, ARR-corrected to a standardized temperature of 15°C. Curves are plotted across a standardized temperature range (0-75°C) and panels are scaled independently along the y-axis.

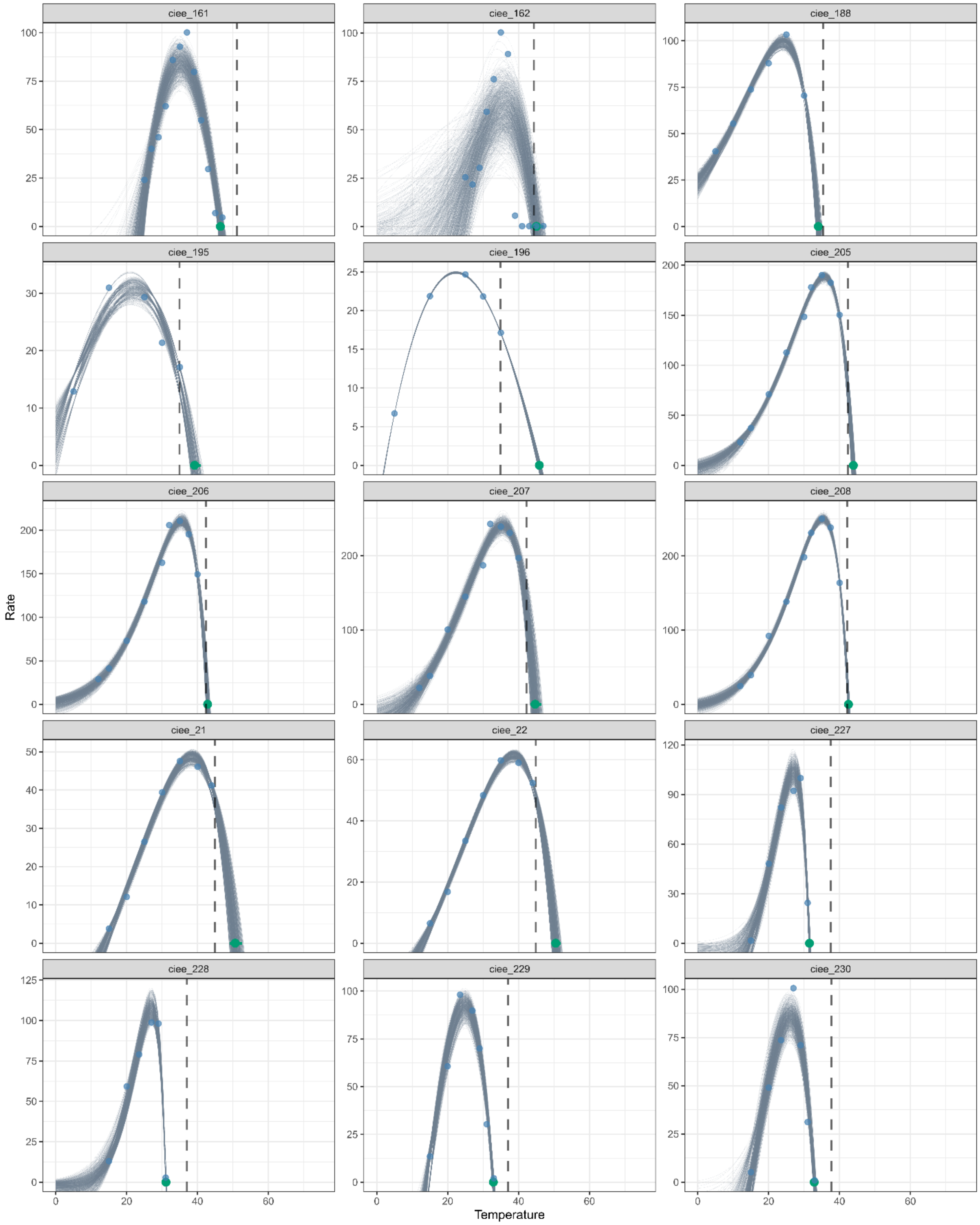
ciee_161
ciee_162
ciee_188
ciee_195
ciee_196
ciee_205
ciee_206
ciee_207
ciee_208
ciee_21
ciee_22
ciee_227
ciee_228
ciee_229
ciee_230
Rate
Temperature

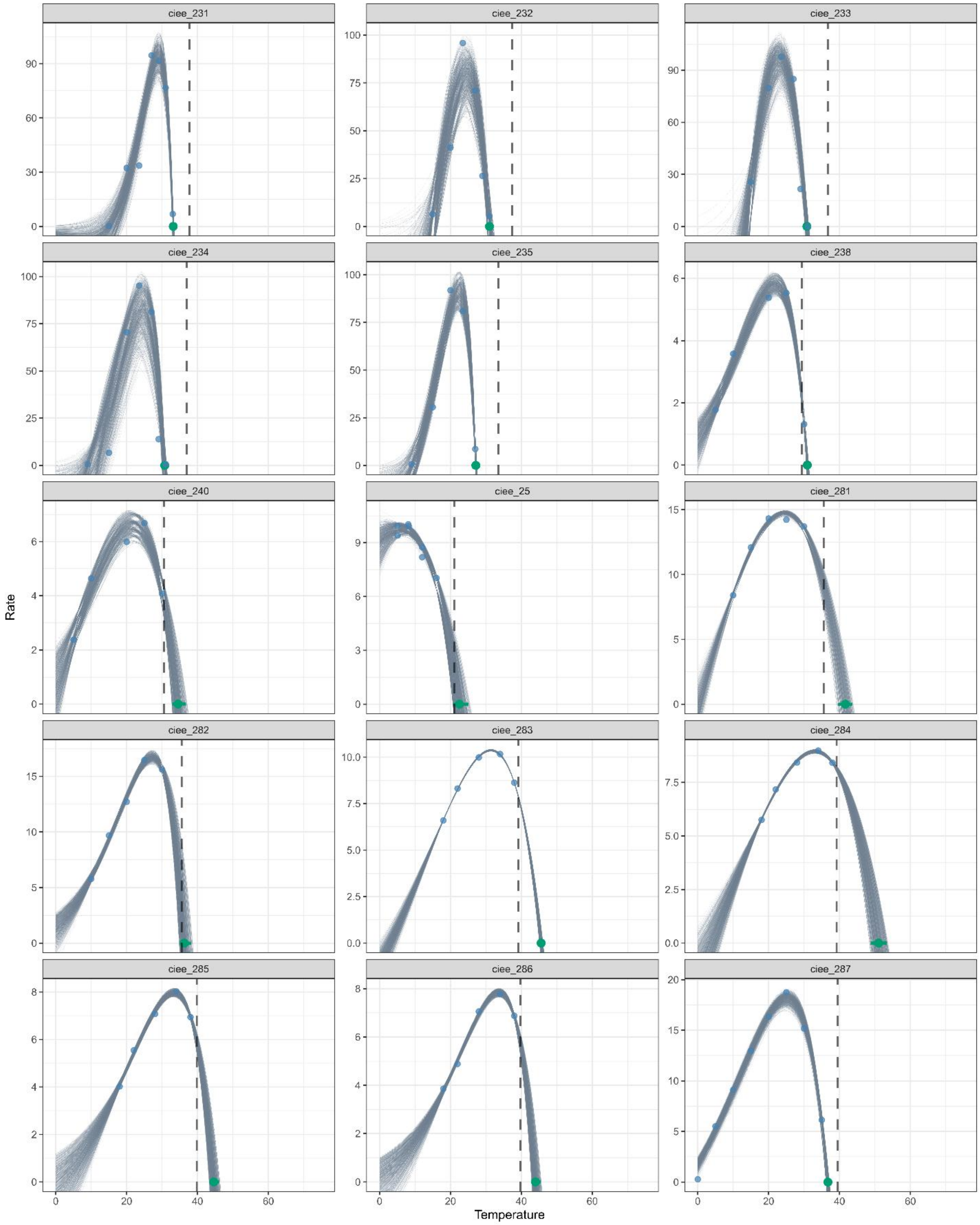
ciee_231
ciee_232
ciee_233
ciee_234
ciee_235
ciee_238
ciee_240
ciee_25
ciee_281
ciee_282
ciee_283
ciee_284
ciee_285
ciee_286
ciee_287
Rate
Temperature

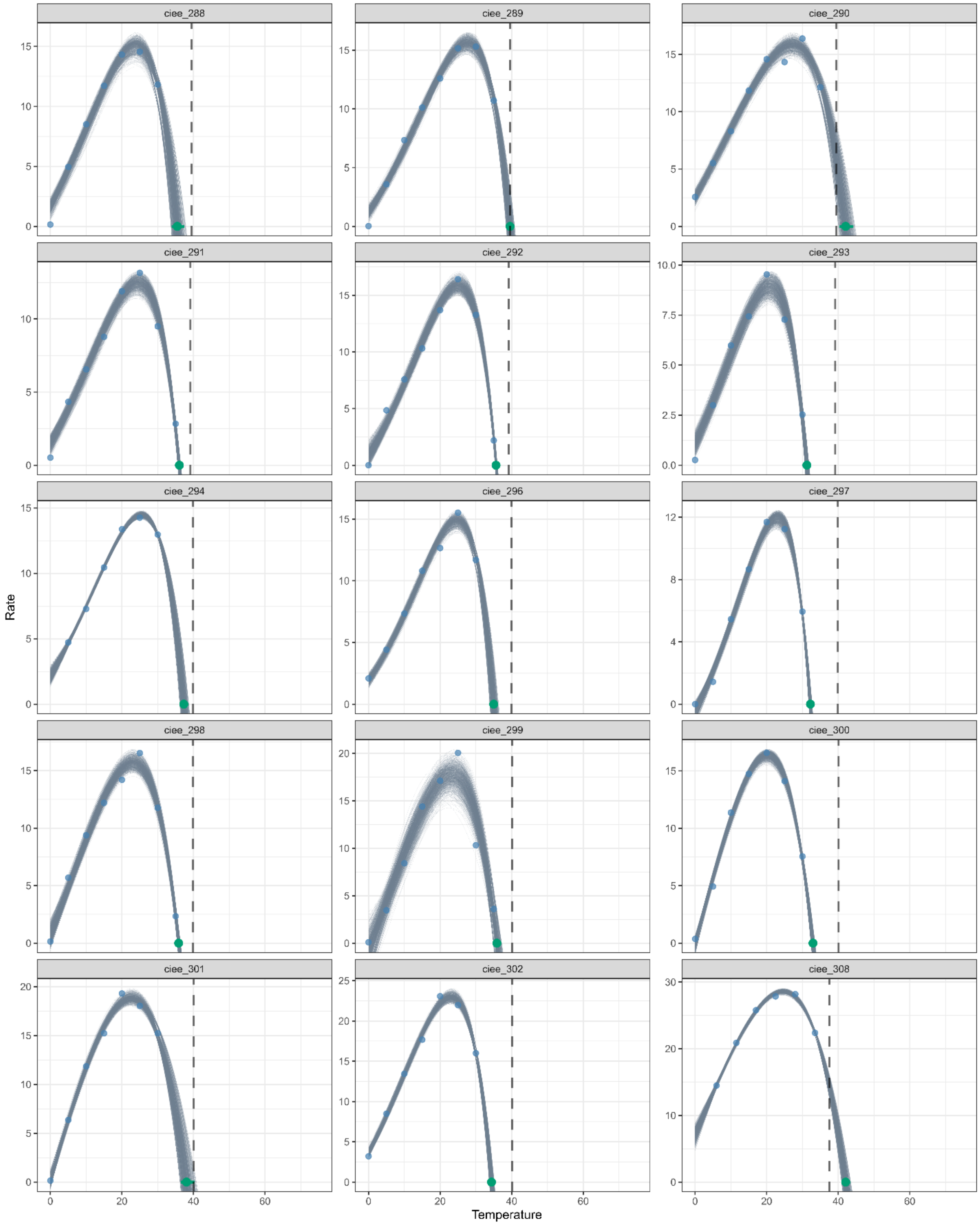
ciee_288
ciee_289
ciee_290
ciee_291
ciee_292
ciee_293
ciee_294
ciee_296
ciee_297
ciee_298
ciee_299
ciee_300
ciee_301
ciee_302
ciee_308
Rate
Temperature

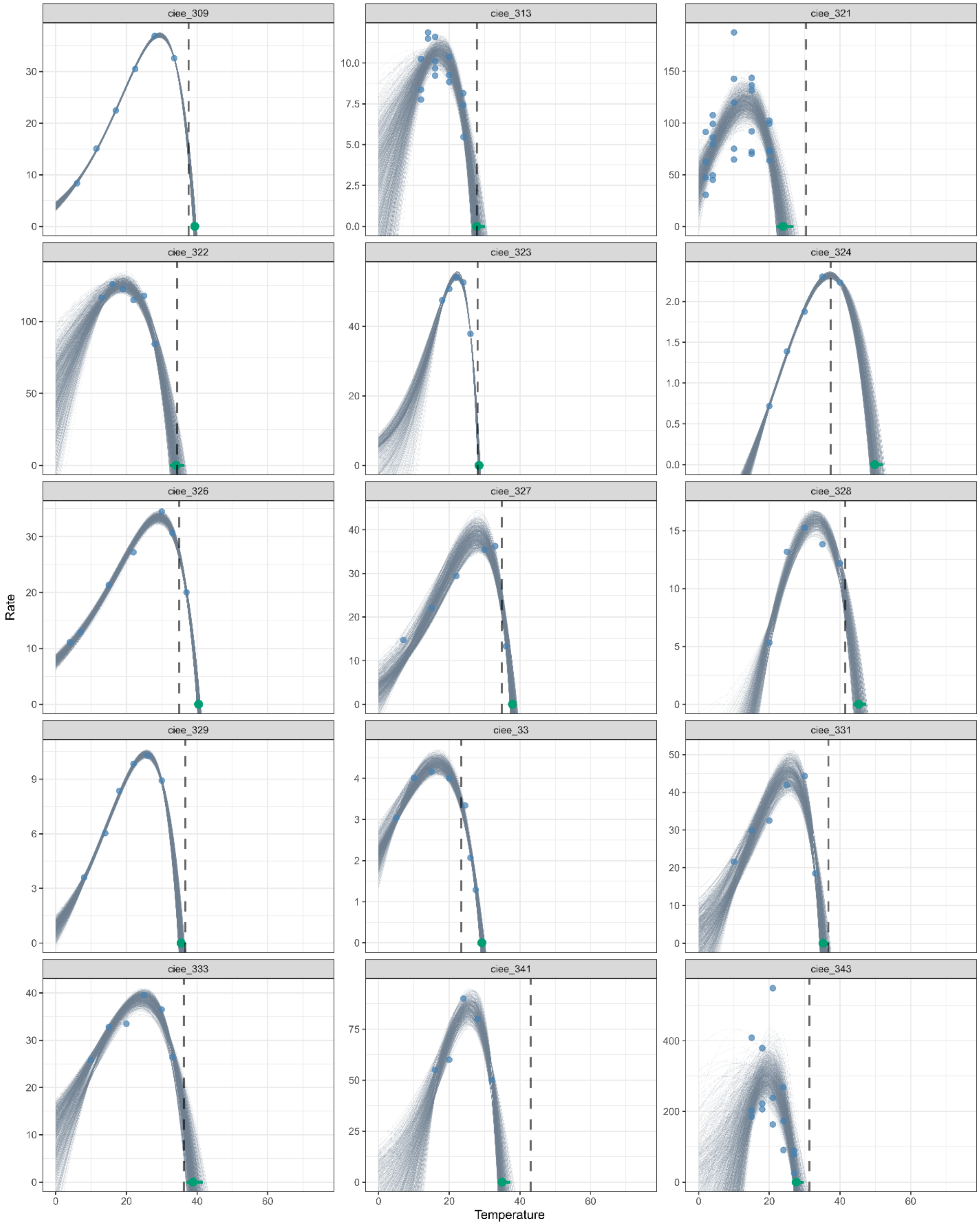
ciee_309
ciee_313
ciee_321
ciee_322
ciee_323
ciee_324
ciee_326
ciee_327
ciee_328
ciee_329
ciee_33
ciee_331
ciee_333
ciee_341
ciee_343
Rate
Temperature

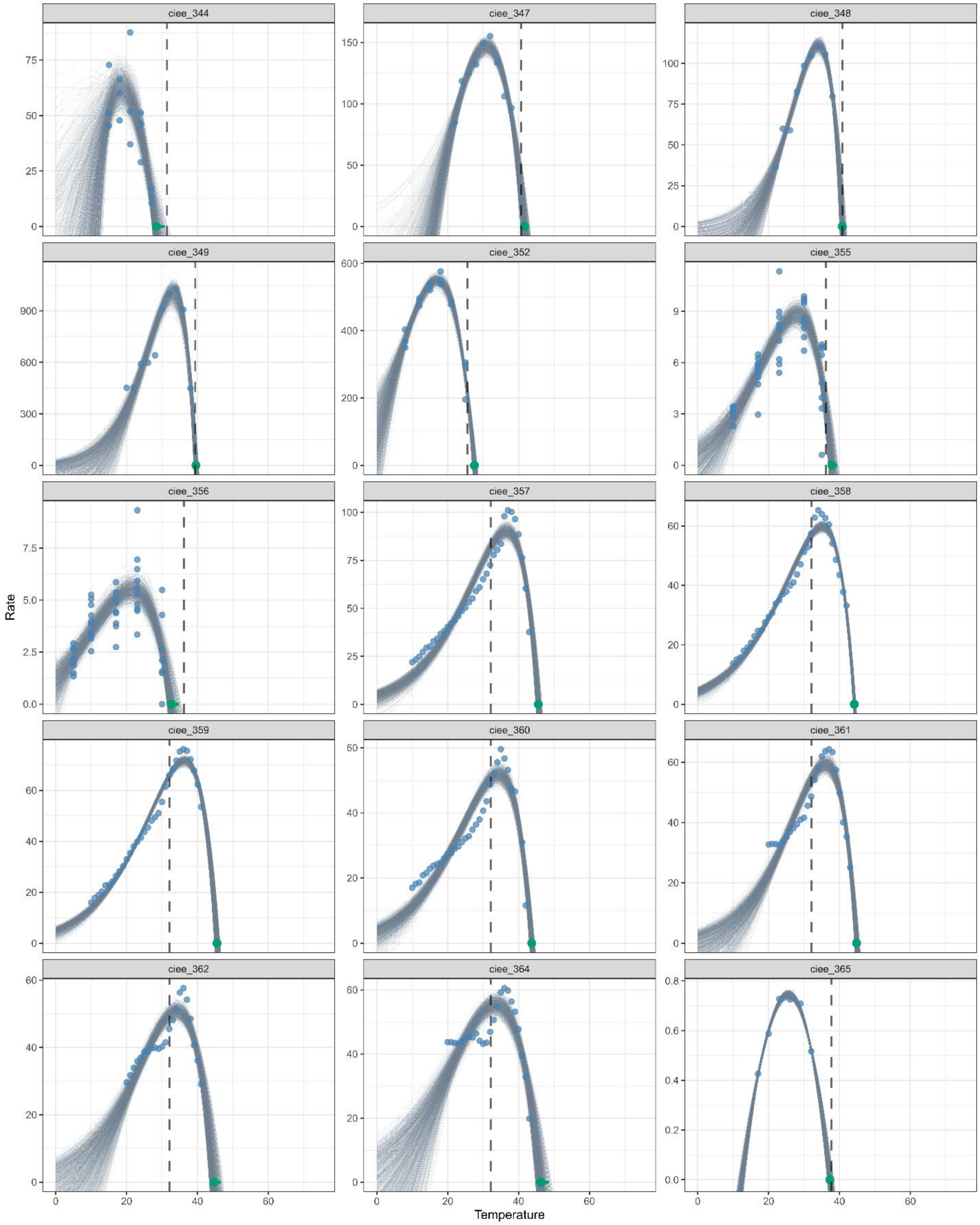
ciee_344
ciee_347
ciee_348
ciee_349
ciee_352
ciee_355
ciee_356
ciee_357
ciee_358
ciee_359
ciee_360
ciee_361
ciee_362
ciee_364
ciee_365
Rate
Temperature

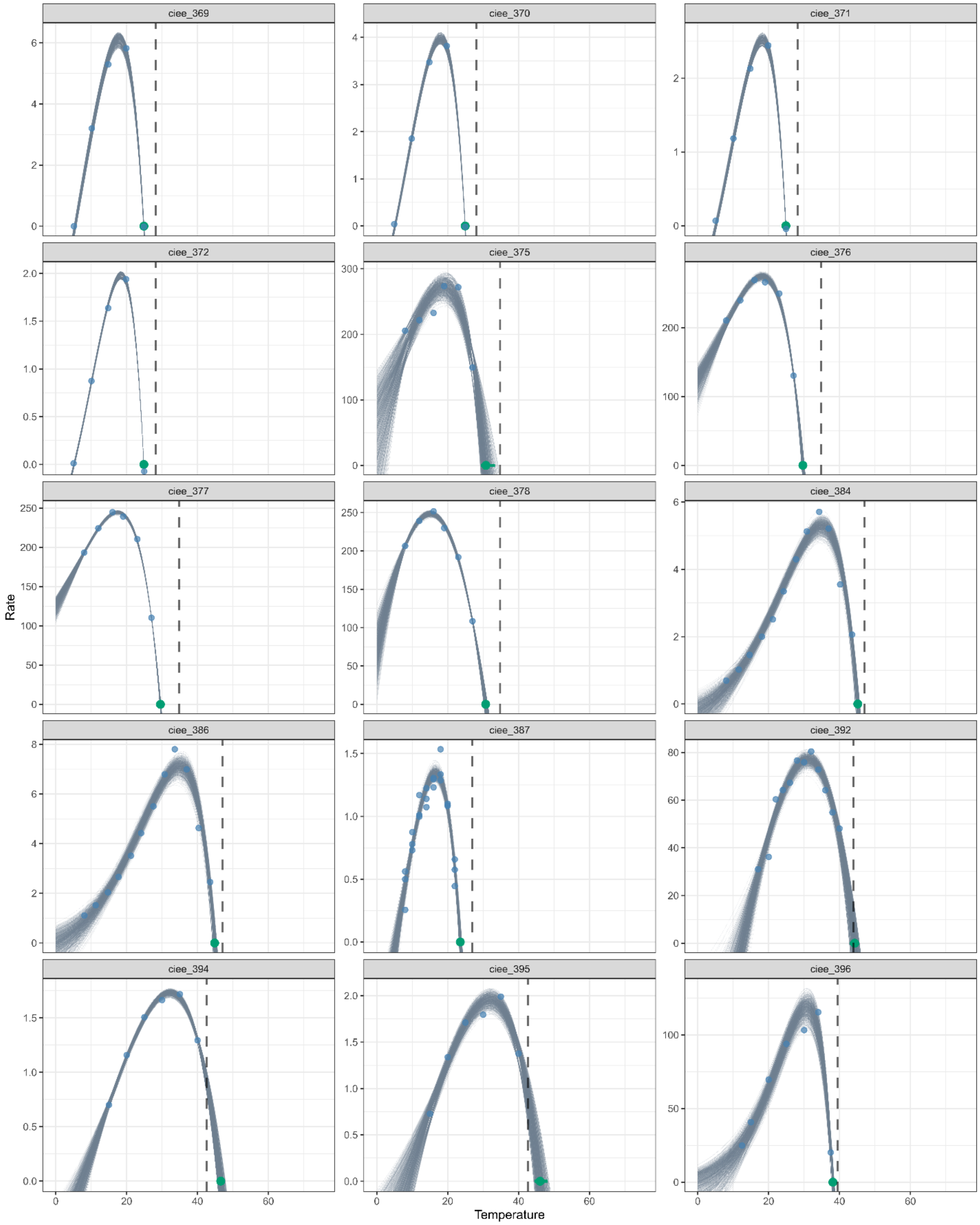

ciee_369
ciee_370
ciee_371
ciee_372
ciee_375
ciee_376
ciee_377
ciee_378
ciee_384
ciee_386
ciee_387
ciee_392
ciee_394
ciee_395
ciee_396
Rate
Temperature

1139

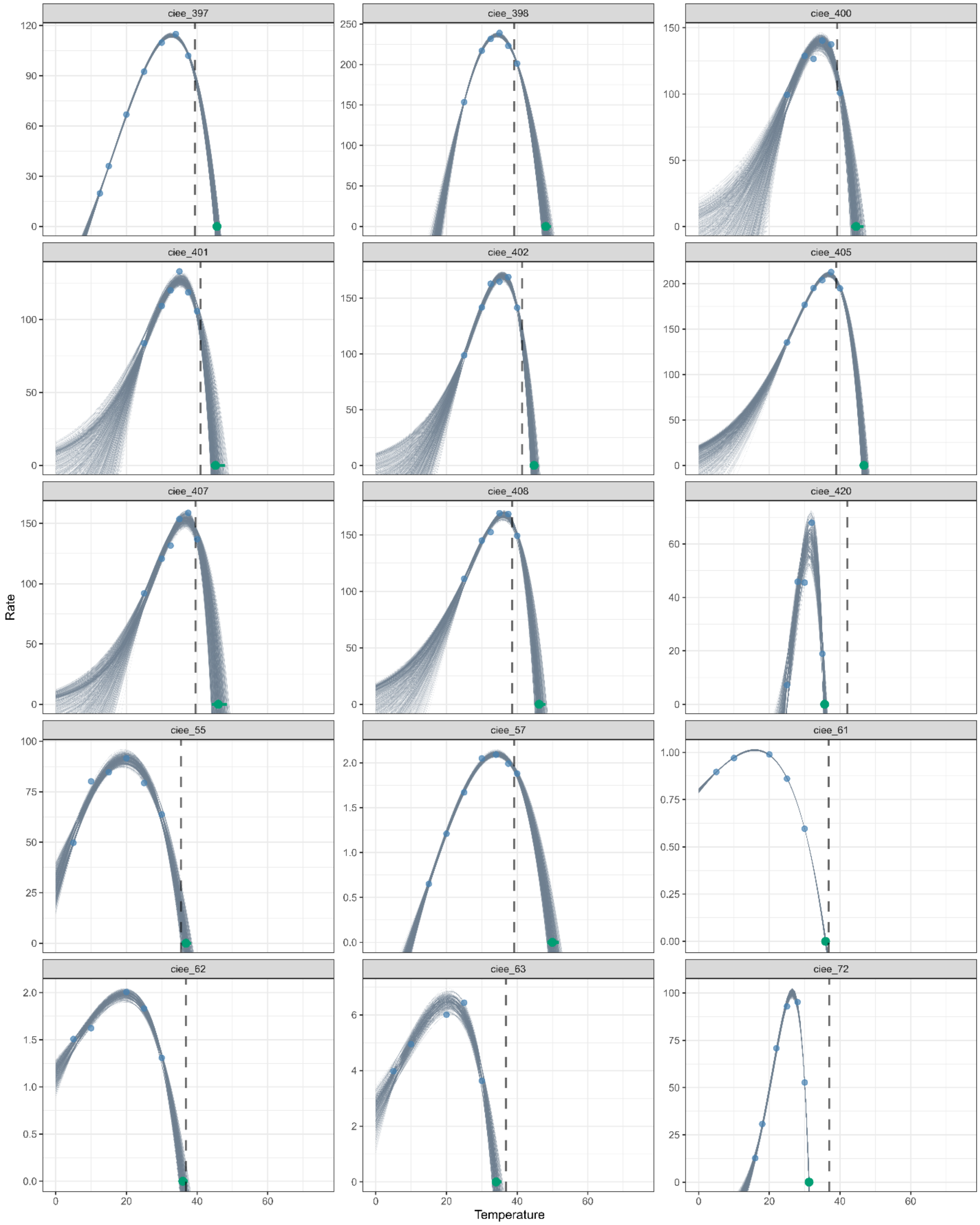
ciee_397
ciee_398
ciee_400
ciee_401
ciee_402
ciee_405
ciee_407
ciee_408
ciee_420
ciee_55
ciee_57
ciee_61
ciee_62
ciee_63
ciee_72
Rate
Temperature

1140

1141

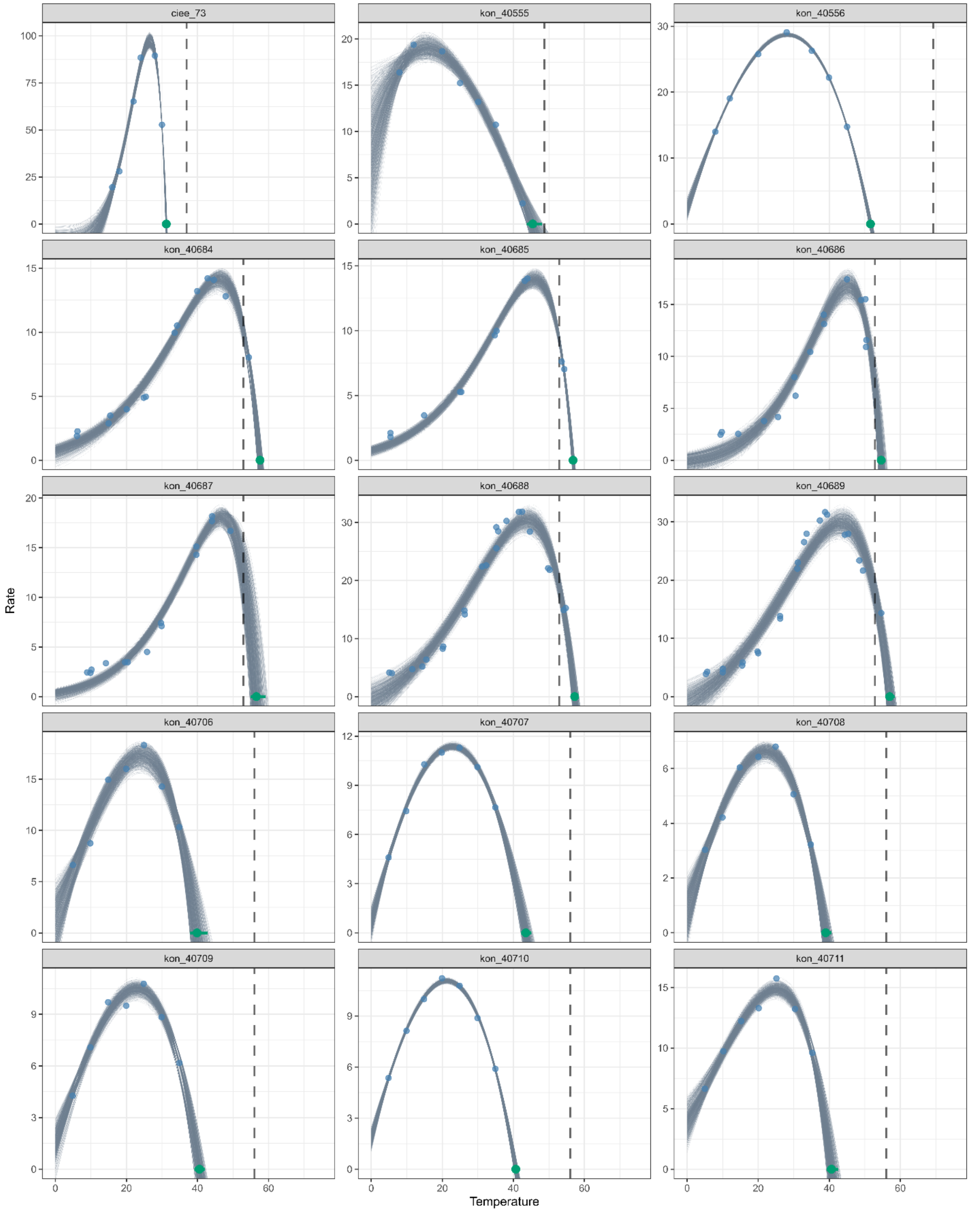

ciee_73
kon_40555
kon_40556
kon_40684
kon_40685
kon_40686
kon_40687
kon_40688
kon_40689
kon_40706
kon_40707
kon_40708
kon_40709
kon_40710
kon_40711
Rate
Temperature

1142
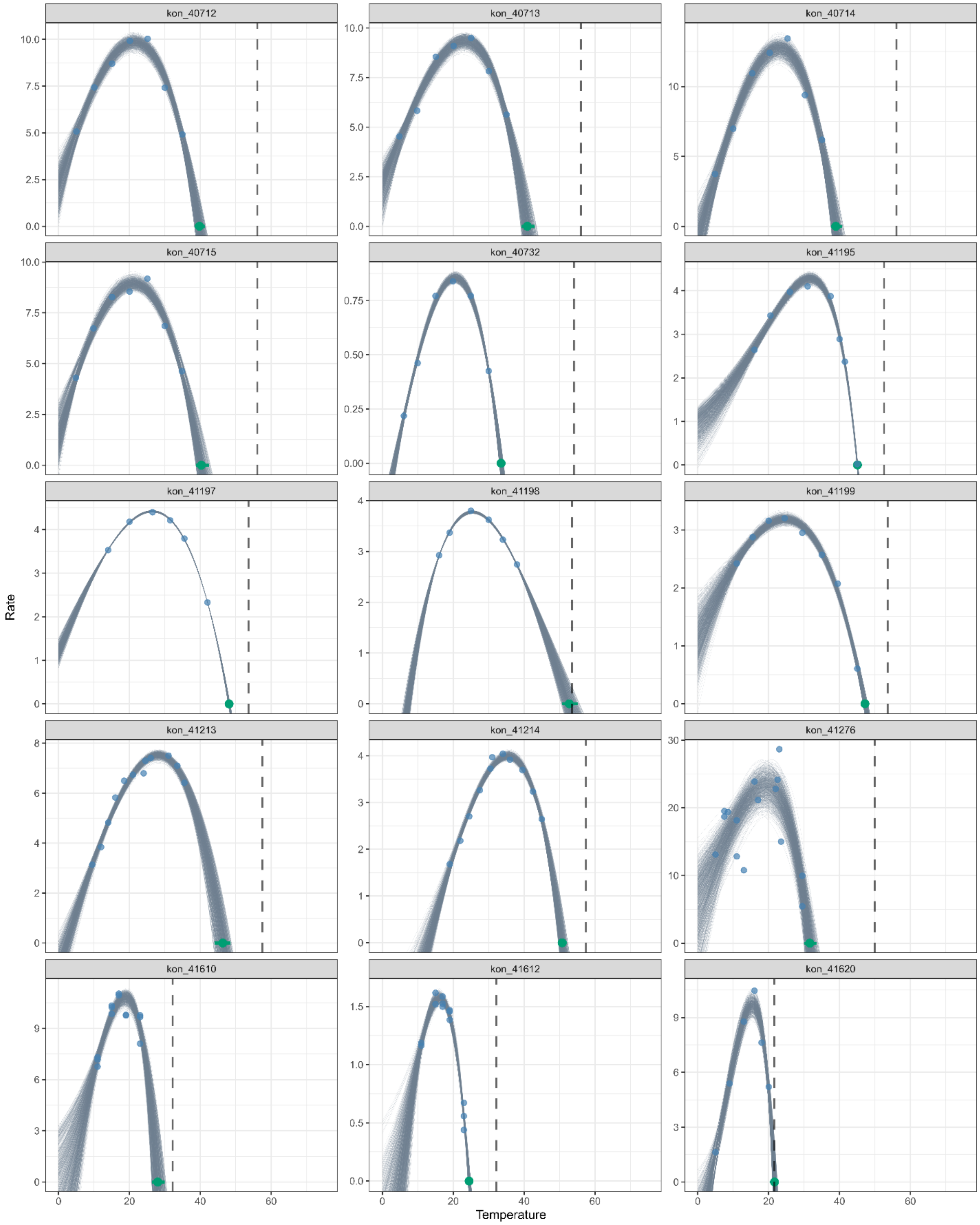
kon_40712
kon_40713
kon_40714
kon_40715
kon_40732
kon_41195
kon_41197
kon_41198
kon_41199
kon_41213
kon_41214
kon_41276
kon_41610
kon_41612
kon_41620
Rate
Temperature

1143

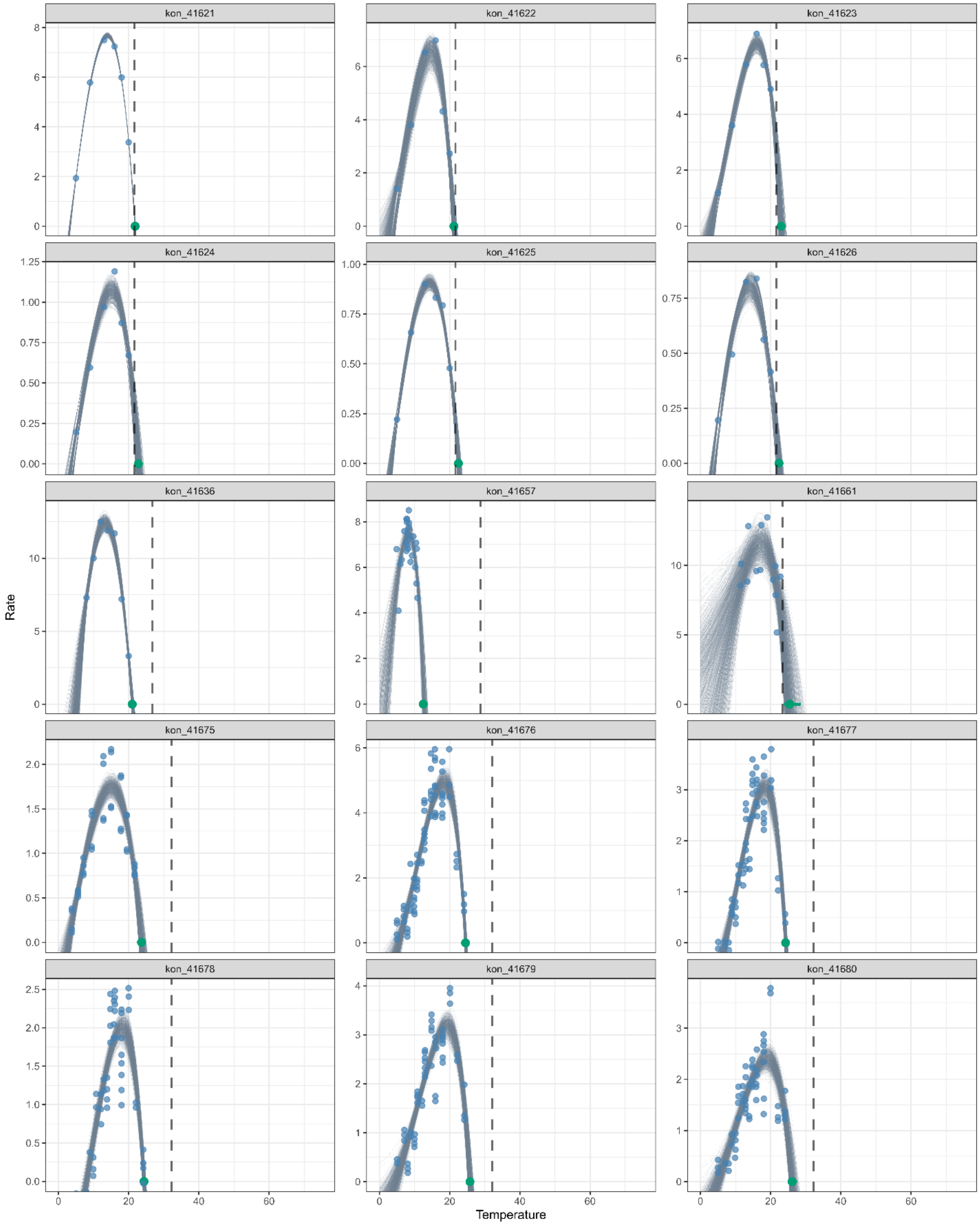

kon_41621
kon_41622
kon_41623
kon_41624
kon_41625
kon_41626
kon_41636
kon_41657
kon_41661
kon_41675
kon_41676
kon_41677
kon_41678
kon_41679
kon_41680
Rate
Temperature

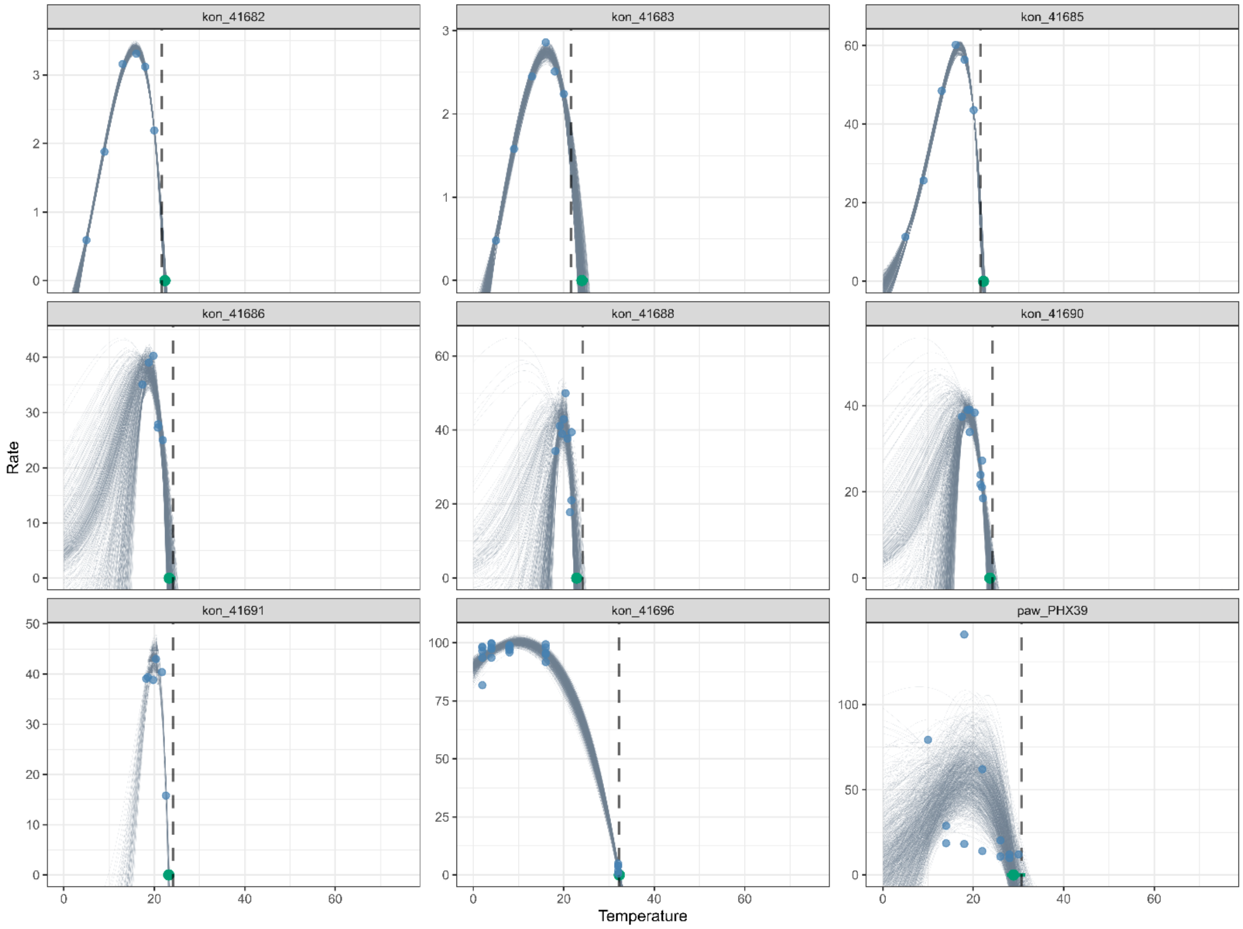
kon_41682
kon_41683
kon_41685
kon_41686
kon_41688
kon_41690
kon_41691
kon_41696
paw_PHX39
Rate
Temperature

References

1. Veritas Health Innovation, Melbourne, Australia., *Covidence systematic review software*.

2. A. Rohatgi, *WebPlotDigitizer*.

23. D. Padfield, G. Matheson, *nls.multstart* (2018).

24. *Bootstrap Methods and their Application* (Cambridge University Press, 1997).
